# Reprogramming adult tendon healing using regenerative neonatal regulatory T cells

**DOI:** 10.1101/2021.05.12.443424

**Authors:** Varun Arvind, Kristen Howell, Alice H. Huang

## Abstract

Tendinopathy is a common clinical problem leading to significant musculoskeletal disability. Using a neonatal mouse model of tendon regeneration compared to adult tendon fibrosis, we identified a unique immune profile in regeneration that is associated with type 2 macrophage polarization and regulatory T cell (Treg) infiltration. Neonatal Treg ablation resulted in a dysregulated immune response leading to failed tenocyte recruitment and loss of functional regeneration. Transcriptional profiling of adult and neonatal tendon Tregs revealed distinct type 1 and type 2 immune signatures that facilitate macrophage polarization following injury. Finally, adoptive transfer of mouse and human neonatal Tregs was sufficient to improve functional regeneration, in contrast to adult Treg transfer. Collectively, these studies uncover a critical role for neonatal Tregs in controlling immune polarization to promote an environment permissive for tendon regeneration. Our findings provide a basis for immune modulating therapies to facilitate regenerative adult tendon healing.

## Introduction

Although tendinopathies and tendon injuries are common and debilitating, treatment options are limited to physical rehabilitation or surgery, which fail to restore native structure or function. In humans, adult tendon rupture results in fibrotic scar formation and impaired function; however, pediatric tendon injuries heal with improved restoration in function and minimal complications^1^. To date, the mechanisms and targets to regenerate tendons remain unknown, due to the paucity of tendon regeneration models. We previously demonstrated functional tendon regeneration in neonatal mice, which was driven by TGFβ-mediate recruitment and differentiation of intrinsic tenocytes^2,3^. Although mitotic potential of tenocytes may be one determinant underlying neonatal regeneration versus adult scar formation, a growing body of evidence also implicates the adult immune environment in poor tendon healing, as excessive inflammation promotes degeneration, fibrosis, and impaired functional outcomes^4–6^. While neonatal immunity has been extensively studied in the context of infection and auto-immunity, the relationship between the unique neonatal immune environment and tissue regeneration is largely unknown. Generally, immunity during the neonatal window is highly tolerant, relatively quiescent, and biased toward a type 2 immune response in response to pathogens^7–9^. Whether this neonatal environment confers a regenerative advantage has not been studied, outside limited contexts.

Although the immune response to wounding is driven by many immune cell types, most wound healing studies focus on macrophages, which are early responders and mediate debris clearance, inflammation, and granulation^10,11^. In regenerative tissues (e.g. neonatal heart, neonatal tendon, adult muscle, adult bone) and regenerative organisms (e.g. salamander, zebrafish), depletion of macrophages results in failed regeneration^12–16^; however macrophages are also indispensable for scar-mediated healing^17,18^. In adult tissues, macrophage polarization toward a Ly-6C^lo^ anti-inflammatory profile appears to promote tissue healing^12,19,20^; however how this polarization is controlled has not been fully defined. There is now increasing evidence that T cells of the adaptive immune system also contribute to regenerative or maladaptive healing. In particular, Foxp3+ regulatory T-cells (Tregs) regulate adult regeneration of muscle, bone, skin, and lung^9,21–23^. The activity of Tregs in non-regenerative adult tissues and whether adaptive immune cells regulate neonatal regeneration is unknown.

## Results

### Neonatal tendon regeneration is defined by rapid polarization from type 1 to type 2 immune response

To determine whether the neonatal immune response to tendon injuries is distinct from the adult response, we profiled immune-related gene expression 3 days after Achilles tendon transection (induced at postnatal day P5 or 4-6 months), using the Nanostring PanCancer Immune Panel. While we initially expected a dampened immune response in neonates compared to adults, neonates showed increased activation of immune genes (127 genes, Fig. 1A). Gene-ontology analysis of differentially expressed genes identified type 1 immune signatures, including production of inflammatory cytokines IL1β and TNF, and angiogenesis in injured adult tendon. In contrast, type 2 immune response and tissue remodeling signatures were identified in injured neonatal tendon (Fig. 1A). Type 2 immune genes included receptors for type 2 cytokines, *Il4ra* and *Il10ra*, which were uniquely upregulated in injured neonatal tendon compared to adult (Fig. 1A). To confirm neonatal signatures, cytokine protein arrays were performed confirming increased abundance of type 2 polarizing cytokines, IL4 and CCL5, in neonatal injured tendon at 3DPI compared to adult (Fig. 1B).

**Figure 1:**
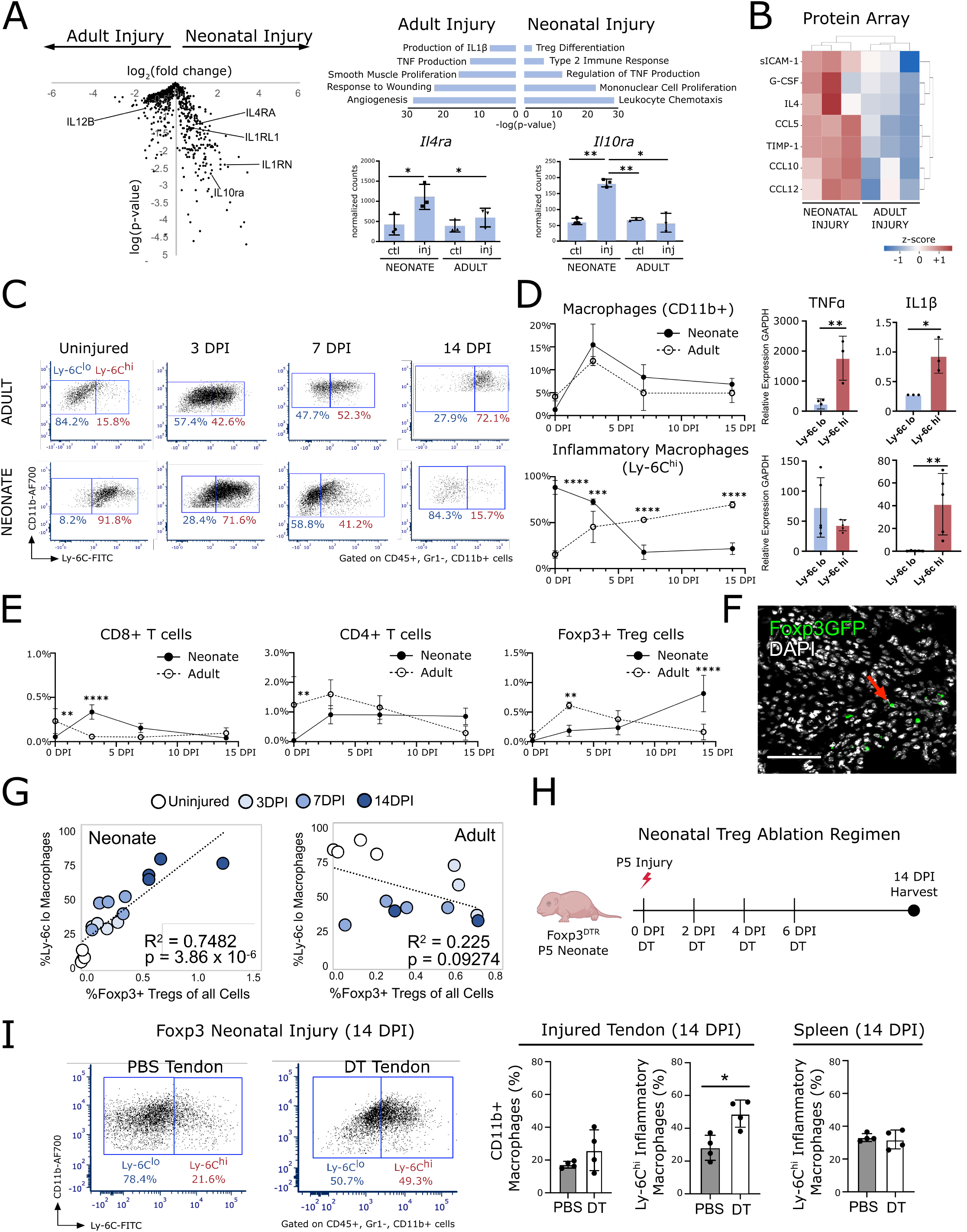
Rapid polarization toward an anti-inflammatory type II immune response in neonates after tendon injury. A) Left: Volcano plot comparing gene expression from NanoString analysis of neonatal vs. adult injured tendon isolated at 3DPI (n=3 mice, p-values adjusted for multiple comparisons). Right: (Top) Gene ontology analysis on differentially expressed genes significantly upregulated (p-adj<0.05) from neonatal injured and adult injured tendon. (Bottom) Expression levels of representative type II immune genes. 1-way ANOVA, Tukey’s posthoc. B) Significantly different representative type II cytokines quantified from cytokine proteome profiling blots of neonatal and adult injured tendon at 3DPI (n=3 mice, 1-way ANOVA, Tukey’s posthoc). C) Flow cytometry profiling of CD11b+ and Ly-6C macrophages in neonatal and adult tendon before and after injury (n=3-7, 2-way ANOVA, Sidak’s posthoc). D) qPCR analysis of inflammatory genes expressed in Ly-6C^lo^ and Ly-6C^hi^ macrophages isolated by FACS from injured neonatal tendon at 14DPI (n=3-4 mice, 2-tailed Student’s t-test). E) Quantification of CD8+, CD4+, and Treg cells (n=3-5, 2-way ANOVA, Sidak’s posthoc). F) Immunofluorescence microscopy of transverse sections through the tendon injury site at 14DPI. Red arrow indicates Foxp3^GFP^ Tregs. Scale, 100µm. G) Pearson’s correlations of Ly-6C^lo^ macrophages and Tregs in tendons after neonatal and adult tendon injury (n=3-4 mice). H) Schematic depicting Treg depletion strategy (created with BioRender.com). (I) Flow cytometry analysis and quantification of macrophages at 14DPI after tendon injury in Foxp3^DTR^ mice treated with PBS or DT (n=4 mice, 2-tailed Student’s t-test). For all quantifications, *p<0.05, **p<0.01, ***p<0.001, ****p<0.0001.

Following adult tendon injury, there is robust recruitment of innate and adaptive immune cells^24^. To define these cell types for neonatal tendon regeneration, we performed flow cytometry and profiled macrophage and T cell subpopulations (Fig. S1A-C). Analysis of the pan-macrophage marker CD11b showed no difference in macrophage recruitment dynamics between neonates and adults. In both neonates and adults, there was a rapid increase in macrophage number to the injured tendon peaking at 3DPI (Fig. 1C). When we distinguished anti-inflammatory and pro-inflammatory macrophages using the marker Ly-6C (CD11b+, Gr1-), we observed polarization toward pro-inflammatory Ly-6C^hi^ macrophages in adults by 14DPI while neonates polarized toward anti-inflammatory Ly-6C^lo^ macrophages (Fig. 1C). Surprisingly, neonatal tendons were enriched for Ly-6C^hi^ macrophages at baseline (and in sham tendons at all timepoints), while adult tendons were enriched for Ly-6C^lo^ macrophages (Fig. 1C, Fig. S2). Importantly, macrophage polarization was specific to the injured tendon as polarization was not observed between neonatal and adult spleen or sham operated tendons, indicating that tendon-specific signals direct polarization (Fig. S2). Analysis of the inflammatory cytokines *Il1β* and *Tnfα* in sorted macrophages showed elevated expression at 3DPI in Ly-6C^hi^ macrophages (Fig. 1D). While *Tnfα* expression levels in Ly-6C^hi^ cells reverted to Ly-6C^lo^ levels at 14DPI, *Il1β* remained dramatically elevated in Ly-6C^hi^ cells (Fig. 1D).

### Neonatal Tregs are required for anti-inflammatory macrophage polarization

Flow cytometry quantification of cytotoxic CD8+ and helper CD4+ T cells revealed no differences between neonates and adults a 14DPI, although differences were observed at baseline and at 3DPI. However, there was increased CD4+, FOXP3+ Tregs in injured neonatal tendon at 14DPI (Fig. 1E). Analysis of transverse cryosections from Foxp3^GFP^ mice at 14DPI further confirmed localization of Foxp3-GFP+ Tregs at the injury site (Fig. 1F). To determine whether neonatal tendon Tregs were peripherally recruited or expanded from tendon-resident Tregs, we inhibited egress of T cells by delivery of FTY720 (daily, 2 weeks) following P5 injury. FTY720 treatment reduced neonatal Treg numbers at 14DPI, indicating peripheral recruitment (Fig. S3).

Treg prevalence was strongly correlated with anti-inflammatory Ly-6C^lo^ macrophage polarization in injured neonatal tendon, but not adult (Fig. 1G), suggesting a potential role for neonatal Tregs in regulating macrophage polarization. To test the requirement for neonatal Tregs in macrophage polarization, neonatal Tregs were selectively ablated using the Foxp3^DTR^ mutant, in which Tregs express the human diphtheria receptor (DTR)^25^. Treg depletion was carried out by delivery of diphtheria toxin (DT) for the first 6 days post-injury (every other day, Fig. 1H)^21,25^, with nearly >90% depletion in the spleen and ∼40% depletion in tendon at 14DPI (Fig. S1D). While total CD11b+ macrophage numbers were unchanged with Treg depletion, neonatal macrophages were polarized toward inflammatory Ly-6C^hi^ macrophages in the injured tendon, with no systemic effects observed in the spleen (Fig. 1I). Collectively, these data show there is a tendon-specific requirement for Tregs in macrophage polarization toward a type 2 response during neonatal regeneration.

### Neonatal Tregs are required for structural and functional tendon regeneration

Having established a role for neonatal Tregs in macrophage polarization following tendon injury, we next tested the requirement for Tregs in neonatal tendon regeneration. Following Achilles tendon transection, impaired healing in Treg-ablated neonates was apparent as early as 7DPI. While normal wound closure was observed in PBS-treated limbs and in sham-operated limbs, wounds remained visible in DT-treated limbs (Fig. 2A). Functional testing using gait analysis and direct tensile testing showed impaired functional recovery of all parameters in Treg-ablated neonates at 28DPI (Fig. 2B, C). To assess structural recovery, we performed collagen hybridizing peptide (CHP) staining to quantify the presence of denatured or damaged collagen^26^. Increased CHP staining was observed in injured tendons of Treg-ablated neonates at 14DPI, indicating persistently damaged collagen structure (Fig. 2D). Collagen quality in the regenerated neotendon was further assessed with polarized light microscopy of Picrosirius Red-stained longitudinal sections to determine collagen alignment ^27,28^. Increased collagen birefringence was observed in PBS-treated neo-tendons compared to Treg-ablated tendons (Fig. 2E). Together, these data revealed a necessary requirement for Tregs in structural and functional neonatal tendon regeneration.

**Figure 2:**
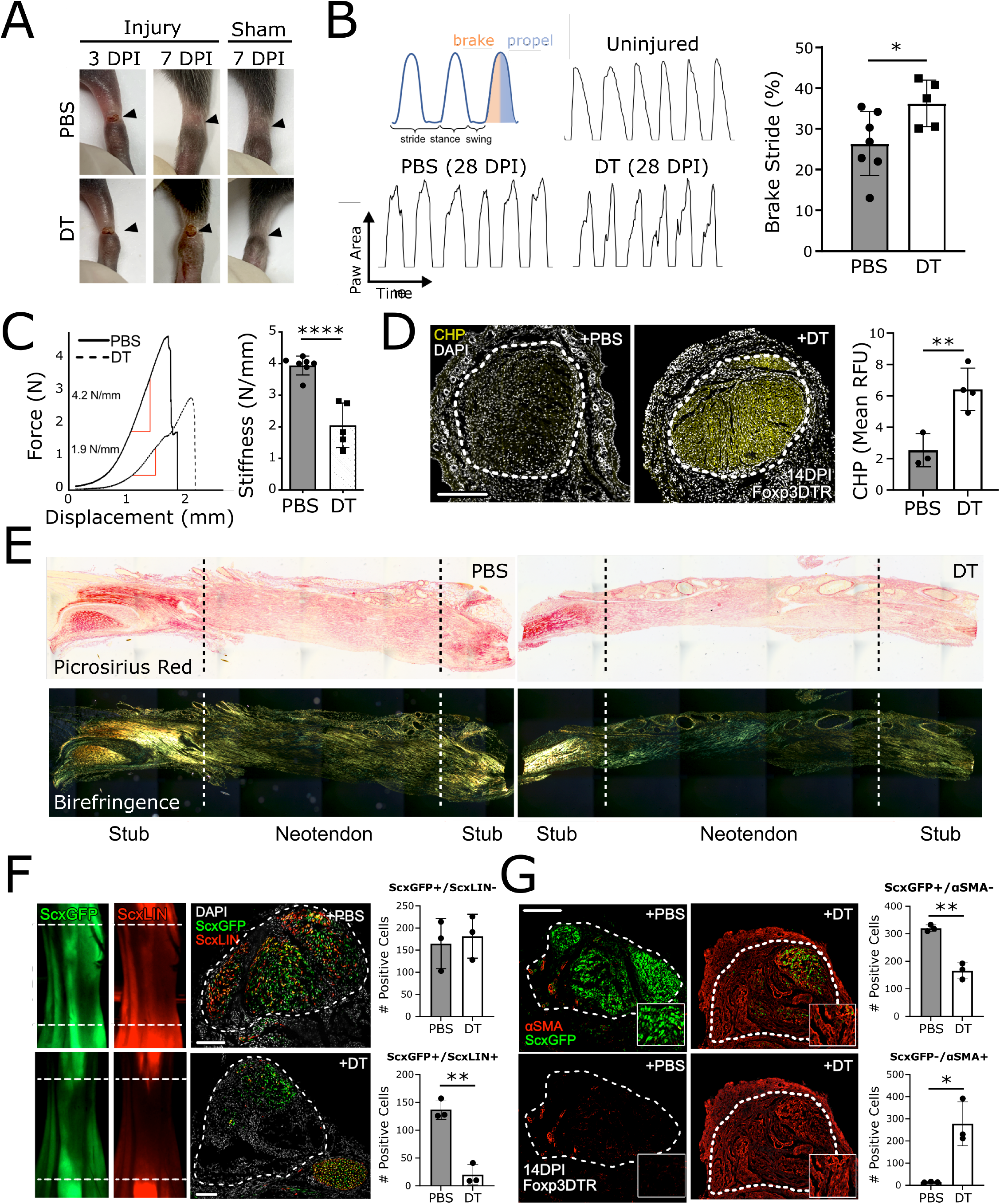
Tregs are required for neonatal tendon regeneration. A) Hindlimb images following Achilles tendon injury or sham surgery in PBS-or DT-treated Foxp3^DTR^ neonates. Black arrowheads indicate site of tendon injury or sham operation. B) Left: Gait schematic and waveform tracings from representative PBS-or DT-treated Foxp3^DTR^ neonatal mice before and after tendon injury. Quantification of gait parameter Brake Stride (n=5-7 mice, 2-tailed Student’s t-test). C) Representative force versus displacement tracings and stiffness quantification from uniaxial mechanical tensile testing of Achilles tendons from PBS-and DT-treated Foxp3^DTR^ neonates at 28DPI (n=5-6 mice, 2-tailed Student’s t-test). D) Fluorescence microscopy and quantification of collagen hybridizing peptide binding in transverse Achilles tendon sections at 14DPI from PBS-or DT-treated Foxp3^DTR^ neonatal mice(n=3-4 mice, 2-tailed Student’s t-test). E) Bright field and polarized light imaging of Picrosirius red stained longitudinal tendon sections from PBS-or DT-treated Foxp3^DTR^ neonatal mice at 14DPI. F) Lineage tracing and quantification of ScxLIN tenocytes visualized by whole mount fluorescence microscopy and transverse Achilles tendon sections from PBS-or DT-treated Foxp3^DTR^; ScxCreERT2; RosaT; ScxGFP neonatal mice at 14DPI. Tamoxifen was administered at P2, P3 (n=3 mice, 2-tailed Student’s t-test). G) Immunofluorescence images and quantification of αSMA+ and ScxGFP+ cells in transverse Achilles tendon sections from PBS-or DT-treated Foxp3^DTR^ neonatal mice at 14DPI (n=3 mice, 2-tailed Student’s t-test). For all quantifications, *p<0.05, **p<0.01, ****p<0.0001. For all scale bars, 100µm.

To identify the cellular basis for impaired regeneration, we next determined recruitment of *Scleraxis-*lineage (ScxLIN) tenocytes since our previous studies implicated these cells in functional tendon regeneration^2,3^. To test whether Tregs are required for tenocyte recruitment, we incorporated the inducible *ScxCreERT2* allele into the *Foxp3*^*DTR*^ background, in combination with the *Rosa26-LSL-tdTomato Ai14; Scx*^*CreERT2*^ reporter and the *ScxGFP* tendon reporter. ScxLIN tenocytes were labeled by tamoxifen administration three days prior to injury (P2, P3) and Treg depletion initiated on the day of injury (P5) for 6 days. At 14DPI, the presence of ScxLIN+, ScxGFP+ tenocytes in the neo-tendon region was greatly reduced with DT treatment compared to PBS control (Fig. 2F). Surprisingly, there was no significant difference in the recruitment of ScxLIN-, ScxGFP+ tenocytes, suggesting that Tregs are required for the recruitment of intrinsically-derived but not extrinsically-derived tenocytes. Analysis of αSMA+ myofibroblasts further revealed persistent αSMA+ cells in Treg-ablated mice at 14DPI, consistent with a non-regenerative, scar-forming phenotype (Fig. 2G).

### Transcriptional profiling by RNA-seq reveals unique anti-inflammatory signatures in neonatal Tregs compared to adult Tregs

Since we observed that deletion of Tregs in neonates resulted in impaired tendon repair similar to adult healing, we next considered whether differences in neonatal and adult Tregs may identify regenerative immune programs. To explore the differences in expression of neonatal and adult Treg immune signatures, we isolated Foxp3^GFP+^ Tregs from injured adult and neonatal tendons at 14DPI and performed RNA-seq. Spleen Tregs isolated from the same mice served as age controls. Principal component analysis (PCA) showed the neonatal tendon Treg (NT) transcriptome was distinct compared to Tregs isolated from neonatal spleen (NS), adult spleen (AS), and adult Tendon (AT) (Fig. 3A). Differential gene expression analysis identified 1,929 significantly upregulated genes (p-adj<0.05) between NT vs. NS, and 513 significantly upregulated genes between AT vs. AS, indicating increased relative transcriptional activity of NTs compared to ATs (Fig. 3B, Fig. S4, Table S1). To determine whether differences in NT and AT signatures were due to differences in Treg activation, we compared differentially expressed genes following subtraction of the canonical Treg activation gene expression signature identified by Hill et al.^29^. Interestingly, differences in NT and AT Treg signatures were not solely attributable to differences associated with an activated Treg signature as 985 genes remained differentially expressed between NT and AT (Fig. 3C)^29,30^. Therefore, we performed comparative analysis of gene expression signatures unique to NT and AT (relative to respective spleen samples to minimize generic stage-specific differences), which revealed a NT-specific transcriptional signature of 1,560 enriched genes (Fig. 3D, Table S2). Interestingly, gene ontology (GO) analysis of the NT gene signature identified GO terms associated with an anti-inflammatory myeloid response, consistent with the anti-inflammatory macrophage polarization observed in neonates (Fig. 3D, 1D). We next performed gene set enrichment analysis (GSEA) on NT and AT DEGs to query immune signatures that distinguish adult and neonatal Tregs. GSEA revealed enrichment for genes associated with “Treg IL4 conversion”^31^ in NT vs. AT, further suggesting that Tregs regulate the rapid initiation of type 2 immunity in neonatal regeneration (Fig. 3E). Hierarchical clustering and GO analysis of interleukin gene signatures (p-adj<0.05) identified terms associated with type 2 and type 1 immune profiles in NT and AT, respectively (Fig. 3F). Collectively, these results show that neonatal tendon Tregs are transcriptionally distinct from adult Tregs and are uniquely capable of polarizing the immune environment toward a type 2 response after injury.

**Figure 3:**
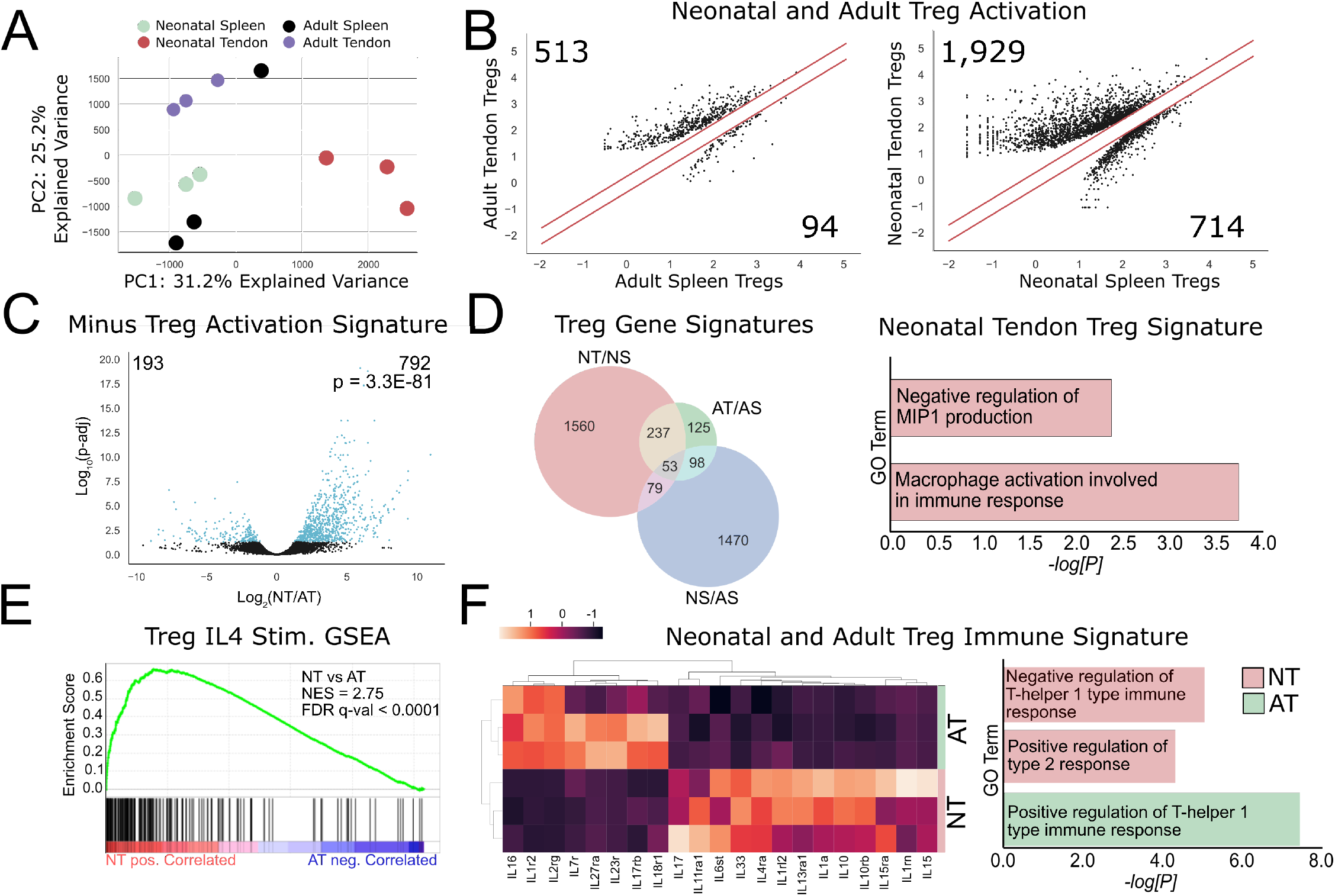
Neonatal and adult tendon Tregs display distinct transcriptional signatures with injury. A) Principal component analysis (PCA) comparing Tregs isolated from neonatal spleen (NS), adult spleen (AS), neonatal tendon (NT), or adult tendon (AT) at 14DPI (n=3 mice). B) Comparison plots of log10-fold gene expression. Numbers indicate the number of upregulated DEGs. Red lines indicate 2-fold difference in expression. C) Volcano plot comparing gene expression of NT versus AT after subtracting a Treg activation gene signature dataset^29^. D) Venn diagram of gene signatures identified by upregulated DEGs. E) Gene set enrichment analysis (GSEA) for genes involved in “Treg IL4 Stimulation” are highly correlated in Tregs isolated from injured neonatal tendon (NT). NES, normalized enrichment score; FDR, false discovery rate. F) Hierarchical clustering and gene ontology analysis of interleukin DEGs between NT and AT, corresponding to Human Genome Organization (HUGO) gene group “Interleukins”.

### Adoptive transfer of neonatal but not adult Tregs rescues functional tendon regeneration

Having identified differences in neonatal and adult Treg transcriptional signatures, we tested whether these differences were functionally meaningful *in vivo*. Neonatal and adult Foxp3^GFP+^ Tregs were isolated from spleens at 3DPI (Fig. S5) and transferred into 5DPI Rag2^-/-^ neonatal mice, which are deficient in T- and B-cells but retain innate immune cells such as macrophages ^32,33^ (Fig. 4A). Analysis of transverse cryosections from injured Rag2^-/-^ hind limbs showed detectable Foxp3^GFP+^ Tregs in the injured tendon at 14DPI, indicating successful Treg engraftment (Fig. 4B). In the absence of adaptive immune cells, we observed impaired immune polarization resulting in a high percentage of pro-inflammatory Ly6c^hi^ macrophages at 14DPI (Fig. 4C). Adoptive transfer of neonatal Tregs led to a marked reduction in Ly6c^hi^ macrophages. While transfer of adult Tregs also improved macrophage polarization relative to control Rag2^-/-^ mice, Ly6c^hi^ cells were still increased compared to neonatal Treg transfer (Fig. 4C). At 28DPI, Rag2^-/-^ neonates showed poor functional recovery compared to wild type (WT) neonates (Fig. 4D). Remarkably, adoptive transfer of neonatal but not adult Tregs was sufficient to rescue gait and tensile stiffness (Fig. 4D). No significant differences were observed between uninjured contralateral hindlimbs (Fig. S6A-C).

**Figure 4:**
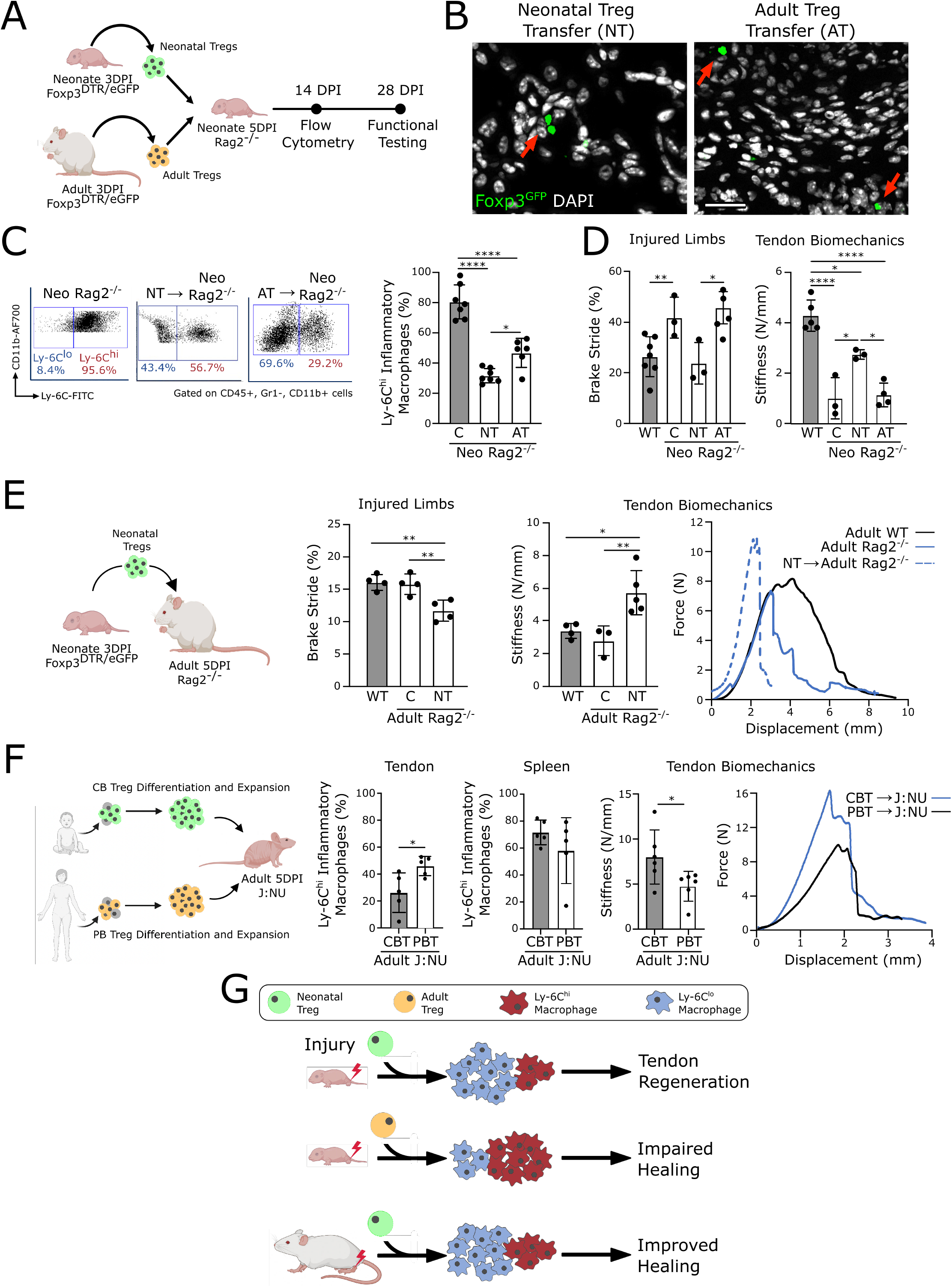
Neonatal Tregs derived from mouse and human are uniquely regenerative and promote tendon healing. A) Experimental design schematic showing adoptive transfer of neonatal and adult mouse Tregs into neonatal immunodeficient Rag2^-/-^ mice. B) Detection of adoptively transferred neonatal and adult Foxp3^GFP^ Tregs (red arrows) in injured Achilles tendon sections at 14DPI. Scale bar: 100µm. C) (Left) Flow cytometry analysis and quantification of Ly-6C macrophages in injured tendons at 14DPI. D) Gait analysis and tensile stiffness quantification of injured tendons at 28DPI (n=6-7 mice, 1-way ANOVA, Tukey’s posthoc). E) Experimental design schematic showing adoptive transfer of neonatal Tregs into adult Rag2^-/-^ mice. Gait analysis and stiffness quantification of adult WT and Rag2^-/-^ mice at 28DPI (n=3-5 mice, 1-way ANOVA, Tukey’s posthoc). F) Experimental design schematic showing derivation of neonatal and adult Tregs from human blood and adoptive transfer into J:NU mice. Quantification of Ly-6C macrophages in injured tendons and spleens of J:NU mice at 14DPI. Right: Quantification of tendon stiffness of injured tendons at 28DPI, with representative force displacement curves (n=5-6 mice, 2-tailed Student’s t-test). G) Summary schematic showing role of neonatal Tregs in immune polarization toward Ly-6C^lo^ macrophages leading to tendon regeneration or improved healing. Panels A, E, F, and G created with BioRender.com. For all quantifications, *p<0.05, **p<0.01, ****p<0.0001.

We next determined whether neonatal Tregs could promote tendon regeneration in an adult environment. In contrast to neonates, there was no difference in functional recovery between WT and Rag2^-/-^ adult mice, suggesting a minimal role for adult adaptive immune cells in adult tendon healing (Fig. 4E). Adoptive transfer of 3DPI neonatal Tregs into 5DPI adult Rag2^-/-^ mice led to significantly improved gait as well as direct tendon tensile stiffness at 28 DPI (Fig. 4E). No differences in gait and tendon tensile stiffness were observed between uninjured tendons (Fig. S6C-D).

To determine whether age-dependent differences in regenerative Tregs are conserved in humans, we then generated neonatal and adult Tregs from human cord blood (CB) and adult peripheral blood (PB), respectively. Since Tregs are a rare population comprising <5% of peripheral CD4+ T-cells in adults and <31% in infants, we pursued a clinically scalable Treg induction and expansion protocol starting from flow-sorted CD4+ T-cells isolated from blood samples^34^. CB-derived T cells exhibit phenotypic characteristics comparable to neonatally-derived lymphocytes with undetectable maternal-fetal T cell contribution^35–38^. Following Treg induction and expansion CB-Tregs and PB-Tregs were adoptively transferred into T cell deficient athymic nude mice at 5DPI (Fig. 4F). Nude mice treated with CB-Tregs showed reduced Ly-6C^hi^ inflammatory macrophages in the neotendon at 14DPI with no significant difference in splenic macrophages, indicating localized regulation at the site of injury (Fig. 4F). At 28DPI, while we observed no difference in gait between CB-Treg and PB-Treg treatment (Fig. S6E), injured mice treated with CB-Tregs had significantly improved tensile stiffness compared to PB-Treg treated mice (Fig. 4F). Taken together, we find that age-dependent differences in Treg-driven immune polarization and regeneration are conserved between mouse and human Tregs, and neonatal Tregs are uniquely capable of promoting functional tendon repair following tendon injury (Fig. 4G).

## Discussion

Analysis of regenerative (axolotl, salamander) and non-regenerative (adult mammals) organisms previously implicated key differences in innate and adaptive immunity underlying regeneration. The observation that regenerative healing in lower-order species is associated with a less sophisticated adaptive immune response led to the hypothesis that adaptive immunity has evolved at the cost of regenerative capacity^39^. Like lower order species, the neonatal immune system is considered immature, which increases susceptibility to infection. One intriguing feature of the neonatal response is bias toward type 2 immunity. While this may predispose infants toward development of allergic reactions, it may also confer regenerative capacity by fostering a permissive immune environment.

Although adaptive immune cells such as T cells are present in neonates, the precise role of individual T cell subtypes (Th1, Th2, and Tregs) in promoting neonatal regeneration remain unclear due to few studies. In Rag2^-/-^ mutants lacking T and B cells, we found that tendon injury resulted in impaired functional regeneration in neonates, but healing was not altered in adults, suggesting a unique requirement for neonatal adaptive immune cells. Using loss of function and adoptive transfer studies, we showed that neonatal tendon Tregs exhibit a unique capacity to polarize tendon macrophages to resolve inflammation and promote functional healing, in both neonatal and adult contexts. While we initially posited that neonates would exhibit a dampened immune response to injury, we observed a robust initial inflammatory response. Adoptive Treg transfer experiments suggested this initial type 1 response is equally important for proper regeneration since Treg transfer at the time of injury resulted in an exacerbated wound that never healed. The rapid transition toward a type 2 response is critical however, since poor healing was consistently observed in cases of sustained type 1 polarization (such as in adult tendon healing, neonatal Rag2^-/-^, or neonatal Treg depletion). Thus a proper balance in type 1 and 2 immune polarization must be achieved.

Since Tregs have long been recognized as potent regulators of inflammation, other studies have investigated whether Tregs promote tissue regeneration^40^. In zebrafish, Tregs are required for organ-specific regenerative programs in the spinal cord, heart, and retina^41^. However, in mammals, the evidence is less clear. Among regeneration-competent adult tissues, (e.g., muscle, lung, skin), release of alarmins following injury stimulates secretion of the EGFR-ligand amphiregulin by Tregs that is critical for tissue repair^19,21,23^. In contrast, adult Tregs are not sufficient to promote regeneration in non-regenerative tissues (e.g., adult heart, brain) despite recruitment and activation^42–44^. While impaired regeneration may be due in part to limited stem/progenitor cell pools, recent studies suggest inflammatory dysregulation also contributes to non-regenerative healing^45,46^. Compared to adult Tregs, neonatal Tregs are known to play a critical role in the maintenance of self-tolerance and have a distinct transcriptional signature from adult Tregs^47^. This may be tissue-specific however; while neonatal cardiac Tregs are required for neonatal cardiac regeneration, transcriptional profiling of regenerative and non-regenerative cardiac Tregs showed surprising similarity and transfer of adult Tregs capably rescued regeneration in the neonatal immunodeficient background^48^. In contrast, not only did we observe decreased abundance of adult tendon Tregs 14 days post-injury, but we also identified dramatic differences in transcriptional signatures and adult transfer was not sufficient for rescue. Interestingly, neonatal tendon Tregs shared DEGs with neonatal cardiac Tregs after injury (not shown), but not with Tregs isolated from adult regenerative tissues such as lung or muscle^19,21,48^. Since we also found that the majority of tendon Tregs after injury are peripherally recruited, these differences suggest that the local injury environment may somehow mediate Treg response and phenotype.

Although our studies showed that neonatal but not adult Tregs rescued tendon regeneration, biomechanical properties were not completely restored, despite restoration of macrophage polarization. Since adoptive transfer studies were carried out in the Rag2^-/-^ background, this may implicate other T or B cells in tendon regeneration. Interestingly, transfer of adult Tregs into the neonatal background also improved macrophage polarization, but no effect on functional healing was observed. These data suggest that there may be an inflammatory threshhold that must be achieved or that neonatal Tregs may have additional functions independent of immune polarization that regulate regeneration. In adult muscle, Tregs are a source of amphiregulin and exogenous application of amphiregulin stimulates satellite cell activation^21^. In the brain, Treg-secreted amphiregulin suppress neurotoxic astrogliosis^43^. With Treg depletion, we found that neonatal tenocyte recruitment was substantially impaired resulting in poor structural and functional recovery. Our previous studies showed a requirement for TGFβ signaling for neonatal tenocyte recruitment after tendon injury^3^. As Tregs are well known to express TGFβ ligands, one function may therefore be to directly activate tenocyte recruitment through TGFβ signaling^49,50^.

Finally, we showed that the regenerative potential of neonatal Tregs may be conserved in humans and that Treg delivery or reprogramming may be a useful approach clinically to improve tendon repair. To our knowledge this is the first study that identifies neonatal Tregs as a regulator for tissue regeneration in both neonatal and adult contexts. We propose that poor healing in adults may be limited, in part, by adult Tregs and that immune modulation is a critical component of tendon regeneration.

## Online Methods

### Peripheral lymphocyte sequestration

Peripheral lymphocyte recruitment was restricted with treatment of FTY720 as per Ito et al.^43^. Briefly, FTY720 (Sigma, Cat: SML0700) was reconstituted to 1 mg/mL and injected intraperitoneally at 25mg/kg body weight every day till the pre-specified endpoint following tendon injury.

### Treg depletion

Treg ablation was carried out in Foxp3^DTR/EGFP^ mice where administration of diphtheria toxin (DT) (Sigma, Cat: 322326) results in apoptosis of Foxp3+ Tregs^51^. DT was administered i.p. at 6 ng/g body weight every other day for 6 days following tendon injury, starting on the day of injury.

### Treg adoptive transfer

For mouse Treg transfer, spleens were harvested 3 days post Achilles tendon injury and processed to prepare a single-cell suspension. Untouched mouse CD4+ T-cells were enriched using magnetic bead isolation as per the manufacturer’s protocol (Thermofisher, Cat: 11416D). Tregs were then isolated via FACS as per above. If insufficient Tregs were collected from a single spleen, Treg populations were pooled from several spleens to achieve sufficient cell quantity. Following isolation, mouse or human Tregs were injected at 5 days post injury via intravenous tail vein injection at 10^3^ cells/ 3g body weight in 150μL or 500μL PBS for neonates and adults, respectively. Murine Tregs were transferred into immunodeficient Rag2^-/-^ mice, while human Tregs were transferred into immunodeficient J:NU mice.

### Human Treg differentiation and expansion

Human neonatal cord blood (Stem Cell Technologies, Cat: 70015) and adult peripheral blood (Stem Cell Technologies, Cat:70026) CD4+ T-cells were differentiated into Tregs using the ImmunoCult Human Treg Differentiation Supplement as per manufacturer’s instructions (Stem Cell Technologies, Cat: 10977). Briefly, Tregs were activated using ImmunoCult human CD3/CD28 T-cell dynabeads (Stem Cell Technologies, Cat: 10971), and cultured in 420IU/mL recombinant human IL-2 (rhIL-2) in ImmunoCult XF-T Cell Expansion Medium (Stem Cell Technologies, Cat: 10981), supplemented with ImmunoCult Human Treg Differentiation Supplement for 14 days. Following Treg induction, human and neonatal Tregs were isolated using the human CD4+, CD127low/-, CD49d-Treg enrichment kit (Stem Cell Technologies, Cat: 19232). Isolated Tregs were expanded for 14 days in ImmunoCult XF-T Cell Expansion Medium supplemented with 500IU/mL rhIL-2 as per manufacturer’s protocol (Stem Cell Technologies, Cat: 10981).

### Mice

ScxGFP mice were used to identify tenocytes as previously described ^2,52^. Foxp3^DTR/EGFP^ (Stock No.: 016958), Rag2^-/-^ (Stock No.: 008449), J:NU (Stock No.: 007850) mice were obtained from Jackson Laboratories. All animal studies were conducted in accordance with Institutional Animal Care and Use Committee (IACUC) at Icahn School of Medicine at Mount Sinai (ISMMS), and all mice were housed in sterile barrier facilities operated by the Center for Comparative Medicine and Surgery (CCMS) at ISMMS. All studies were performed with at least three different mice, with specific sample sizes listed in figure legends. All experiments were performed in both sexes (by random allocation), with all major findings present in both.

Scx^CreERT2^; R26^LSL-tdTomato^ (ScxCET) mice were crossed with Foxp3^DTR/EGFP^ to generate ScxCET; Foxp3^DTR/EGFP^ mice in a C57BL/6 background. Lineage tracing was carried out by tamoxifen gavage at P2 and P3 prior to injury at P5 in neonatal mice. 100mg/mL tamoxifen dissolved in ethanol was dissolved 1:10 in corn oil to achieve 10mg/mL tamoxifen in corn oil. 25uL of 10mg/mL tamoxifen in corn oil was administered by pharyngeal gavage at P2 and P3, prior to injury at P5.

### Mouse tendon injury model

Neonatal (P5) and adult (4-5 month old) mice were anesthetized by hypothermia or isoflurane inhalation, respectively. Achilles tendon injury was achieved by full transection without repair. In mouse hindlimbs, a small incision was made followed by complete tendon transection at the midsubstance. After injury, the skin was closed using Prolene sutures and animals were returned to full cage activity. Buprenorphine was administered intraperitoneally after injury for pain management. Sham injuries were performed in contralateral hindlimbs by skin incision to expose the Achilles tendon, followed by closure with Prolene sutures, without tendon transection.

### FACS isolation and analysis of mouse macrophages and Tregs

#### Tendon

Tendon digestion and single-cell suspension preparation was performed prior to FACS isolation and analysis. At the pre-specified endpoint, tendons were dissected out and incubated in collagenase solution for 4 hours at 37C on a rocker. Collagenase solution was prepared with 5mg/mL Collagenase I (Cat. # LS004196, Worthington Biochemical) and 1mg/mL Collagenase IV (Cat. # LS004188, Worthington Biochemical) in serum free media. Following tendon digestion, tendons were triturated, spin washed (500 x *g* for 5 min at 4C) and resuspended in FACs buffer (2% FBS in 2mM EDT in PBS supplemented with penicillin and streptomycin). Resuspended cells were passed through a 70μm sterile filter to force cell clumps into a single cell suspension. Single cell suspensions were then kept on ice and used for FACS.

#### Spleen

Spleens were harvested and macerated in FCS buffer using a sterile syringe plunger. Spleen suspensions were then passed through a 70μm sterile filter to force cell clumps into a single cell suspension. Splenic single cell suspensions were kept on ice and used for FACS.

#### Macrophage polarization

Cells were stained against CD45 (Biolegend, Cat: 10311, Pe/Cy7, 1:100), CD11b (Biolegend, Cat: 101222, AF700, 1:100), Gr-1 (Biolegend, Cat: 108407, Pe, 108407), Ly-6C (Biolegend, Cat: 128008, FITC, 1:100) for 30mins at 4C in the dark. DAPI was added for live/ dead cell identification where indicated.

#### CD4, CD8, and Treg staining

Cells were stained against CD8 (Biolegend, Cat: 100731, PerCP, 1:100), CD4 (Biolegend, Cat: 116005, Pe, 1:100), CD25 (Biolegend, Cat: 102041, BUV510, 1:100). In mice without Foxp3^DTR/EGFP^, cells were stained against Foxp3 (Biolegend, Cat: 126407, A647, 1:100) using the Foxp3/ Transcription Factor Staining Buffer Kit (eBioscience, Cat: 00-5523-00). Otherwise, Foxp3 was identified using EGFP. DAPI was added for live/ dead cell identification where indicated.

Cells were analyzed and sorted using either a BD LSR II, BD Fortessa, or BD FACSAria III Cell Sorter at the ISMMS Flow Core Facility. Cell sorting was conducted using purity precision mode. FACS profiles were analyzed using FCS Express software.

### RNA isolation, reverse transcription, and qRT-PCR

For RNA isolation from bulk tendon, tendons were dissected out and snap-frozen in liquid nitrogen (3 tendons from 3 mice were combined per sample). Frozen tendons were then pulverized in a Geno/Grinder (SPEX Sample Prep) at 1500 RPM for 30 seconds at room temperature. Following pulverization, RNA isolation was carried out using Trizol/ chloroform extraction. For FACS sorted cells, RNAs were extracted on pelleted cells using Trizol/ chloroform. After Trizol/ chloroform RNA isolation, all RNAs were quantified using NanoDrop2000. Reverse transcription was performed using SuperScript VILO (ThermoFisher, Cat: 11754050) and qRT-PCR performed using SYBR Green PCR Master Mix (ThermoFisher, Cat: 4309155). Mouse primer sequences used are: TNFα (FWD: 5’-TACTGAACTTCGGGGTGATTGGTCC-3’, REV: 5’- CAGCCTTGTCCCTTGAAGAGAACC-3’), IL1β (FWD: 5’-AGTTGACGGACCCCAAAAGAT-3’, REV: 5’-GTTGATGTGCTGCTGCGAGA-3’), and GAPDH (FWD: 5’-TGATGACATCAAGAAGGTGGTGAAG-3’, 5’-TCCTTGGAGGCCATGTAGGCCAT-3’). RNA samples were collected from 3-5 independent mice and ran in triplicate.

### NanoString gene expression and analysis

RNA was isolated from bulk tendons from neonatal and adult mice as described above. Gene expression analysis was performed using the NanoString nCounter platform (NanoString Technologies). Gene counts were normalized using the NanoString nSolver software using background thresholding (threshold count value: 10). Counts were normalized to positive control and housekeeping genes. Normalized gene counts were exported and gene expression was compared with adjusting for multiple comparisons using a Bonferonni correction. Significantly upregulated genes (p-adj < 0.05) were identified and gene ontology analysis was performed using g:Profiler ^53^.

### Cytokine Proteomic Profiling

Protein was isolated from dissected out bulk tendons at prespecified timepoints following injury. Each sample represents 3 tendons combined from 3 separate mice. Following dissection, tendons were snap-frozen in liquid nitrogen and pulverized using a Geno/Grinder (SPEX Sample Prep) at 1500 RPM for 30 seconds at room temperature. Pulverized tendons were resuspended in ice-cold tissue protein extraction reagent (Thermo Fisher, Cat: 78510) supplemented with 1X HALT protease and phosphatase inhibitor cocktail and 1X EDT (ThermoFisher, Cat: 78446) for 5 minutes on ice. After tissue digestion, samples were spun at 10 x *g* for 5 min at 4C, and supernatant was stored at -80C until use. Bradford protein assays were performed, and 150ug of protein was used from each sample for cytokine profiling. Cytokine proteomic profiling was performed as per manufacturer’s instructions (R&D, Cat: ARY006). Densitometry analysis was performed using ImageJ to quantify protein abundance ^54^.

### Histology and immunofluorescence

#### Immunostaining

For immunofluorescence histology, samples were fixed in 4% paraformaldehyde overnight at 4C and then were decalcified in 0.5M EDT, replaced every 3 days, until bones were pliable. Limbs were embedded in OCT and frozen and stored at -80C until use. Alternating transverse cryosections were collected at 12μm thickness, along the length of the tendon to capture the tendon bony insertion to muscle origin. Immunostaining against GFP (Life Technologies, Cat: A11122, 1:200) and α-smooth muscle actin (Sigma, Cat: A5228, 1:100) was performed by incubating overnight at 4C, following cellular permeabilization with 0.1% triton X-100 for 5 minutes at room temperature and blocking for 1 hour. Secondary antibody staining was performed with either Cy3 or Cy5 (Jackson ImmunoResearch), and slides were counterstained with DAPI to visualize nuclei. For quantification, immunofluorescence microscopy was performed on serial transverse sections along the tendon sample. Quantifications were performed across transverse sections spanning the injured portion of the tendon, and were then averaged to achieve an individual result per tendon.

#### Collagen characterization

Denatured or damaged collagen was labeled using a fluorescently conjugated collagen hybridizing peptide as per manufacturer’s instructions (3Helix, Cat: RED60). For picrosirius red staining, limbs were fixed in zinc formalin fixative (z-fix) (Sigma, Cat: Z2902) overnight at 4C. Following fixation, tendons were dissected out, dehydrated, and embedded in methacrylate monomer. 6μm sections were collected and stained with picrosirius red as per manufacturer’s instructions (Abcam, Cat: ab150681). Picosirius red birefringence was imaged using a polarized light filter with the Leica DMB6 to visualize collagen alignment and integrity. All other histology was imaged using a Zeiss Axio Imager Z2 with ApoTome for optical sectioning.

### Whole mount fluorescence imaging

Limbs were first fixed in 4% paraformaldehyde and incubated at 4C overnight. For whole mount fluorescence imaging, skin was dissected away and tendons were visualized using a Leica M165FC stereomicroscope with filters for fluorescence.

### Functional gait analysis

To assess gait, right hindlimb Achilles tendons were injured in mice with the uninjured left contralateral limb serving as a control ^2^. DigiGait Imaging System (Mouse Specifics Inc. Quincy MA) was used to analyze mouse gait following injury. Without pretraining, neonatal and adult mice were gaited at 10cm/s and 50cm/s, respectively. Mice were gaited for 3-5s on a transparent treadmill, with paw contact and position captured using a high speed camera. Mice were gaited in both directions and the average of both results were used for analysis. Footage was analyzed using the DigiGait Analysis Software (DigiGait v12.4). To normalize for age or sex differences, gait parameters (brake) were measured and normalized to stride length to calculate %Brake Stride.

### Biomechanical tendon testing

For biomechanical testing, limbs were harvested and stored at -25C until use. On the day of biomechanical testing, limbs were thawed, and Achilles tendons were dissected out carefully with the bony calcaneal tuberosity. Mechanical tensile testing of mouse Achilles tendons was performed using custom grips to clamp the calcaneal tuberosity and Achilles tendon origin. The tendons were then immersed in a PBS bath at room temperature and preloaded to 0.05N for ∼1min followed by a ramp to failure at 1% strain/second. Force and displacement were recorded using an Instron 8872 Universal Testing System (Instron), which was used to calculate tissue stiffness.

### RNA sequencing and analysis

RNA was isolated using the Arcturus PicoPure RNA Isolation Kit (ThermoFisher, Cat: KIT0204). RNA concentrations were measured with a NanoDrop spectrophotometer (Thermofisher), and quality was assessed with an Agilent TapeStation with RNA integrity number (RIN) > 8 for all samples. RNA amplification, library preparation, and sequencing were performed by Genewiz. Samples were sequenced on the Illumina HiSeq using a 2×150 bp sequencing setting at Genewiz. Sequence reads were trimmed to remove possible adapter sequences and nucleotides with poor quality using Trimmomatic v.0.36 ^55^. The trimmed reads were mapped to the Mus musculus GRCm38 reference genome (ENSEMBLE) using STAR aligner v.2.5.2b ^56^. Unique gene counts were calculated using featureCounts from Subread package v.1.5.2 ^57^. After quantification of gene hit counts, differential gene expression analysis was performed using DESeq2 ^58^. Differentially expressed genes (DEGs) were identified with Benjamini-Hochberg correction for multiple comparisons with a significance of adjusted p-value (p-adj) < 0.05.

Principal component analysis and hierarchical clustering were performed on all DEGs using sci-kit learn v.0.24.1. Treg gene signatures were defined by identifying DEGs with p-adj < 0.05 and log_2_ fold-change > 1 compared to respective populations (e.g., NT vs. NS, AT vs. AS, NS vs. AS). Intersectionality analysis was then performed to identify shared and unique DEGs that comprise distinct Treg signatures. Gene ontology analysis was performed on DEGs using g:Profiler ^53^.

### Statistics

All quantitative data are presented as means ± standard deviation with each point representing independent biological replicates. For comparison of two groups, two-tailed unpaired Student’s t-tests were performed. ANOVA with Tukey’s honestly significant difference (HSD) correction for multiple hypothesis testing was performed for comparison of >2 groups. All statistical analysis was performed using GraphPad Prism v9.0.0.

## Data Availability

All data generated in this study are available from the corresponding author upon reasonable request. RNA-seq data is deposited in Gene Expression Omnibus GSE173770.

## Acknowledgements

We thank the Flow Cytometry Core at Icahn School of Medicine at Mount Sinai for technical assistance. Many thanks to Phillip Nasser and Damien Laudier for assistance with mechanical testing and plastic sectioning. This work was supported by NIH/NIAMS R01 AR069537 and R56 AR076984 to AHH, and NIH/NIAMS F31 AR076905 to VA. VA was also supported by training grant T32 GM007280 from NIH/NIGMS.

## Author Contributions

VA and AH contributed to study conception and experimental design. Experiments were performed by VA and KH. All authors contributed to data analysis and data interpretation. Manuscript preparation was performed by VA and AH. The manuscript was edited and approved by all authors prior to submission.

