## supplemental data for "Reprogramming adult tendon healing using regenerative neonatal regulatory T cells"

**Supplementary Materials**

**Figure S1: Flow cytometry gating strategy and Treg ablation**

A) Flow cytometry gating strategy for identification of Ly-6C$^{lo}$ and Ly-6C$^{hi}$ macrophages. B) Flow cytometry gating strategy for identification of CD4+, Foxp3+ Tregs. C) Viability of flow sorted Tregs (top) and macrophages (bottom). D) Flow plots and quantification demonstrating persistent ablation of Tregs 14DPI following administration of DT or PBS every other day for six days, starting on day of administration in neonatal spleen and tendon (n=3-5, Student's t-test). **p<0.01.

**Figure S2: Macrophage polarization and Treg recruitment is minimal in sham-operated tendons and autologous spleens**

Flow cytometry quantification of immune cells in the contralateral sham-injured tendon or in the spleen after neonatal and adult tendon injury (n=3-7, 2-way ANOVA, Sidak's posthoc). *p<0.05 **p<0.01 ***p<0.001 ****p<0.0001.

**Figure S3: Sequestration of peripheral lymphocytes with FTY720 treatment reduces Treg recruitment to the injured tendon**

Quantification of Treg recruitment to the injured tendon at 14DPI by immunofluorescence microscopy of Foxp3 in transverse sections of injured tendon (n=5, Student's t-test). *p<0.05.

**Figure S4: Neonatal and adult Tregs represent distinct populations**

Volcano plots of DEGs comparing respective Treg populations isolated at 14DPI from injured neonatal tendon (NT), injured adult tendon (AT), autologous neonatal spleen (NS), and autologous neonatal spleen (AS) at 14DPI. Red line corresponds to p-adj = 0.05.

**Figure S5: Treg sorting scheme**

Tregs were sorted by FACS from cells prepared in a single suspension from tendon, spleen, or culture by gating for live, CD4+, Foxp3-EGFP+ Tregs. Tregs were sorted using purity precision mode.

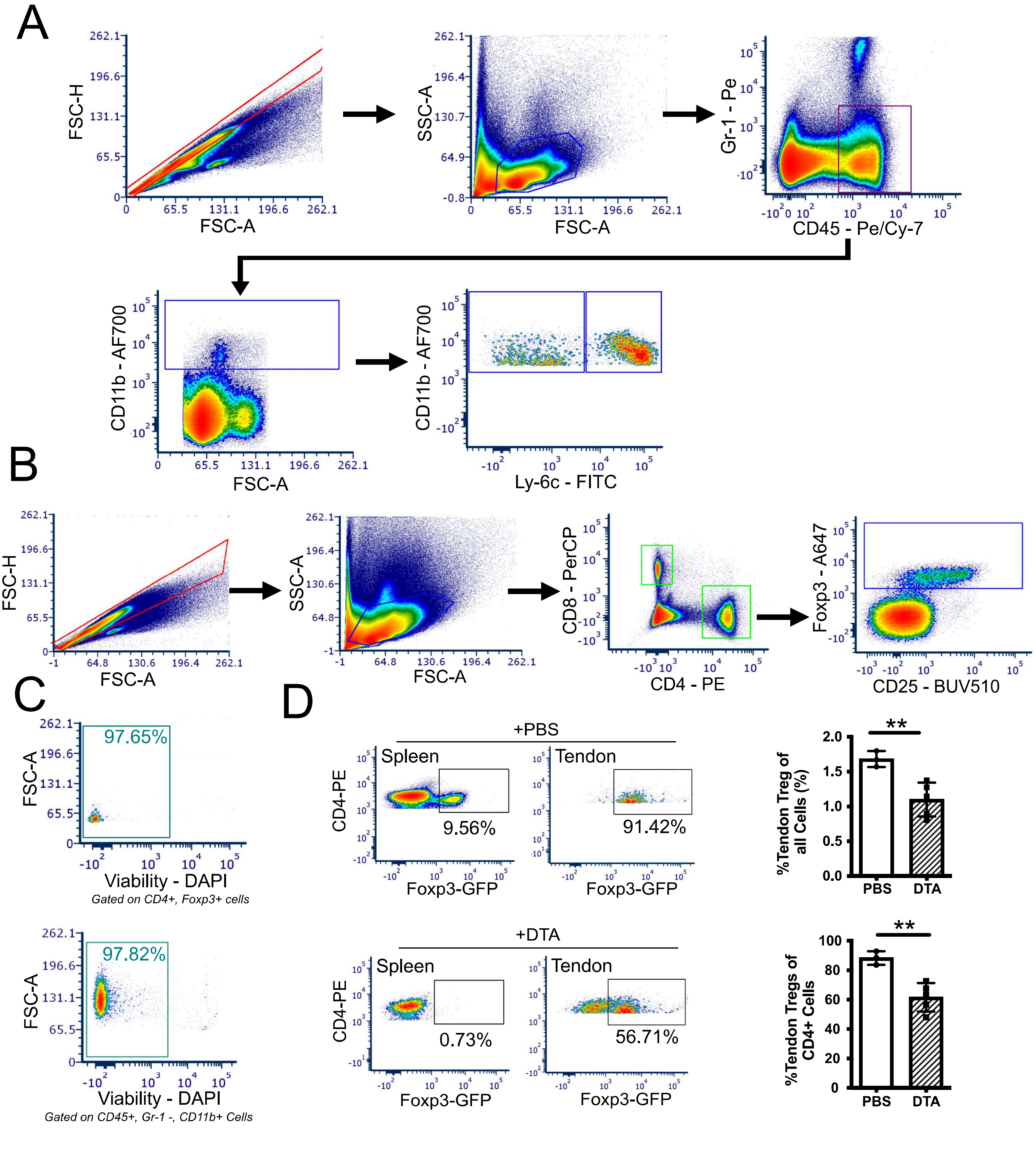

A
FSC-H
FSC-A
SSC-A
FSC-A
Gr-1 - Pe
CD45 - Pe/Cy-7
CD11b - AF700
FSC-A
CD11b - AF700
Ly-6c - FITC
B
FSC-H
FSC-A
SSC-A
FSC-A
CD8 - PerCP
CD4 - PE
Foxp3 - A647
CD25 - BUV510
C
97.65%
FSC-A
Viability - DAPI
Gated on CD4+, Foxp3+ cells
97.82%
FSC-A
Viability - DAPI
Gated on CD45+, Gr-1 -, CD11b+ Cells
D
+PBS
Spleen
Tendon
CD4-PE
9.56%
91.42%
Foxp3-GFP
Foxp3-GFP
+DTA
Spleen
Tendon
CD4-PE
0.73%
56.71%
Foxp3-GFP
Foxp3-GFP
**
%Tendon Treg of all Cells (%)
PBS
DTA
**
%Tendon Tregs of CD4+ Cells
PBS
DTA

Figure S1

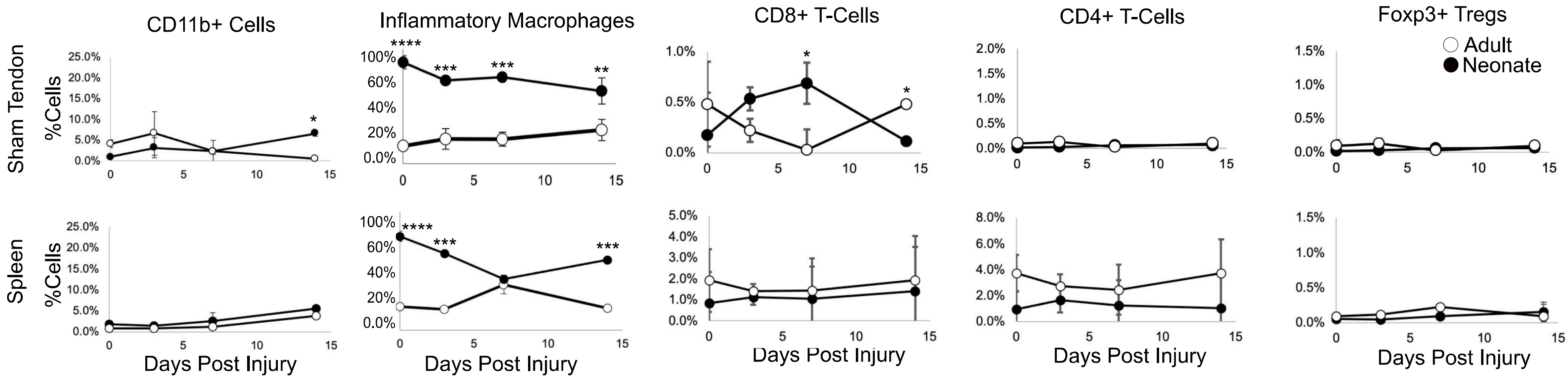
CD11b+ Cells
Inflammatory Macrophages
CD8+ T-Cells
CD4+ T-Cells
Foxp3+ Tregs
Adult
Neonate
Sham Tendon
Spleen
%Cells
Days Post Injury

Figure S2

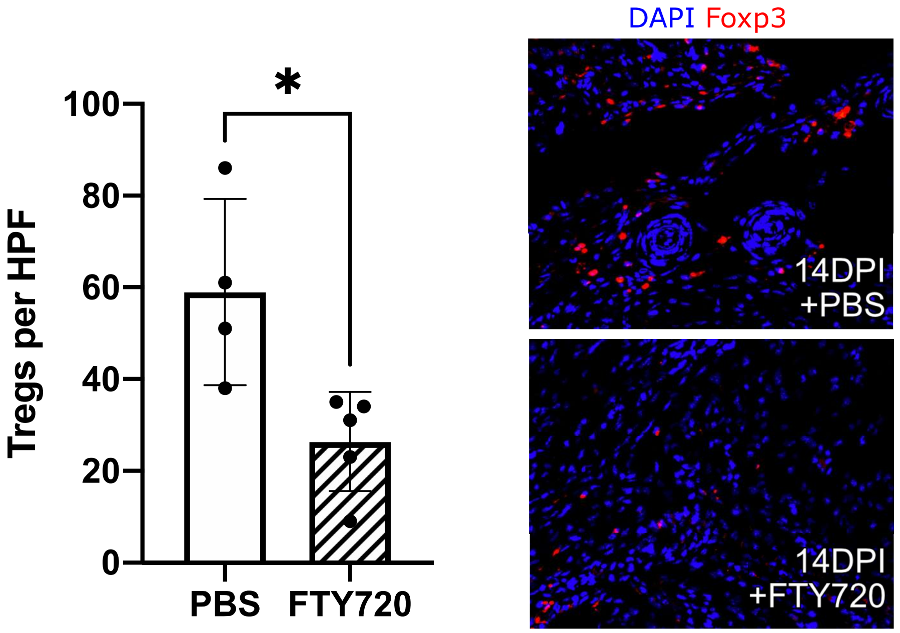
DAPI Foxp3
Tregs per HPF
100
80
60
40
20
0
*
PBS
FTY720
14DPI
+PBS
14DPI
+FTY720

Figure S3

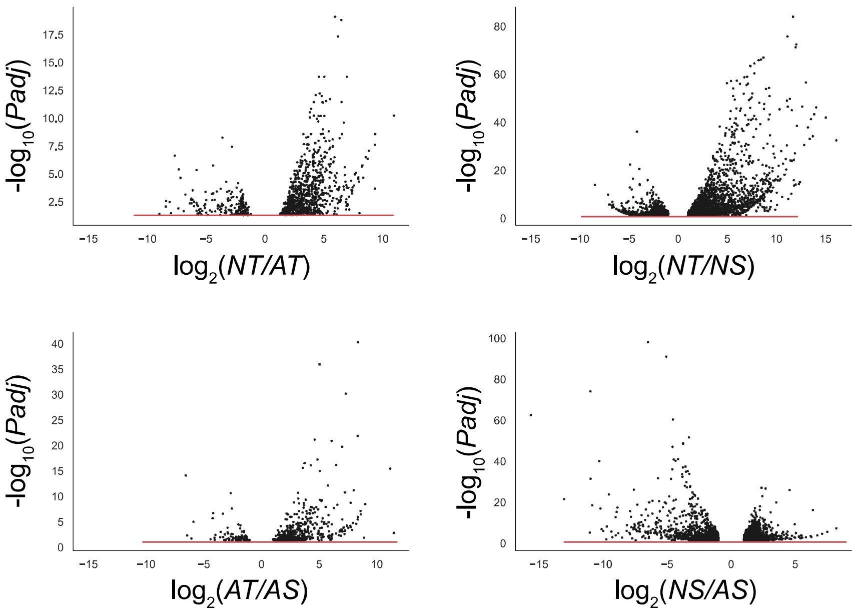
-log10(Padj)
log2(NT/AT)
-log10(Padj)
log2(NT/NS)
-log10(Padj)
log2(AT/AS)
-log10(Padj)
log2(NS/AS)

### Figure S4

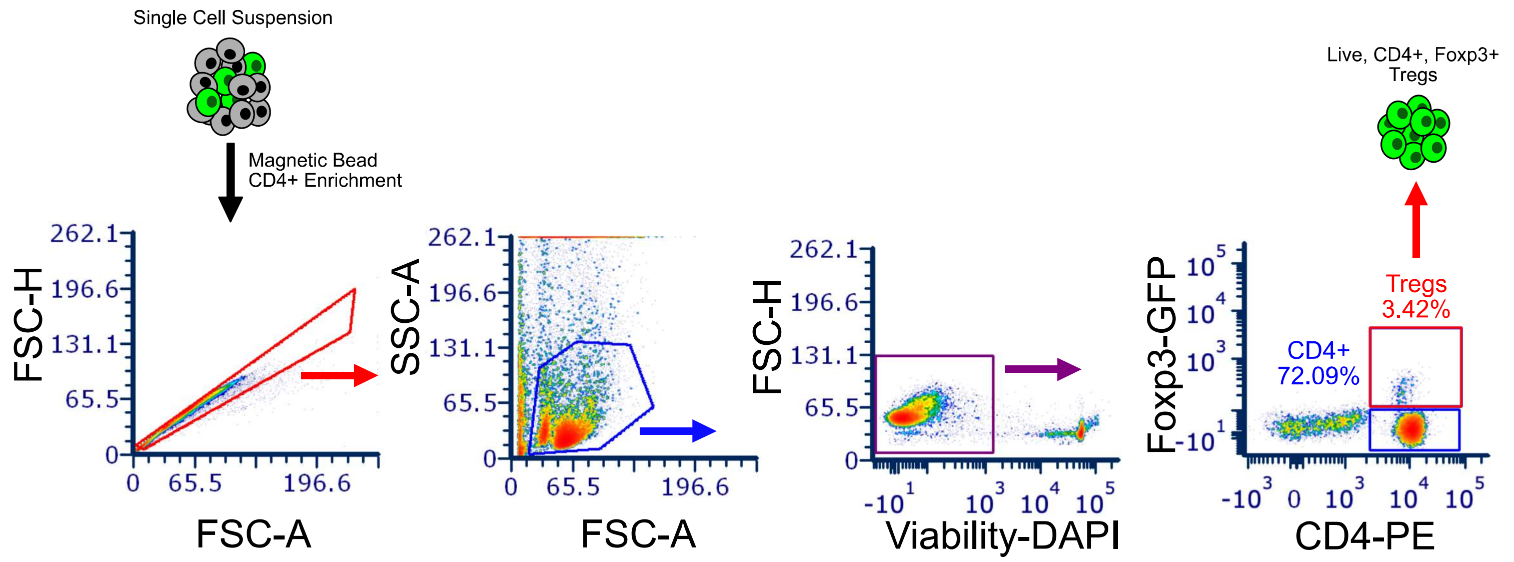

Single Cell Suspension
Magnetic Bead
CD4+ Enrichment
FSC-H
FSC-A
SSC-A
FSC-A
FSC-H
Viability-DAPI
Foxp3-GFP
CD4-PE
CD4+
72.09%
Tregs
3.42%
Live, CD4+, Foxp3+
Tregs

### Figure S5

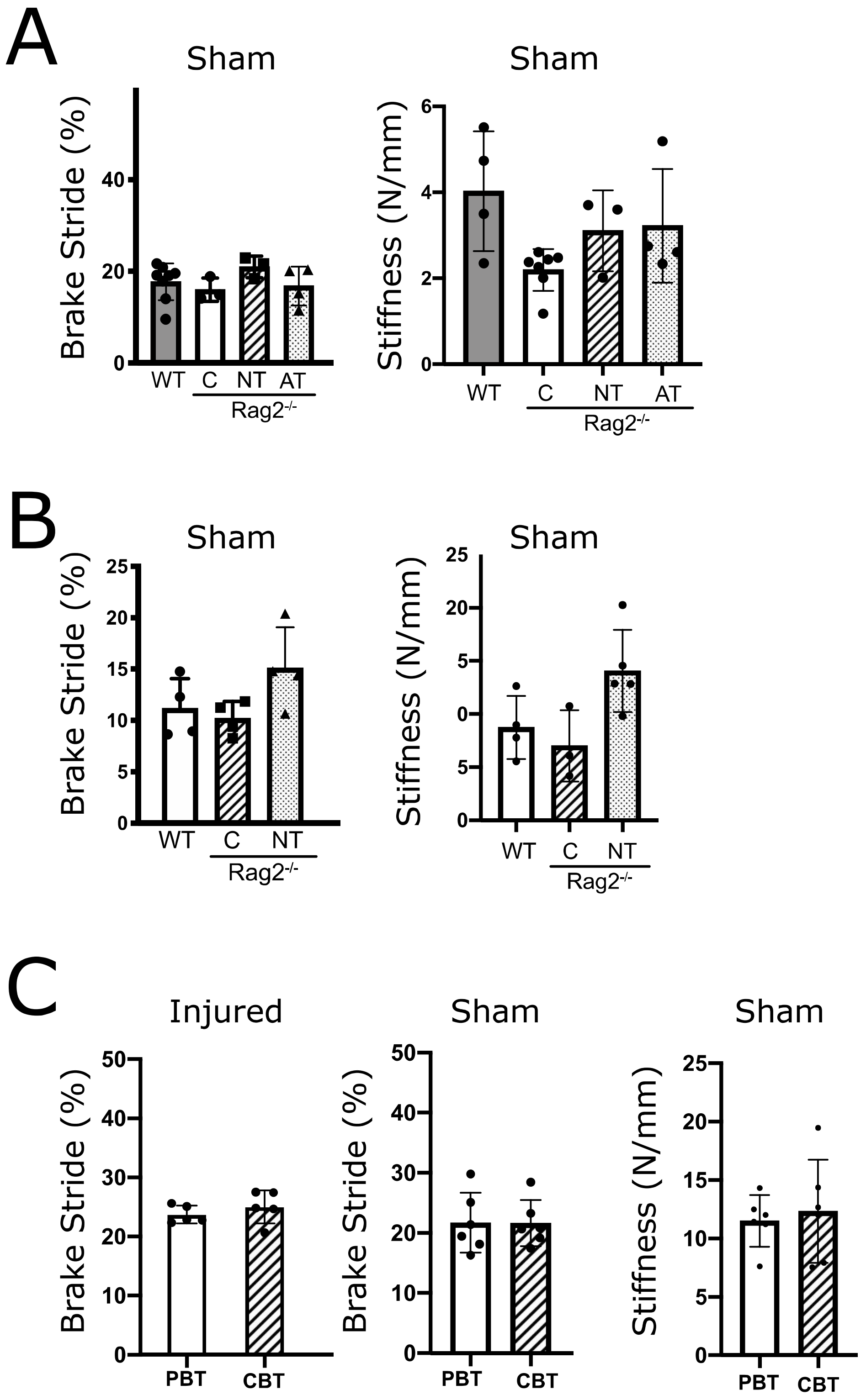
A
Sham
Brake Stride (%)
40
20
0
WT
C
NT
AT
Rag2-/-
Sham
Stiffness (N/mm)
6
4
2
0
WT
C
NT
AT
Rag2-/-
B
Sham
Brake Stride (%)
25
20
15
10
5
0
WT
C
NT
Rag2-/-
Sham
Stiffness (N/mm)
25
20
5
0
5
0
WT
C
NT
Rag2-/-
C
Injured
Brake Stride (%)
50
40
30
20
10
0
PBT
CBT
Sham
Brake Stride (%)
50
40
30
20
10
0
PBT
CBT
Sham
Stiffness (N/mm)
25
20
15
10
5
0
PBT
CBT

Figure S6

**Figure S6: Gait analysis and tendon stiffness from adoptive transfer experiments**

Gait and tendon stiffness quantification from contralateral sham hindlimbs and tendons at 28DPI with adoptive transfer. A) Neonatal Tregs (NT) or adult Tregs (AT) were transferred into neonatal Rag2$^{-/-}$ mice (n=3-5, 1-way ANOVA). B) Neonatal Tregs were transferred into adult Rag2$^{-/-}$ mice (n=4, 1-way ANOVA). C) Human Tregs derived from neonatal cord blood (CB) or adult peripheral blood (PB) were transferred into J:NU mice (n=5, Student's t-test). p>0.05 for all comparisons.

**Table S1:** Top differentially expressed genes from Tregs isolated at 14DPI from injured neonatal and adult tendon or autologous spleen.

| A. Neonatal tendon | | | | | | | |
|---|---|---|---|---|---|---|---|
| | Upregulated | Neonatal tendon vs Neonatal spleen | | | Downregulated | Neonatal tendon vs Neonatal spleen | |
| | **Gene** | Log2(Fold change) | P value | | **Gene** | Log2(Fold change) | P value |
| 1 | Ccl7 | 16.11756401 | 3.24E-33 | 1 | Dtx1 | -8.485450415 | 1.30E-14 |
| 2 | Ccl2 | 15.04297441 | 9.36E-43 | 2 | Klk1 | -7.142399178 | 1.40E-10 |
| 3 | Cxcl1 | 14.05894346 | 5.77E-47 | 3 | Ryr2 | -7.048603714 | 1.72E-06 |
| 4 | Pmp22 | 13.85656522 | 4.30E-44 | 4 | Rasal1 | -6.898174124 | 1.71E-06 |
| 5 | Ccl12 | 13.72105455 | 6.01E-35 | 5 | Olfr164 | -6.727927255 | 3.38E-05 |
| 6 | Ccl8 | 13.54348036 | 4.51E-42 | 6 | Apol7e | -6.704652421 | 7.80E-06 |
| 7 | Ifnb1 | 13.40217462 | 8.33E-34 | 7 | Unc5cl | -6.701333622 | 1.23E-07 |
| 8 | Clec4e | 13.22040876 | 1.19E-38 | 8 | Adam11 | -6.617981038 | 1.87E-06 |
| 9 | Cxcl2 | 13.00907636 | 2.42E-57 | 9 | Iglv3 | -6.550126407 | 2.50E-05 |
| 10 | Gdf15 | 12.83266464 | 2.74E-31 | 10 | Grm8 | -6.30362467 | 0.00013508 |
| 11 | C3ar1 | 12.64222572 | 2.74E-47 | 11 | Upb1 | -6.283211637 | 0.0001212 |
| 12 | Col3a1 | 12.40602799 | 8.29E-16 | 12 | Ap1m2 | -6.111327882 | 0.00028861 |
| 13 | Clec4d | 12.30260428 | 6.05E-15 | 13 | Gm37176 | -6.043332527 | 0.00074513 |
| 14 | Acod1 | 12.0879283 | 3.08E-39 | 14 | Ampd1 | -5.970218159 | 0.00038802 |
| 15 | Spp1 | 12.03273785 | 3.25E-73 | 15 | 1700109H08Rik | -5.811300356 | 0.00190772 |
| 16 | Dab2 | 11.973167 | 5.02E-72 | 16 | Gsta4 | -5.764142953 | 5.12E-07 |
| 17 | Cbr2 | 11.96442607 | 8.44E-15 | 17 | 2610204G07Rik | -5.753509975 | 6.02E-06 |
| 18 | H19 | 11.89183113 | 7.30E-28 | 18 | Gm37387 | -5.663719364 | 0.00045563 |
| 19 | Cxcl3 | 11.81247959 | 2.15E-14 | 19 | Prkag2os1 | -5.652024744 | 0.00315474 |
| 20 | Ptgs2 | 11.77531257 | 1.29E-16 | 20 | Gm26551 | -5.619302549 | 0.00030465 |
| 21 | Ahnak2 | 11.77202851 | 4.52E-25 | 21 | Havcr1 | -5.616943252 | 0.00244117 |
| 22 | Hspa1a | 11.69952041 | 9.70E-85 | 22 | Cacna2d4 | -5.588121211 | 3.09E-05 |
| 23 | Ednrb | 11.57799041 | 8.36E-46 | 23 | Gm42141 | -5.553527057 | 0.00283674 |
| 24 | Il6 | 11.57485 | 1.43E-24 | 24 | Pip5k1b | -5.533350881 | 0.00340617 |
| 25 | Il1rn | 11.52706704 | 5.68E-11 | 25 | Drc1 | -5.494529752 | 0.00340615 |
| 26 | Cyr61 | 11.44411573 | 7.16E-25 | 26 | Zbtb33 | -5.486478046 | 0.00591966 |
| 27 | Gas6 | 11.36501262 | 1.49E-50 | 27 | Bcl2l14 | -5.480163099 | 0.00422257 |
| 28 | Csf3 | 11.21059771 | 9.68E-09 | 28 | Gm12253 | -5.463763217 | 0.00848596 |

| | | | | | | | |
|---|---|---|---|---|---|---|---|
| 29 | Cfh | 11.16563035 | 2.74E-47 | 29 | B3galt5 | -5.444441666 | 0.00623709 |
| 30 | Stab1 | 11.13392526 | 1.67E-76 | 30 | Gm32856 | -5.420240072 | 0.00044821 |
| 31 | Atf3 | 11.08114984 | 8.95E-50 | 31 | Dapl1 | -5.40512046 | 9.91E-05 |
| 32 | Col1a2 | 11.07187683 | 3.67E-31 | 32 | Cd96 | -5.318697753 | 0.00080963 |
| 33 | Dcn | 11.00043169 | 4.48E-21 | 33 | E430014B02Rik | -5.317127646 | 0.00165361 |
| 34 | Ms4a14 | 10.96368788 | 1.94E-25 | 34 | 1700056N10Rik | -5.278750898 | 0.01256707 |
| 35 | Folr2 | 10.84057863 | 6.82E-19 | 35 | Tbxa2r | -5.274714908 | 3.11E-07 |
| 36 | Plau | 10.68372292 | 1.74E-23 | 36 | Dennd3 | -5.186882539 | 0.00287171 |
| 37 | Lyve1 | 10.59651415 | 1.27E-09 | 37 | Car12 | -5.168986055 | 0.00206891 |
| 38 | Cav1 | 10.49769221 | 9.69E-29 | 38 | Wnt3 | -5.143550653 | 0.0156104 |
| 39 | Col6a1 | 10.42415032 | 3.83E-17 | 39 | Gm14085 | -5.123739854 | 1.11E-07 |
| 40 | Pdpn | 10.33497969 | 1.62E-16 | 40 | Trbv12-1 | -5.057194974 | 0.00315413 |
| 41 | Il1a | 10.33250942 | 3.68E-46 | 41 | Nsg2 | -5.055279474 | 6.75E-08 |
| 42 | Ccl4 | 10.23305103 | 1.27E-22 | 42 | Gm11655 | -5.044438941 | 0.00746834 |
| 43 | 8430408G22Rik | 10.15755831 | 3.53E-18 | 43 | Gabrr2 | -5.035635808 | 0.00065772 |
| 44 | Flrt3 | 10.13811396 | 1.55E-16 | 44 | Camk2b | -4.999284837 | 0.02211967 |
| 45 | Tnmd | 10.06990681 | 2.80E-06 | 45 | St8sia1 | -4.968826909 | 5.84E-05 |
| 46 | Col1a1 | 10.0470372 | 2.07E-11 | 46 | Cyp4f17 | -4.951602028 | 0.02528698 |
| 47 | Tcim | 9.939662471 | 5.90E-17 | 47 | Iglc3 | -4.914743508 | 1.38E-13 |
| 48 | Col5a2 | 9.891104683 | 0.000226087 | 48 | Aldh5a1 | -4.907964851 | 0.02938475 |
| 49 | Trem2 | 9.837350247 | 4.21E-26 | 49 | Siglech | -4.90147248 | 3.40E-23 |
| 50 | Creb5 | 9.83661342 | 1.39E-16 | 50 | Gen1 | -4.885714355 | 0.01013772 |
| 51 | Tm4sf19 | 9.824153814 | 1.65E-16 | 51 | Cmya5 | -4.874089911 | 0.03126386 |
| 52 | Fabp4 | 9.72064798 | 1.11E-27 | 52 | Dzip1 | -4.815484336 | 0.00143248 |
| 53 | Gja1 | 9.688496272 | 8.79E-31 | 53 | Igkv9-120 | -4.811748181 | 0.01670075 |
| 54 | Cd14 | 9.68325677 | 1.62E-16 | 54 | Hs3st3b1 | -4.798022348 | 0.04676431 |
| 55 | Adamts1 | 9.664856974 | 1.38E-13 | 55 | Tlr12 | -4.788736606 | 0.04011566 |
| 56 | P2ry2 | 9.628041056 | 1.02E-21 | 56 | Cdh1 | -4.760401164 | 0.00994161 |
| 57 | Colec12 | 9.609077402 | 2.24E-20 | 57 | Gm45718 | -4.750034526 | 0.04286794 |
| 58 | Fbn1 | 9.544775666 | 3.57E-12 | 58 | Ighj2 | -4.734422853 | 0.04420606 |
| 59 | Col12a1 | 9.46560151 | 1.28E-11 | 59 | Tcf7 | -4.733943698 | 8.92E-13 |
| 60 | Rhoj | 9.459294196 | 1.59E-14 | 60 | Gm44901 | -4.716516268 | 0.04703337 |
| 61 | Ifi205 | 9.458293328 | 6.61E-08 | 61 | Akap5 | -4.695962198 | 0.0159006 |
| 62 | Mmp2 | 9.41106256 | 1.99E-15 | 62 | B630019A10Rik | -4.654279427 | 0.00188188 |
| 63 | Lum | 9.403529786 | 2.17E-05 | 63 | Gpr83 | -4.618279215 | 3.15E-17 |
| 64 | Mmp19 | 9.351511736 | 9.94E-14 | 64 | Klc3 | -4.614801181 | 0.00141686 |
| 65 | Rab7b | 9.35140251 | 2.65E-35 | 65 | Gm37642 | -4.606504874 | 0.02216313 |
| 66 | Fmod | 9.340234966 | 4.29E-12 | 66 | Art2b | -4.582675809 | 1.80E-09 |
| 67 | Serpinb8 | 9.328222009 | 3.06E-17 | 67 | Eldr | -4.580713979 | 3.35E-05 |
| 68 | Mcub | 9.319705662 | 5.46E-44 | 68 | Gm42646 | -4.494070769 | 0.00415647 |
| 69 | Fndc1 | 9.311520388 | 0.000160997 | 69 | Gm16575 | -4.492259683 | 0.02192042 |
| 70 | Pf4 | 9.304974869 | 1.16E-51 | 70 | A530030E21Rik | -4.414773008 | 0.03390889 |
| 71 | Igf2 | 9.297426687 | 1.12E-19 | 71 | NA | -4.399950373 | 0.00218986 |
| 72 | Fstl1 | 9.285708086 | 5.64E-22 | 72 | A930024E05Rik | -4.326399067 | 0.0187735 |
| 73 | Gng4 | 9.265471155 | 9.05E-16 | 73 | Cd7 | -4.301532157 | 1.33E-07 |
| 74 | Ccl3 | 9.254655143 | 4.44E-55 | 74 | Kcnip2 | -4.277423217 | 0.01599512 |
| 75 | Arg2 | 9.253196169 | 5.67E-15 | 75 | Vangl2 | -4.252793418 | 0.03850236 |
| 76 | Cd209g | 9.229710944 | 2.42E-12 | 76 | Ighj1 | -4.241871626 | 0.02600944 |

| | | | | | | | |
|---|---|---|---|---|---|---|---|
| 77 | Sparc | 9.211689541 | 5.73E-42 | 77 | Obscn | -4.22209815 | 0.02566959 |
| 78 | Lhfpl2 | 9.166594294 | 9.12E-19 | 78 | Zpbp | -4.20265919 | 0.01230544 |
| 79 | C5ar1 | 9.166550207 | 9.59E-13 | 79 | Sell | -4.199590875 | 7.02E-37 |
| 80 | Pxdn | 9.160479451 | 0.001097873 | 80 | Pard6g | -4.190177678 | 0.00226682 |
| 81 | Nxpe5 | 9.125973118 | 3.23E-13 | 81 | Ddx43 | -4.181389028 | 0.04361187 |
| 82 | Chp2 | 9.115743402 | 2.49E-16 | 82 | Slco4a1 | -4.158820436 | 0.00167288 |
| 83 | Kctd19 | 9.11426844 | 4.69E-12 | 83 | Aff3 | -4.147570633 | 3.58E-07 |
| 84 | Mgp | 9.093019889 | 1.44E-11 | 84 | D630039A03Rik | -4.138126025 | 0.01245085 |
| 85 | C4b | 9.083126853 | 7.26E-32 | 85 | Fgf2 | -4.10826711 | 0.04988967 |
| 86 | Tnfaip6 | 9.080019622 | 4.93E-12 | 86 | Btd | -4.076789916 | 0.00023427 |
| 87 | Col18a1 | 9.075726969 | 7.08E-19 | 87 | Gm37593 | -3.990890271 | 0.02847181 |
| 88 | Thbs4 | 9.068067737 | 0.000597563 | 88 | Ms4a4b | -3.950952267 | 2.84E-21 |
| 89 | Slc7a11 | 9.008413612 | 2.68E-05 | 89 | Ift80 | -3.922953375 | 5.84E-15 |
| 90 | Lpar1 | 8.994118281 | 3.53E-18 | 90 | Gm4956 | -3.912743357 | 1.40E-14 |
| 91 | Edn1 | 8.990135682 | 7.67E-12 | 91 | Kynu | -3.908391305 | 0.04011566 |
| 92 | Nrep | 8.985042201 | 1.66E-16 | 92 | Ehd3 | -3.886480816 | 6.23E-07 |
| 93 | Bgn | 8.98027527 | 8.85E-23 | 93 | Cd8b1 | -3.875076896 | 1.76E-05 |
| 94 | Hspa1b | 8.973364606 | 5.95E-17 | 94 | Apobec2 | -3.87279994 | 0.04568882 |
| 95 | 4930430E12Rik | 8.950899631 | 1.58E-22 | 95 | Fam89a | -3.840349666 | 0.00981464 |
| 96 | Wfdc17 | 8.946011811 | 1.11E-54 | 96 | Degs2 | -3.827877574 | 0.01336971 |
| 97 | Tm4sf1 | 8.917055527 | 6.68E-17 | 97 | Cacna1e | -3.820231608 | 0.01888902 |
| 98 | Reps2 | 8.905190527 | 5.36E-12 | 98 | Gm28942 | -3.816048787 | 0.00883299 |
| 99 | Car13 | 8.874317655 | 3.82E-15 | 99 | Klf16 | -3.809614394 | 0.04262613 |
| 100 | C1qtnf3 | 8.818029701 | 0.000806455 | 100 | Samd10 | -3.805228901 | 1.81E-05 |
| 101 | Mfap5 | 8.813384537 | 1.39E-09 | 101 | Sidt1 | -3.778480951 | 6.12E-07 |
| 102 | Hspg2 | 8.802896402 | 0.000118 | 102 | Smim5 | -3.76397878 | 0.00662149 |
| 103 | Slc13a3 | 8.787533805 | 2.09E-19 | 103 | Rundc3a | -3.74650178 | 0.01924224 |
| 104 | Thbs1 | 8.76828235 | 9.24E-17 | 104 | Btbd11 | -3.746319419 | 0.01022039 |
| 105 | Cd209f | 8.765534232 | 2.46E-05 | 105 | Gins3 | -3.732202335 | 0.00111262 |
| 106 | Hspb1 | 8.763945706 | 7.16E-08 | 106 | Ctsw | -3.725522017 | 6.53E-09 |
| 107 | Cxcl16 | 8.754986654 | 2.40E-48 | 107 | Klk1b27 | -3.721801385 | 0.02283831 |
| 108 | Igfbp5 | 8.748992892 | 5.26E-11 | 108 | H2-Ob | -3.706344057 | 4.50E-07 |
| 109 | Rab34 | 8.747495063 | 1.80E-10 | 109 | Ccr9 | -3.6391057 | 6.97E-08 |
| 110 | Ccl9 | 8.746967269 | 2.63E-46 | 110 | Hsh2d | -3.63118836 | 7.17E-05 |
| 111 | Aplnr | 8.739028758 | 1.65E-10 | 111 | AW146154 | -3.622893644 | 0.00165688 |
| 112 | Gm15726 | 8.714598808 | 8.81E-16 | 112 | Ankrd13b | -3.591977856 | 0.03852266 |
| 113 | Bicc1 | 8.701053434 | 6.72E-09 | 113 | Actg2 | -3.589495502 | 4.46E-05 |
| 114 | Prune2 | 8.688173394 | 5.26E-30 | 114 | Trbj2-1 | -3.553778272 | 0.03079132 |
| 115 | Oasl1 | 8.686652797 | 9.71E-68 | 115 | Serpini1 | -3.53804316 | 0.02216313 |
| 116 | Ch25h | 8.674061022 | 2.30E-10 | 116 | Ptprf | -3.531317425 | 0.00423098 |
| 117 | Tnfsf15 | 8.660815618 | 6.09E-16 | 117 | Gm16973 | -3.517025469 | 0.03523768 |
| 118 | Olfml2a | 8.625612159 | 1.26E-08 | 118 | Card6 | -3.49335076 | 1.13E-06 |
| 119 | Fabp3 | 8.624267625 | 5.69E-15 | 119 | Invs | -3.395234586 | 0.01342918 |
| 120 | Gm42793 | 8.624104436 | 1.27E-10 | 120 | Rhebl1 | -3.392027055 | 0.01798246 |
| 121 | Il15 | 8.591137204 | 5.53E-23 | 121 | Gm27252 | -3.376963551 | 0.04935232 |
| 122 | Clec5a | 8.586503371 | 1.90E-05 | 122 | Pglyrp2 | -3.364905479 | 8.34E-05 |

| | | | |
|---|---|---|---|
| 123 | Tmem171 | 8.581584042 | 9.81E-12 |
| 124 | Jam2 | 8.558158421 | 3.69E-13 |
| 125 | Alox5 | 8.534472455 | 4.18E-17 |
| 126 | Slc7a2 | 8.527243027 | 1.19E-09 |
| 127 | Cfb | 8.523997628 | 8.38E-17 |
| 128 | Gm48065 | 8.488491716 | 3.35E-09 |
| 129 | Gm38158 | 8.46639819 | 6.41E-10 |
| 130 | Cxcl10 | 8.434524994 | 7.06E-67 |
| 131 | Mir99ahg | 8.415808184 | 2.14E-12 |
| 132 | Sparcl1 | 8.392755359 | 1.57E-16 |
| 133 | Csf2 | 8.387559079 | 8.69E-09 |
| 134 | Postn | 8.38685843 | 3.71E-06 |
| 135 | Cracr2b | 8.378836799 | 1.21E-09 |
| 136 | Fcrls | 8.37085458 | 5.93E-40 |
| 137 | Atp13a4 | 8.369627178 | 1.50E-09 |
| 138 | Gm13391 | 8.366841057 | 7.49E-09 |
| 139 | Car3 | 8.344486096 | 5.72E-08 |
| 140 | Col4a1 | 8.326925806 | 7.40E-07 |
| 141 | Rassf8 | 8.314603557 | 6.78E-09 |
| 142 | Efcab6 | 8.278675765 | 4.51E-08 |
| 143 | Ms4a7 | 8.265358233 | 6.46E-49 |
| 144 | Cdkn1a | 8.196074843 | 1.23E-66 |
| 145 | Cxcl11 | 8.186972184 | 2.03E-08 |
| 146 | Col4a2 | 8.185357734 | 4.62E-25 |
| 147 | Sh3pxd2b | 8.148878226 | 2.68E-05 |
| 148 | Serpine1 | 8.138713416 | 9.13E-08 |
| 149 | Gpnmb | 8.13575179 | 3.63E-14 |
| 150 | Adgrf5 | 8.13572828 | 5.24E-08 |
| 151 | Meox2 | 8.129598388 | 1.67E-08 |
| 152 | C1qtnf1 | 8.123781046 | 1.52E-08 |
| 153 | Tlr8 | 8.121243531 | 1.27E-12 |
| 154 | Arpin | 8.103316402 | 2.40E-08 |
| 155 | Npl | 8.093011968 | 2.71E-07 |
| 156 | Myof | 8.076890501 | 2.92E-12 |
| 157 | Gstm2 | 8.06201375 | 6.70E-08 |
| 158 | Psd3 | 8.049015334 | 4.39E-13 |
| 159 | Gm14636 | 8.037213879 | 1.97E-15 |
| 160 | Mfsd7a | 8.006081016 | 1.81E-07 |
| 161 | Il1b | 7.999541099 | 5.59E-10 |
| 162 | Tmem119 | 7.9874294 | 2.13E-10 |
| 163 | Inhba | 7.987178347 | 1.20E-18 |
| 164 | Mmp13 | 7.984908542 | 2.28E-08 |
| 165 | Ace | 7.969652163 | 0.001098629 |
| 166 | Cxcl12 | 7.962761995 | 0.000645354 |
| 167 | Eps8 | 7.945734691 | 5.72E-38 |
| 168 | Ifi204 | 7.936155068 | 5.03E-42 |

| | | | |
|---|---|---|---|
| 123 | Zfp472 | -3.352859757 | 0.0051949 |
| 124 | Rpgrip1 | -3.328040937 | 0.00013391 |
| 125 | Ppfia4 | -3.32323897 | 0.00264397 |
| 126 | Osbpl5 | -3.312062781 | 0.00951935 |
| 127 | Gm37274 | -3.307975862 | 0.04286794 |
| 128 | Bicdl1 | -3.276822393 | 0.00318702 |
| 129 | Clec2i | -3.243174556 | 3.68E-06 |
| 130 | Spib | -3.234462397 | 1.71E-05 |
| 131 | Trbv13-3 | -3.225853868 | 0.00356659 |
| 132 | Blk | -3.224378554 | 0.04225086 |
| 133 | Xrcc5 | -3.213122129 | 6.22E-05 |
| 134 | Mettl15 | -3.204208081 | 0.02644264 |
| 135 | Xxylt1 | -3.191043961 | 0.00019288 |
| 136 | Trbv17 | -3.18606856 | 0.0051567 |
| 137 | Trbv13-1 | -3.182452057 | 0.00240737 |
| 138 | Bcl2 | -3.164705431 | 1.03E-10 |
| 139 | Aven | -3.136452089 | 0.00031301 |
| 140 | Trib2 | -3.134135208 | 2.55E-11 |
| 141 | Mettl27 | -3.132653917 | 0.03577153 |
| 142 | Gimap4 | -3.108974684 | 1.15E-16 |
| 143 | Gm30211 | -3.083442148 | 0.03791325 |
| 144 | Trbj1-5 | -3.079740784 | 0.02385426 |
| 145 | Pars2 | -3.073176442 | 0.03840837 |
| 146 | 6720464F23Rik | -3.062961014 | 0.00819684 |
| 147 | Foxk1 | -3.059493195 | 0.00886526 |
| 148 | Tet1 | -3.047863742 | 0.00337992 |
| 149 | C530050E15Rik | -3.046905937 | 0.00255717 |
| 150 | Cd160 | -3.037447639 | 0.00333703 |
| 151 | Klhl42 | -2.956767408 | 0.00877044 |
| 152 | Gm39323 | -2.95085742 | 0.00296138 |
| 153 | Rap1gap2 | -2.943627561 | 0.0163122 |
| 154 | Bdh1 | -2.930224271 | 0.00596265 |
| 155 | Trbv13-2 | -2.927334941 | 1.53E-05 |
| 156 | Reck | -2.926763503 | 0.04206911 |
| 157 | Tango6 | -2.924174908 | 0.04214118 |
| 158 | F2rl1 | -2.922630596 | 0.00575058 |
| 159 | Gm15708 | -2.916436322 | 0.01785134 |
| 160 | Bora | -2.894682683 | 0.00184456 |
| 161 | Gpr18 | -2.88805044 | 6.99E-06 |
| 162 | Harbi1 | -2.876796491 | 0.02339093 |
| 163 | Fcrl1 | -2.876789413 | 0.02324611 |
| 164 | Prkca | -2.863393984 | 2.26E-07 |
| 165 | Rbm38 | -2.859076347 | 3.40E-08 |
| 166 | Txk | -2.855039938 | 4.40E-06 |
| 167 | Clint1 | -2.847860643 | 2.09E-11 |
| 168 | Hgh1 | -2.835658333 | 0.02779492 |

| | | | | | | | |
|---|---|---|---|---|---|---|---|
| 169 | Exoc3l4 | 7.929882273 | 1.29E-07 | 169 | Trbv26 | -2.814916054 | 0.01471402 |
| 170 | Oas1g | 7.922964028 | 1.00E-11 | 170 | Rfx7 | -2.790485269 | 1.82E-05 |
| 171 | Htra3 | 7.879000432 | 1.39E-07 | 171 | Scml4 | -2.790393312 | 3.68E-06 |
| 172 | Ctf2 | 7.865820543 | 2.37E-07 | 172 | Zmym1 | -2.782665016 | 0.00820611 |
| 173 | Nlrp1c-ps | 7.852094641 | 5.42E-07 | 173 | Piga | -2.770251471 | 0.01519413 |
| 174 | Il7 | 7.846870123 | 3.02E-07 | 174 | Tgtp1 | -2.769135451 | 0.00044821 |
| 175 | Thbd | 7.844081563 | 2.14E-14 | 175 | Gm37529 | -2.763695689 | 0.03218191 |
| 176 | Serping1 | 7.807064582 | 3.35E-07 | 176 | Trbv1 | -2.75511838 | 0.00165688 |
| 177 | Cmklr1 | 7.799689855 | 2.97E-17 | 177 | Rwdd2b | -2.753531307 | 0.0298314 |
| 178 | A4galt | 7.790288983 | 2.68E-07 | 178 | Trbv5 | -2.727187192 | 0.00642081 |
| 179 | Mrc1 | 7.769648115 | 3.67E-61 | 179 | Jaml | -2.726208908 | 0.00260011 |
| 180 | Cd63 | 7.761907328 | 2.66E-65 | 180 | Nipal1 | -2.720125276 | 0.01314724 |
| 181 | Msr1 | 7.756134436 | 5.73E-09 | 181 | Pitpnm2 | -2.718914737 | 0.00019483 |
| 182 | Srxn1 | 7.746061518 | 2.08E-43 | 182 | Pcgf6 | -2.718431445 | 0.01517506 |
| 183 | Capn6 | 7.728208887 | 4.90E-07 | 183 | Abi3 | -2.712947296 | 0.0093028 |
| 184 | Mfap4 | 7.715271741 | 1.12E-06 | 184 | Zfp287 | -2.711079995 | 0.00466877 |
| 185 | Nfib | 7.707896543 | 9.15E-10 | 185 | Tesc | -2.705173987 | 0.00036848 |
| 186 | S100a16 | 7.686291281 | 1.31E-11 | 186 | Lef1 | -2.705162456 | 1.91E-12 |
| 187 | Tspan7 | 7.684710964 | 5.74E-08 | 187 | Lta | -2.69742592 | 0.00038183 |
| 188 | Gm14461 | 7.682914011 | 1.84E-06 | 188 | 2610035D17Rik | -2.696472903 | 0.00118219 |
| 189 | Mid1-ps1 | 7.680186326 | 3.36E-06 | 189 | Cd226 | -2.69441282 | 0.00086419 |
| 190 | Copz2 | 7.679603276 | 1.79E-11 | 190 | Prkcq | -2.68235213 | 5.40E-10 |
| 191 | Mt2 | 7.6756513 | 1.42E-34 | 191 | S1pr4 | -2.682091596 | 6.45E-09 |
| 192 | Olr1 | 7.674116841 | 2.80E-13 | 192 | Galnt12 | -2.679415444 | 0.00029309 |
| 193 | Wwc1 | 7.669935755 | 2.77E-06 | 193 | Toe1 | -2.673966146 | 0.03094493 |
| 194 | Pgf | 7.642809122 | 1.25E-11 | 194 | Cd300c | -2.673281438 | 0.01010743 |
| 195 | Plvap | 7.635490839 | 1.32E-17 | 195 | Trbv31 | -2.645175901 | 0.00433417 |
| 196 | Mamdc2 | 7.629007346 | 3.61E-06 | 196 | Gimap1os | -2.644106867 | 0.00023425 |
| 197 | Ifi202b | 7.628162889 | 2.08E-06 | 197 | Btla | -2.641219062 | 3.19E-05 |
| 198 | Siglec1 | 7.627864632 | 1.76E-24 | 198 | Miga1 | -2.631663278 | 0.02878657 |
| 199 | Uaca | 7.626769189 | 7.15E-11 | 199 | Mgst2 | -2.605310493 | 3.47E-05 |
| 200 | Gm44386 | 7.622870248 | 2.60E-06 | 200 | Abhd8 | -2.599195309 | 0.00017859 |

**B. Adult tendon**

| | Upregulated<br>**Gene** | **Adult tendon vs Adult spleen**<br>Log2(Fold change) | P value | | Downregulated<br>**Gene** | **Adult tendon vs Adult spleen**<br>Log2(Fold change) | P value |
|---|---|---|---|---|---|---|---|
| 1 | Cxcl1 | 11.4846412 | 1.50E-03 | 1 | Gpr83 | -6.572772472 | 7.61E-15 |
| 2 | Tff1 | 11.15844395 | 3.57E-16 | 2 | Ighv1-53 | -6.443102027 | 4.84E-03 |
| 3 | Ednrb | 8.98934269 | 3.38E-09 | 3 | Igkv4-72 | -6.083601388 | 1.91E-02 |
| 4 | Fcrls | 8.885498046 | 1.39E-02 | 4 | Epx | -5.895988336 | 9.95E-06 |
| 5 | Col18a1 | 8.612669814 | 3.40E-07 | 5 | Noxa1 | -4.414282758 | 2.65E-02 |
| 6 | Emp1 | 8.592748791 | 7.53E-08 | 6 | 2610204G07Rik | -4.229170003 | 1.95E-06 |
| 7 | Hspa1a | 8.369699381 | 4.99E-41 | 7 | Gsta4 | -4.184093686 | 1.92E-07 |
| 8 | Jcad | 8.35937801 | 1.14E-06 | 8 | Tmem17 | -4.076616919 | 1.42E-02 |
| 9 | Ky | 8.345257789 | 3.57E-06 | 9 | Aicda | -4.055049277 | 3.65E-02 |
| 10 | Hspa1b | 8.337559048 | 1.30737E-22 | 10 | Dnah7a | -4.003052513 | 3.84E-03 |
| 11 | Col3a1 | 8.215569972 | 2.45095E-05 | 11 | Traip | -3.578080465 | 1.09E-02 |

| | | | | | | | |
|---|---|---|---|---|---|---|---|
| 12 | Pcsk1 | 7.996936646 | 5.99274E-12 | 12 | Art2b | -3.537985175 | 9.71E-04 |
| 13 | Kcna4 | 7.927997105 | 1.69293E-05 | 13 | Trbv12-1 | -3.346234472 | 2.55E-07 |
| 14 | Ahnak2 | 7.860584597 | 0.000397644 | 14 | Ear1 | -3.318503178 | 9.69E-03 |
| 15 | Dab2 | 7.737180559 | 0.00011502 | 15 | Atp9a | -3.229633184 | 3.66E-02 |
| 16 | Ier3 | 7.734885396 | 1.75E-09 | 16 | F830016B08Rik | -3.183134869 | 2.89E-03 |
| 17 | Comp | 7.699892268 | 5.90E-05 | 17 | Ccr9 | -3.125372976 | 4.68E-02 |
| 18 | 4933407L21Rik | 7.52238626 | 0.000157744 | 18 | Gm8113 | -3.037111791 | 2.64E-02 |
| 19 | Hlf | 7.415732483 | 5.68697E-05 | 19 | Mis18bp1 | -2.948118665 | 1.41E-04 |
| 20 | Cxcl2 | 7.30783506 | 6.58697E-31 | 20 | Fam160a1 | -2.942144042 | 1.14E-03 |
| 21 | Tmtc2 | 7.256696068 | 0.000483412 | 21 | Gm14085 | -2.885045425 | 2.07E-03 |
| 22 | Stab1 | 7.235883033 | 1.72E-11 | 22 | Pars2 | -2.866230227 | 1.84E-02 |
| 23 | Muc4 | 7.207588925 | 0.000630259 | 23 | Bambi-ps1 | -2.798963735 | 3.87E-02 |
| 24 | Fmod | 7.203478589 | 0.002536669 | 24 | Gm17764 | -2.791099847 | 2.57E-02 |
| 25 | Adam12 | 7.14744242 | 0.000134351 | 25 | Izumo1r | -2.669008618 | 2.30E-11 |
| 26 | Hsbp1l1 | 7.108153705 | 0.000140502 | 26 | Slc7a10 | -2.662652154 | 7.52E-03 |
| 27 | Cables1 | 7.022622514 | 0.000849299 | 27 | Ckap2l | -2.605058352 | 2.41E-03 |
| 28 | Gas2l3 | 7.009137072 | 3.36394E-06 | 28 | Gm4956 | -2.571228351 | 2.60E-08 |
| 29 | Mcub | 6.991155858 | 1.80189E-20 | 29 | Fgf13 | -2.555615022 | 4.42E-02 |
| 30 | Gm45774 | 6.96056685 | 0.000342994 | 30 | Cdca5 | -2.503442274 | 2.57E-03 |
| 31 | Glis2 | 6.940510159 | 6.30E-04 | 31 | Ifit1bl1 | -2.494981765 | 1.11E-02 |
| 32 | Mgp | 6.934440846 | 0.000624731 | 32 | Dnph1 | -2.488646649 | 3.13E-02 |
| 33 | Gm44851 | 6.903947167 | 0.000349904 | 33 | Abhd17c | -2.379926521 | 1.70E-02 |
| 34 | Arhgef28 | 6.887883222 | 0.005599278 | 34 | Capn3 | -2.292802921 | 1.84E-02 |
| 35 | Nmur1 | 6.808619146 | 5.48E-04 | 35 | Nuf2 | -2.203103699 | 4.20E-02 |
| 36 | Gng4 | 6.766943938 | 0.000825826 | 36 | P2rx7 | -2.192583726 | 9.16E-03 |
| 37 | Gm17590 | 6.704568645 | 0.004390986 | 37 | Tbc1d4 | -2.128810658 | 1.80E-02 |
| 38 | Npnt | 6.575805514 | 4.41396E-09 | 38 | Trbv13-1 | -2.116991902 | 2.71E-02 |
| 39 | Mt2 | 6.480765489 | 7.13E-17 | 39 | Shcbp1 | -2.086342323 | 4.64E-02 |
| 40 | Prune2 | 6.464332956 | 0.002029968 | 40 | Aurkb | -2.055983255 | 1.94E-02 |
| 41 | Dgat2 | 6.286657102 | 1.98E-08 | 41 | Tgtp1 | -2.01948297 | 4.38E-02 |
| 42 | Tm4sf1 | 6.149845254 | 0.00266846 | 42 | Iigp1 | -2.005967008 | 3.21E-05 |
| 43 | Ophn1 | 6.122186966 | 0.017379645 | 43 | Ccnb1 | -1.989040268 | 2.40E-02 |
| 44 | Gzmb | 6.096465676 | 1.26056E-21 | 44 | Serpina3f | -1.98038315 | 7.91E-03 |
| 45 | Itga3 | 6.095730942 | 3.06E-05 | 45 | Suv39h1 | -1.929198734 | 4.42E-02 |
| 46 | Csf2 | 6.093836579 | 0.003924431 | 46 | Ndc80 | -1.9211858 | 1.18E-02 |
| 47 | Mid2 | 6.069541259 | 6.04E-03 | 47 | Cdc45 | -1.907994352 | 4.45E-02 |
| 48 | Jam2 | 6.049483195 | 0.031111357 | 48 | Toe1 | -1.878438918 | 0.02834455 |
| 49 | Magi1 | 6.021235215 | 3.04E-02 | 49 | Cxcr3 | -1.847453879 | 1.95E-04 |
| 50 | Gm37488 | 5.975661965 | 0.031709345 | 50 | Shmt1 | -1.838682236 | 3.37E-02 |
| 51 | Gm37352 | 5.907088121 | 0.012390362 | 51 | Ephx1 | -1.821443425 | 5.03E-05 |
| 52 | Pld1 | 5.893318598 | 0.004855902 | 52 | Itm2a | -1.807230216 | 4.73E-02 |
| 53 | Fam129b | 5.864333279 | 0.000722155 | 53 | Bcl2 | -1.806915696 | 9.49E-03 |
| 54 | Bcar1 | 5.862506777 | 0.005478109 | 54 | Cep19 | -1.740444215 | 1.84E-02 |
| 55 | Rnf152 | 5.837556231 | 0.01566241 | 55 | Ssbp2 | -1.717645614 | 4.55E-03 |
| 56 | Sgip1 | 5.82123071 | 1.24672E-05 | 56 | Gpm6b | -1.699379621 | 1.49E-02 |
| 57 | Flt1 | 5.813706038 | 0.006271065 | 57 | Rrm2 | -1.695370626 | 8.26E-04 |
| 58 | Col1a2 | 5.81296624 | 4.77889E-07 | 58 | Ifi214 | -1.672602354 | 1.81E-02 |
| 59 | Lzts1 | 5.780438845 | 8.17E-03 | 59 | Trbv1 | -1.658640237 | 6.56E-03 |
| 60 | Il1rl1 | 5.759833005 | 7.5662E-13 | 60 | Elp3 | -1.61802281 | 2.04E-02 |
| 61 | Ceacam15 | 5.758208494 | 7.70711E-05 | 61 | Zfp429 | -1.563913235 | 2.32E-02 |

| | | | | | | | |
|---|---|---|---|---|---|---|---|
| 62 | Dcn | 5.628412553 | 2.48081E-05 | 62 | Tfb2m | -1.551189731 | 4.96E-02 |
| 63 | Sparc | 5.572664103 | 1.20E-08 | 63 | Rcn1 | -1.532179375 | 4.42E-02 |
| 64 | Epas1 | 5.438552241 | 1.15059E-05 | 64 | Rars | -1.527520761 | 2.39E-02 |
| 65 | Gm13684 | 5.408300833 | 2.58428E-05 | 65 | Gimap4 | -1.515574486 | 6.35E-04 |
| 66 | Mmp2 | 5.377956538 | 8.44E-03 | 66 | Dennd2d | -1.51455172 | 1.09E-03 |
| 67 | Cd34 | 5.344085787 | 1.03E-02 | 67 | Snip1 | -1.509591793 | 4.10E-02 |
| 68 | Nckap5 | 5.318239752 | 0.002515303 | 68 | Trbv3 | -1.48588276 | 2.40E-02 |
| 69 | Arnt2 | 5.304126792 | 2.04307E-06 | 69 | Lta | -1.451272185 | 0.0203693 |
| 70 | Gm8237 | 5.285327653 | 0.005473921 | 70 | B4galnt1 | -1.45107198 | 7.53E-03 |
| 71 | Tnfaip3 | 5.246997572 | 5.43446E-10 | 71 | Tgtp2 | -1.450982997 | 1.85E-02 |
| 72 | Gm48611 | 5.240293589 | 0.000140502 | 72 | Fech | -1.432243395 | 1.39E-02 |
| 73 | Frmd5 | 5.215671836 | 9.10E-06 | 73 | Gimap7 | -1.417663752 | 3.20E-02 |
| 74 | Gm38405 | 5.202337837 | 0.013452171 | 74 | Nudt19 | -1.400445984 | 1.52E-02 |
| 75 | Gm42684 | 5.199991212 | 0.037110296 | 75 | Serpina3g | -1.398199791 | 3.10E-02 |
| 76 | Ccr2 | 5.190582749 | 8.1499E-10 | 76 | Rcn2 | -1.394453176 | 3.83E-02 |
| 77 | Rgs11 | 5.166202857 | 0.002365207 | 77 | Ifi47 | -1.388030329 | 4.86E-03 |
| 78 | Wwtr1 | 5.091581275 | 0.005005106 | 78 | Scoc | -1.35701235 | 4.13E-02 |
| 79 | Il23r | 5.081753991 | 1.76E-04 | 79 | Tubg1 | -1.345363874 | 3.13E-02 |
| 80 | Gem | 5.069827246 | 4.63535E-10 | 80 | Cd96 | -1.318338263 | 0.04419804 |
| 81 | Gla | 5.057331409 | 1.04427E-15 | 81 | Farsb | -1.317008307 | 3.30E-02 |
| 82 | Mt1 | 5.031984264 | 1.15895E-36 | 82 | Trbc2 | -1.305730273 | 1.44E-02 |
| 83 | Jag1 | 4.964300241 | 4.68E-02 | 83 | Ruvbl1 | -1.303868022 | 4.66E-02 |
| 84 | Ppp1r3b | 4.950959415 | 0.00056731 | 84 | AI467606 | -1.302116969 | 2.21E-02 |
| 85 | Anxa11os | 4.949710216 | 0.001795092 | 85 | Cd27 | -1.294073978 | 1.74E-02 |
| 86 | Gm10800 | 4.941833843 | 0.001026649 | 86 | Pdcd4 | -1.289788511 | 1.07E-02 |
| 87 | Nr4a1 | 4.905999646 | 3.73953E-08 | 87 | Oat | -1.287025278 | 3.37E-02 |
| 88 | Cobl | 4.887525895 | 1.91E-02 | 88 | Tmem173 | -1.285985272 | 0.00787603 |
| 89 | Pomc | 4.882750698 | 1.15E-02 | 89 | Sh2d1a | -1.272873942 | 4.68E-02 |
| 90 | Zdhhc23 | 4.873352516 | 2.29E-03 | 90 | Mpv17 | -1.258076049 | 2.12E-02 |
| 91 | Lmna | 4.858300657 | 5.85206E-18 | 91 | Phgdh | -1.250817577 | 4.30E-02 |
| 92 | Zbtb46 | 4.844384197 | 1.05E-06 | 92 | Aaas | -1.156929665 | 4.79E-02 |
| 93 | Dscaml1 | 4.807492777 | 3.39E-05 | 93 | Eif3l | -1.067921726 | 3.42E-02 |
| 94 | Ccr5 | 4.748137341 | 1.31753E-08 | 94 | Irf1 | -1.035295458 | 4.10E-02 |
| 95 | Ccdc184 | 4.676263359 | 1.64017E-05 | | | | |
| 96 | Fancf | 4.633391963 | 1.37939E-07 | | | | |
| 97 | Cdkn1a | 4.607567145 | 6.98933E-22 | | | | |
| 98 | Acod1 | 4.583212384 | 0.000209975 | | | | |
| 99 | Ccl4 | 4.581218039 | 0.000151152 | | | | |
| 100 | Hspb1 | 4.534863543 | 2.66E-08 | | | | |
| 101 | Fbxl2 | 4.470783589 | 1.41E-02 | | | | |
| 102 | Ifrd1 | 4.455979618 | 2.00412E-07 | | | | |
| 103 | Plxnd1 | 4.449610334 | 3.47066E-05 | | | | |
| 104 | Hspa2 | 4.432400699 | 3.38701E-05 | | | | |
| 105 | Il17rb | 4.411156034 | 0.01484705 | | | | |
| 106 | Palm3 | 4.398534061 | 3.34E-03 | | | | |
| 107 | Dgat1 | 4.379422553 | 1.33124E-08 | | | | |

| | | | |
|---|---|---|---|
| 108 | Dusp4 | 4.321437969 | 1.67E-06 |
| 109 | Gm17056 | 4.296089137 | 8.26E-04 |
| 110 | Il18rap | 4.290611325 | 6.20E-08 |
| 111 | Tub | 4.279593691 | 0.011532852 |
| 112 | Rgs2 | 4.274977131 | 7.69675E-17 |
| 113 | Gpr55 | 4.270936516 | 1.23E-08 |
| 114 | AC160637.1 | 4.266678749 | 0.019406664 |
| 115 | Mmp14 | 4.259718071 | 0.011844432 |
| 116 | Bhlhe40 | 4.24101256 | 1.64106E-05 |
| 117 | Mlf1 | 4.239660656 | 8.53024E-06 |
| 118 | Adam8 | 4.220900386 | 2.51E-07 |
| 119 | Nxnl2 | 4.215070263 | 0.019065194 |
| 120 | Snora31 | 4.204184312 | 0.02522749 |
| 121 | Sv2c | 4.193708348 | 4.23517E-08 |
| 122 | Plvap | 4.182595711 | 1.39E-02 |
| 123 | Perp | 4.162226897 | 0.007657684 |
| 124 | Odc1 | 4.148203058 | 2.22221E-06 |
| 125 | Sytl2 | 4.14811482 | 1.8524E-07 |
| 126 | Gm16008 | 4.124474075 | 0.001913065 |
| 127 | Pmaip1 | 4.051033158 | 0.000141421 |
| 128 | Tnfrsf8 | 4.048742874 | 8.68727E-05 |
| 129 | Ern1 | 4.038387828 | 7.22E-04 |
| 130 | Echdc2 | 4.031249995 | 7.57E-03 |
| 131 | Phlda1 | 4.017536701 | 0.000872563 |
| 132 | Bambi | 4.005990313 | 0.000308999 |
| 133 | Gas2 | 3.995336249 | 6.07E-06 |
| 134 | Rab26 | 3.981963382 | 2.69883E-06 |
| 135 | Firre | 3.979104954 | 0.000735522 |
| 136 | Rora | 3.942380201 | 2.37031E-07 |
| 137 | Gm15283 | 3.938095657 | 0.019406664 |
| 138 | A830012C17Rik | 3.902781803 | 8.46E-04 |
| 139 | Pcyt1a | 3.900848592 | 7.95138E-08 |
| 140 | Gm45222 | 3.886945295 | 1.01E-03 |
| 141 | Ero1l | 3.882163426 | 1.64017E-05 |
| 142 | Gm38158 | 3.874881809 | 9.86E-03 |
| 143 | Ptpn5 | 3.856799131 | 1.04862E-06 |
| 144 | Il10 | 3.834284728 | 4.59742E-10 |
| 145 | Ets2 | 3.819354912 | 3.15763E-05 |
| 146 | Nfil3 | 3.815237404 | 0.001189287 |
| 147 | Hpn | 3.809664814 | 0.037110296 |
| 148 | Col1a1 | 3.794587131 | 0.011832584 |
| 149 | Havcr2 | 3.79127663 | 1.57164E-05 |
| 150 | Pim3 | 3.790804336 | 0.000680544 |
| 151 | Rasgrf1 | 3.785213564 | 0.016944772 |
| 152 | Gm45220 | 3.772977503 | 0.013115794 |
| 153 | Ttc39c | 3.751201783 | 1.13526E-05 |

| | | | |
|---|---|---|---|
| 154 | Tex2 | 3.746304398 | 3.20505E-05 |
| 155 | Cd80 | 3.74349052 | 2.89E-04 |
| 156 | Hist1h2bc | 3.740180076 | 2.90176E-17 |
| 157 | Dsel | 3.727706944 | 0.032826797 |
| 158 | Gdap10 | 3.717913969 | 0.005478109 |
| 159 | Gm3558 | 3.707876036 | 0.003007681 |
| 160 | Atf3 | 3.698481156 | 5.54865E-09 |
| 161 | Slco2b1 | 3.679493468 | 1.64E-02 |
| 162 | Gm13522 | 3.666300215 | 0.044138576 |
| 163 | Rnd1 | 3.662400705 | 0.002443322 |
| 164 | Crem | 3.662026653 | 1.97E-09 |
| 165 | Hist1h2bg | 3.656253889 | 1.71E-08 |
| 166 | G530011O06Rik | 3.650960402 | 1.12E-02 |
| 167 | Ptp4a1 | 3.602274076 | 3.56E-03 |
| 168 | Apaf1 | 3.578479618 | 3.21857E-05 |
| 169 | Hilpda | 3.577766917 | 2.38868E-16 |
| 170 | Nid2 | 3.56947071 | 6.85E-03 |
| 171 | Neat1 | 3.557413964 | 2.65E-06 |
| 172 | Dusp1 | 3.555838756 | 1.46228E-07 |
| 173 | Plk2 | 3.547653124 | 0.004506008 |
| 174 | Snord118 | 3.545784143 | 0.000599659 |
| 175 | B930095G15Rik | 3.544326057 | 0.037673518 |
| 176 | B130021K23Rik | 3.521944075 | 0.001533353 |
| 177 | Ramp3 | 3.507614106 | 0.001055261 |
| 178 | Fabp4 | 3.493020634 | 0.005952379 |
| 179 | Mmp25 | 3.490741463 | 0.016944772 |
| 180 | Dusp5 | 3.476534687 | 0.000630259 |
| 181 | Plppr3 | 3.457931871 | 0.043122416 |
| 182 | Fosl2 | 3.442150525 | 1.64106E-05 |
| 183 | 44447 | 3.422101596 | 0.00076269 |
| 184 | Gm45266 | 3.413872923 | 0.014983363 |
| 185 | Thbs1 | 3.408909952 | 0.039213085 |
| 186 | Arl5b | 3.406886722 | 6.86E-05 |
| 187 | Snx18 | 3.356175573 | 7.11632E-05 |
| 188 | Cxcl10 | 3.35301457 | 0.000283153 |
| 189 | Gm45223 | 3.351839066 | 0.000722155 |
| 190 | Plcd1 | 3.342580985 | 3.01E-03 |
| 191 | Dip2a | 3.336080382 | 2.19E-02 |
| 192 | Arrdc3 | 3.327722743 | 0.000825826 |
| 193 | Arl5a | 3.304908219 | 7.09394E-05 |
| 194 | Errfi1 | 3.292300558 | 1.85936E-05 |
| 195 | Raph1 | 3.289504518 | 2.12288E-05 |
| 196 | C030034L19Rik | 3.286437188 | 0.025973098 |
| 197 | Wdr66 | 3.260042572 | 3.85E-02 |
| 198 | Slc15a3 | 3.258449138 | 2.45411E-09 |
| 199 | Laptm4b | 3.257403821 | 7.51E-08 |

| 200 | Gm6225 | 3.247198961 | 0.006260715 |
|---|---|---|---|

**C. Neonatal tendon**

| | Upregulated | Neonatal tendon vs Adult tendon | | | Downregulated | Neonatal tendon vs Adult tendon | |
|---|---|---|---|---|---|---|---|
| | **Gene** | Log2(Fold change) | P value | | **Gene** | Log2(Fold change) | P value |
| 1 | Lyve1 | 10.9598885 | 5.64E-11 | 1 | Ighg1 | -8.972595717 | 3.88E-02 |
| 2 | Tm4sf19 | 9.366011546 | 2.67E-09 | 2 | Cr2 | -8.427131241 | 8.28E-03 |
| 3 | Olr1 | 9.34088448 | 2.42E-08 | 3 | Ighg2c | -8.404315625 | 2.67E-03 |
| 4 | Retnla | 9.340528156 | 2.06E-04 | 4 | Fcer2a | -8.106603271 | 3.35E-03 |
| 5 | Gm20056 | 8.817188878 | 8.18E-08 | 5 | Igkv1-135 | -7.984934972 | 3.26E-02 |
| 6 | Kctd19 | 8.718479279 | 4.39E-07 | 6 | Klrc1 | -7.67190416 | 2.31E-07 |
| 7 | Fabp3 | 8.564621404 | 3.07E-07 | 7 | Jchain | -7.359485967 | 1.22E-02 |
| 8 | Olfml2a | 8.3529054 | 2.14E-05 | 8 | Dscaml1 | -7.320004773 | 3.86E-06 |
| 9 | Dnm1 | 8.277679116 | 1.70E-06 | 9 | Il23r | -7.180313999 | 2.24E-05 |
| 10 | Gm42793 | 8.245197533 | 2.18242E-06 | 10 | Ighv5-17 | -6.893990444 | 2.16E-02 |
| 11 | Thbs2 | 8.027711342 | 0.032783184 | 11 | Prf1 | -6.762760554 | 6.98E-04 |
| 12 | Efcab6 | 7.984156784 | 4.84347E-05 | 12 | H2-DMb2 | -6.389309211 | 4.33E-03 |
| 13 | Nectin4 | 7.935628994 | 2.86213E-05 | 13 | Dtx1 | -6.284851468 | 2.04E-03 |
| 14 | Adgrg6 | 7.935364761 | 2.0181E-05 | 14 | Ly6k | -6.261368276 | 1.62E-03 |
| 15 | Tnfaip6 | 7.734666616 | 7.81955E-06 | 15 | Hbq1b | -6.237880421 | 4.35E-03 |
| 16 | Pyroxd2 | 7.645708788 | 1.08E-05 | 16 | Klrb1c | -6.153532522 | 2.10E-03 |
| 17 | Edn1 | 7.635660231 | 1.04E-05 | 17 | Prg2 | -6.148239512 | 2.94E-04 |
| 18 | Ctf2 | 7.562593026 | 0.000108547 | 18 | Treml4 | -6.032770315 | 3.18E-03 |
| 19 | Capn6 | 7.438938081 | 0.000161357 | 19 | Ncr1 | -6.025210363 | 7.90E-03 |
| 20 | Gm14461 | 7.419301419 | 0.000319209 | 20 | Klre1 | -5.825636001 | 2.03E-02 |
| 21 | Plekha4 | 7.410655206 | 0.000552797 | 21 | Dennd3 | -5.825488827 | 3.80E-03 |
| 22 | Btbd3 | 7.394018299 | 3.45E-04 | 22 | Ighd | -5.819609067 | 4.48E-06 |
| 23 | Ifi202b | 7.372047212 | 0.000345302 | 23 | Apol7e | -5.750530108 | 5.80E-03 |
| 24 | 9130230L23Rik | 7.352082158 | 0.000265822 | 24 | Igkv3-1 | -5.733152283 | 3.82E-02 |
| 25 | Gm12031 | 7.32204566 | 4.13808E-05 | 25 | Rasal1 | -5.692130543 | 7.55E-03 |
| 26 | Il1a | 7.169484686 | 0.015490157 | 26 | Klra13-ps | -5.683052282 | 3.67E-02 |
| 27 | Tnfsf18 | 7.143500995 | 0.001103349 | 27 | Camk2b | -5.601902935 | 1.61E-02 |
| 28 | Gm13391 | 7.070149649 | 0.000177133 | 28 | Tmem108 | -5.59384979 | 3.30E-02 |
| 29 | Cck | 7.024856561 | 0.005612215 | 29 | Fcrl5 | -5.577403219 | 3.06E-02 |
| 30 | Maob | 7.021980805 | 0.005829191 | 30 | Cxcr2 | -5.524009309 | 2.66E-02 |
| 31 | Gm5345 | 7.01266558 | 5.04E-03 | 31 | Cobl | -5.489319813 | 1.40E-02 |
| 32 | Gnat3 | 6.985423657 | 0.007035329 | 32 | Tmem63c | -5.41905085 | 4.17E-02 |
| 33 | Cd163 | 6.969582254 | 1.89453E-14 | 33 | Ear1 | -5.335798056 | 2.08E-02 |
| 34 | Dhrs9 | 6.94696694 | 0.000161439 | 34 | Unc5cl | -5.14984867 | 6.72E-03 |
| 35 | Nova1 | 6.931119859 | 7.45E-03 | 35 | Rhag | -5.103384251 | 1.14E-02 |
| 36 | Cd300ld2 | 6.920543772 | 0.001832045 | 36 | Gm32856 | -5.080398114 | 4.97E-03 |
| 37 | Pxdn | 6.913040501 | 0.000100097 | 37 | Fcrl1 | -5.04830001 | 4.06E-04 |
| 38 | Vsig4 | 6.899947775 | 0.015271556 | 38 | Blk | -4.99561038 | 2.20E-03 |
| 39 | Stox2 | 6.878701212 | 3.50E-03 | 39 | Trav10 | -4.995049346 | 2.93E-02 |
| 40 | Mmp13 | 6.831553645 | 4.95141E-05 | 40 | Hemgn | -4.847112584 | 4.82E-02 |
| 41 | Timp3 | 6.782850328 | 2.45E-06 | 41 | Tbxa2r | -4.676655765 | 1.85E-02 |
| 42 | Etv1 | 6.776131604 | 1.29977E-05 | 42 | Kynu | -4.517594595 | 2.38E-02 |
| 43 | Clec14a | 6.722814828 | 0.000244682 | 43 | H2-Ob | -4.507585396 | 6.19E-04 |
| 44 | Gprc5c | 6.692415614 | 2.46528E-10 | 44 | Serpini1 | -4.474593704 | 2.24E-03 |
| 45 | Fam20c | 6.688013231 | 1.33E-04 | 45 | Ctsw | -4.404106864 | 1.70E-06 |

| | | | | | | | |
|---|---|---|---|---|---|---|---|
| 46 | Scamp5 | 6.53562355 | 2.6685E-09 | 46 | Invs | -4.399227841 | 5.55E-04 |
| 47 | Map3k6 | 6.506708777 | 9.31E-04 | 47 | Gpr18 | -4.27018692 | 2.09E-03 |
| 48 | C4b | 6.500697212 | 3.49247E-12 | 48 | Gm6225 | -4.233996575 | 0.02298811 |
| 49 | Ptgs2 | 6.494979732 | 1.58E-19 | 49 | Cd96 | -4.088317139 | 3.77E-04 |
| 50 | Wwc1 | 6.463755714 | 0.002290727 | 50 | Ms4a1 | -4.009562999 | 7.54E-03 |
| 51 | Gm29340 | 6.421317966 | 0.00127601 | 51 | Ar | -4.006883698 | 1.21E-02 |
| 52 | Ak1 | 6.364071594 | 0.003375037 | 52 | Rhebl1 | -3.975820097 | 5.96E-03 |
| 53 | NA | 6.341704518 | 6.23073E-09 | 53 | Trim10 | -3.838489594 | 1.68E-02 |
| 54 | Tmem202 | 6.321277758 | 0.000148603 | 54 | Dkkl1 | -3.829920222 | 8.27E-03 |
| 55 | Pstpip2 | 6.283838965 | 0.000208276 | 55 | AI838599 | -3.795060973 | 4.78E-02 |
| 56 | 4930512H18Rik | 6.279683699 | 0.002302834 | 56 | Ighm | -3.769007504 | 2.17E-04 |
| 57 | Slc7a2 | 6.246070751 | 0.000277324 | 57 | Hp | -3.728443793 | 2.62E-02 |
| 58 | H19 | 6.20474021 | 4.60227E-18 | 58 | Btd | -3.716280527 | 1.32E-02 |
| 59 | Igf2bp2 | 6.188055108 | 4.17E-03 | 59 | Trbv13-3 | -3.656695062 | 4.71E-03 |
| 60 | 2310043M15Rik | 6.179109731 | 0.012135181 | 60 | Napsa | -3.641165626 | 7.66E-03 |
| 61 | Gpr157 | 6.140254531 | 0.000234504 | 61 | Ccr6 | -3.615092663 | 5.48E-09 |
| 62 | Afap1l1 | 6.096473174 | 8.37606E-06 | 62 | Samd10 | -3.566056282 | 6.15E-03 |
| 63 | Clmp | 6.081718424 | 1.69E-03 | 63 | Snord118 | -3.42184804 | 3.06E-02 |
| 64 | Ms4a14 | 6.073938678 | 0.043596858 | 64 | Gm12216 | -3.359653942 | 1.88E-02 |
| 65 | Fbn1 | 6.047312319 | 5.67469E-05 | 65 | Hsh2d | -3.332836329 | 1.03E-02 |
| 66 | Saa3 | 6.009867414 | 2.09E-05 | 66 | Zfp472 | -3.322879361 | 3.41E-02 |
| 67 | F13a1 | 5.95310018 | 7.99E-20 | 67 | 1110034G24Rik | -3.319396974 | 2.49E-02 |
| 68 | Adora3 | 5.943699307 | 0.011061019 | 68 | Ms4a4b | -3.295774967 | 5.04E-05 |
| 69 | Fosl1 | 5.940611462 | 0.000122697 | 69 | Bicd1 | -3.289481071 | 0.02524917 |
| 70 | Cd276 | 5.934383994 | 1.36909E-05 | 70 | Ttn | -3.203553754 | 3.97E-04 |
| 71 | Mill2 | 5.9175856 | 0.022576777 | 71 | St6galnac3 | -3.165575029 | 1.89E-03 |
| 72 | Igf2 | 5.902040496 | 3.17857E-07 | 72 | Zfp692 | -3.132948722 | 2.66E-03 |
| 73 | Tppp | 5.884733323 | 4.13E-05 | 73 | Cd226 | -3.126022346 | 1.51E-03 |
| 74 | Fbln2 | 5.87664435 | 0.038773976 | 74 | Gypa | -3.092995203 | 1.57E-02 |
| 75 | B3galnt1 | 5.819335055 | 7.50027E-07 | 75 | Serpina3g | -3.028714139 | 3.18E-04 |
| 76 | Serpinb8 | 5.814067034 | 9.13394E-07 | 76 | A830012C17Rik | -2.984504817 | 1.86E-02 |
| 77 | Tlr3 | 5.802614069 | 0.001453198 | 77 | Trat1 | -2.958101194 | 2.24E-03 |
| 78 | Pdgfc | 5.774094916 | 3.18894E-08 | 78 | Gpank1 | -2.938700057 | 4.76E-02 |
| 79 | Colec12 | 5.752548242 | 7.91E-09 | 79 | Xlr4c | -2.895169906 | 6.07E-04 |
| 80 | Cfh | 5.741576412 | 3.6491E-08 | 80 | 2610035D17Rik | -2.894012042 | 0.00753943 |
| 81 | F830208F22Rik | 5.716508449 | 0.002549665 | 81 | Gpr15 | -2.862413956 | 1.74E-02 |
| 82 | Tlr13 | 5.66713029 | 0.000344848 | 82 | Gprin3 | -2.845975309 | 2.76E-02 |
| 83 | Alox5 | 5.640913305 | 7.92E-06 | 83 | Tff1 | -2.841386001 | 1.28E-03 |
| 84 | Atg4c | 5.591759034 | 4.7618E-07 | 84 | Grhpr | -2.828594774 | 6.54E-03 |
| 85 | Csf3 | 5.577797978 | 1.28062E-06 | 85 | E130308A19Rik | -2.806694585 | 4.18E-02 |
| 86 | Epb41l1 | 5.557795679 | 0.000318026 | 86 | Il1r2 | -2.806363269 | 3.65E-08 |
| 87 | Nrep | 5.540715721 | 4.34539E-08 | 87 | Gm19585 | -2.803049807 | 4.02E-04 |
| 88 | Ifit2 | 5.532094204 | 1.96E-12 | 88 | Cd1d1 | -2.69962728 | 0.00112574 |
| 89 | Mamdc2 | 5.531776674 | 5.31E-03 | 89 | Camkmt | -2.677381974 | 4.39E-02 |
| 90 | Cd109 | 5.511695795 | 2.06E-06 | 90 | Galnt12 | -2.57073942 | 1.13E-02 |
| 91 | Cd209g | 5.502428593 | 0.000133475 | 91 | Gm28100 | -2.545219857 | 4.17E-02 |
| 92 | Slc13a3 | 5.501808766 | 6.84E-06 | 92 | Atic | -2.49347164 | 3.43E-02 |

| | | | | | | | |
|---|---|---|---|---|---|---|---|
| 93 | BC022687 | 5.500206613 | 9.09E-04 | 93 | Tuba4a | -2.490917272 | 4.07E-02 |
| 94 | Cygb | 5.481429755 | 2.1519E-05 | 94 | Trbv13-2 | -2.449627438 | 2.25E-02 |
| 95 | Dab2 | 5.472146578 | 6.0926E-08 | 95 | Slco3a1 | -2.430677731 | 3.82E-02 |
| 96 | Ptprb | 5.468513359 | 0.00017215 | 96 | Pvrig | -2.418799343 | 5.26E-03 |
| 97 | Lhfpl2 | 5.466680742 | 1.04378E-08 | 97 | 1110038F14Rik | -2.412620744 | 3.01E-02 |
| 98 | Gas6 | 5.433992567 | 1.38167E-06 | 98 | Gimap4 | -2.407573739 | 1.69E-04 |
| 99 | Abca9 | 5.417290472 | 1.51178E-09 | 99 | Pglyrp1 | -2.382813632 | 1.23E-03 |
| 100 | C1qtnf1 | 5.367169838 | 5.80E-04 | 100 | Uros | -2.381242005 | 0.04392807 |
| 101 | Cgref1 | 5.346072101 | 2.30E-02 | 101 | Ccdc28a | -2.369112672 | 4.37E-02 |
| 102 | F10 | 5.275770642 | 0.002741659 | 102 | Lsm5 | -2.366500738 | 0.01971363 |
| 103 | Tnmd | 5.250544589 | 4.69555E-06 | 103 | Exosc7 | -2.352395117 | 3.05E-02 |
| 104 | Sgms2 | 5.249288025 | 1.24722E-07 | 104 | Mir142hg | -2.331380379 | 2.11E-02 |
| 105 | Cfd | 5.244509822 | 0.032935028 | 105 | Podnl1 | -2.326975308 | 4.22E-02 |
| 106 | Gm5431 | 5.207459432 | 5.35E-04 | 106 | Ly6a | -2.287272847 | 6.69E-05 |
| 107 | Tlr2 | 5.197835994 | 2.88679E-12 | 107 | Rpf1 | -2.277897678 | 3.67E-02 |
| 108 | Csf1r | 5.187394292 | 3.27E-06 | 108 | Btla | -2.249564543 | 4.28E-02 |
| 109 | Il15 | 5.163753878 | 1.17E-08 | 109 | Gimap1os | -2.236547529 | 3.63E-02 |
| 110 | Mfsd7a | 5.162488278 | 4.91E-03 | 110 | Ttc39c | -2.235044537 | 3.77E-02 |
| 111 | Cpne8 | 5.120243304 | 0.00017215 | 111 | Sell | -2.228138372 | 1.67E-02 |
| 112 | Pigz | 5.111609757 | 2.69786E-07 | 112 | Thap3 | -2.228035448 | 4.69E-03 |
| 113 | A4galt | 5.10680566 | 2.19E-03 | 113 | Trim59 | -2.221353197 | 1.72E-02 |
| 114 | Hlx | 5.084150803 | 6.06715E-06 | 114 | Snhg20 | -2.207894662 | 1.78E-02 |
| 115 | Pmp22 | 5.083060896 | 1.08831E-07 | 115 | Gbp8 | -2.205374339 | 1.07E-02 |
| 116 | Frmd4a | 5.053652026 | 3.54789E-07 | 116 | Ptprcap | -2.18737523 | 8.07E-03 |
| 117 | Gm14636 | 5.052085981 | 5.71439E-09 | 117 | Hsd11b1 | -2.166489124 | 2.46E-03 |
| 118 | Itga5 | 5.051793338 | 6.32E-11 | 118 | Arpc5l | -2.162074383 | 1.55E-02 |
| 119 | Marcksl1 | 5.049201703 | 1.89453E-14 | 119 | Sh2d1a | -2.130798621 | 1.51E-02 |
| 120 | Cfb | 5.047890674 | 3.17857E-07 | 120 | Oard1 | -2.112280253 | 1.68E-02 |
| 121 | Eln | 5.033677787 | 0.009299501 | 121 | Gm26740 | -2.095492236 | 2.73E-03 |
| 122 | Tspan7 | 5.022477906 | 5.35E-04 | 122 | Cd3d | -2.076589579 | 3.84E-04 |
| 123 | Met | 4.963183312 | 0.000509184 | 123 | Pold4 | -2.076543189 | 3.59E-02 |
| 124 | Thbs1 | 4.959959787 | 3.33391E-05 | 124 | Mthfsl | -2.067375913 | 1.36E-02 |
| 125 | Fblim1 | 4.954461707 | 0.000761784 | 125 | Tasp1 | -2.062000001 | 2.45E-02 |
| 126 | Rsad2 | 4.934542614 | 2.51565E-09 | 126 | Gimap5 | -2.060108646 | 1.32E-02 |
| 127 | Stard8 | 4.93200113 | 3.93096E-12 | 127 | Cd52 | -2.047778042 | 3.56E-03 |
| 128 | Ropn1l | 4.925849427 | 0.041730192 | 128 | Mrpl32 | -2.039374559 | 1.94E-03 |
| 129 | Fkbp10 | 4.922609814 | 2.39E-03 | 129 | Slamf6 | -2.039035508 | 1.31E-02 |
| 130 | P2ry2 | 4.922456175 | 4.13E-05 | 130 | Mif4gd | -2.036102832 | 5.04E-03 |
| 131 | Acod1 | 4.922337283 | 9.92008E-10 | 131 | Fam189b | -2.031423902 | 8.02E-03 |
| 132 | Qpct | 4.915052389 | 2.0181E-05 | 132 | Sesn3 | -2.016742851 | 3.01E-02 |
| 133 | Tlr4 | 4.912966264 | 2.51E-06 | 133 | Xlr4a | -2.012458243 | 1.29E-02 |
| 134 | Npl | 4.903327889 | 5.39155E-05 | 134 | Ctla2a | -2.01087719 | 1.44E-03 |
| 135 | Tns3 | 4.893173266 | 7.50027E-07 | 135 | Borcs8 | -1.992170663 | 3.12E-02 |
| 136 | Plk2 | 4.890230159 | 3.61024E-12 | 136 | Cd3g | -1.976886318 | 2.62E-04 |
| 137 | Tlr8 | 4.882666283 | 0.000525234 | 137 | Pla2g16 | -1.968874618 | 1.22E-02 |
| 138 | Fgd5 | 4.882454645 | 5.92E-03 | 138 | Cers4 | -1.965014305 | 4.36E-02 |

| | | | | | | | |
|---|---|---|---|---|---|---|---|
| 139 | Best1 | 4.869881778 | 0.006719909 | 139 | Gimap9 | -1.962183271 | 5.25E-03 |
| 140 | Fndc1 | 4.864919232 | 3.12E-04 | 140 | Bmyc | -1.948749739 | 2.52E-02 |
| 141 | Oasl1 | 4.855294035 | 1.38713E-05 | 141 | Ltb | -1.939584884 | 1.72E-03 |
| 142 | Pik3r6 | 4.854913941 | 5.62E-04 | 142 | Traf3ip3 | -1.936028172 | 2.39E-02 |
| 143 | C5ar1 | 4.84980048 | 0.030579659 | 143 | Trbc2 | -1.931929126 | 2.58E-03 |
| 144 | Anpep | 4.826013309 | 0.000288019 | 144 | Limd2 | -1.924439737 | 8.20E-03 |
| 145 | Ifih1 | 4.81983481 | 1.03564E-12 | 145 | Lockd | -1.924195856 | 1.05E-02 |
| 146 | Rgl1 | 4.813290224 | 4.45455E-07 | 146 | Pura | -1.923886775 | 3.56E-02 |
| 147 | Plau | 4.795721042 | 1.54204E-06 | 147 | Cd2 | -1.916285669 | 9.18E-04 |
| 148 | Tnfrsf11a | 4.789511384 | 1.14139E-05 | 148 | Spn | -1.906278443 | 7.76E-03 |
| 149 | Gfra2 | 4.776416458 | 0.024923103 | 149 | H2-T23 | -1.904177977 | 4.75E-03 |
| 150 | Ace | 4.762541678 | 0.006983531 | 150 | Penk | -1.89143017 | 1.16E-03 |
| 151 | Mrc1 | 4.754711865 | 2.30951E-07 | 151 | Mmp9 | -1.890311097 | 3.06E-02 |
| 152 | Rin2 | 4.754390554 | 2.67086E-09 | 152 | Ikzf4 | -1.884133863 | 4.32E-02 |
| 153 | Col1a1 | 4.751562574 | 3.7989E-05 | 153 | Rps27 | -1.875247008 | 1.98E-02 |
| 154 | Jup | 4.743461392 | 6.31931E-11 | 154 | Clint1 | -1.864900604 | 2.16E-02 |
| 155 | Gpt2 | 4.741945142 | 7.37E-04 | 155 | Tspan32 | -1.862678435 | 1.59E-03 |
| 156 | Gab1 | 4.741429272 | 0.001629919 | 156 | AW112010 | -1.852957804 | 6.76E-04 |
| 157 | Il1rn | 4.728415296 | 7.47272E-06 | 157 | Myl12b | -1.852509897 | 3.33E-03 |
| 158 | Fgfr1 | 4.725513832 | 3.78982E-06 | 158 | Emg1 | -1.833752437 | 4.85E-02 |
| 159 | Rnd2 | 4.72180351 | 0.005738652 | 159 | Sla2 | -1.833660531 | 1.08E-02 |
| 160 | Hspg2 | 4.701575108 | 6.10654E-05 | 160 | Ccndbp1 | -1.830526688 | 4.81E-02 |
| 161 | Gm14052 | 4.681326256 | 1.46E-03 | 161 | Spint2 | -1.828427587 | 5.83E-03 |
| 162 | Fam198b | 4.673666484 | 0.020489193 | 162 | Cisd3 | -1.818166546 | 3.36E-02 |
| 163 | Sh3pxd2b | 4.668703522 | 8.64663E-07 | 163 | Tnfrsf18 | -1.817829159 | 5.72E-03 |
| 164 | Fnip2 | 4.659050954 | 4.11E-08 | 164 | Laptm4b | -1.809657933 | 2.05E-02 |
| 165 | Gm45167 | 4.65687366 | 4.14E-02 | 165 | Glipr1 | -1.808166748 | 0.00955382 |
| 166 | Zfhx3 | 4.65410783 | 1.42E-05 | 166 | Sorl1 | -1.783334871 | 0.04464447 |
| 167 | Rassf8 | 4.646029365 | 5.95E-03 | 167 | Fkbp3 | -1.772934807 | 1.01E-02 |
| 168 | Rab31 | 4.64095712 | 6.00742E-13 | 168 | Psip1 | -1.760883461 | 3.63E-02 |
| 169 | Aoah | 4.615572821 | 0.000101499 | 169 | BE692007 | -1.759910997 | 2.26E-02 |
| 170 | Ehd2 | 4.615088772 | 1.24E-06 | 170 | Satb1 | -1.758856325 | 4.69E-02 |
| 171 | Cyr61 | 4.611813345 | 7.40E-07 | 171 | Ppcdc | -1.738352933 | 2.27E-02 |
| 172 | Lifr | 4.611717249 | 7.45208E-06 | 172 | Tmem154 | -1.734299843 | 2.76E-02 |
| 173 | Rasal2 | 4.590097576 | 0.001065326 | 173 | Ccdc12 | -1.729560025 | 2.85E-02 |
| 174 | Emp1 | 4.587315912 | 1.89453E-14 | 174 | Slc50a1 | -1.710204021 | 9.10E-03 |
| 175 | 4932438H23Rik | 4.585110008 | 0.012989674 | 175 | Gimap6 | -1.702819333 | 4.30E-02 |
| 176 | Rab11fip5 | 4.582543137 | 1.90847E-05 | 176 | Cd5 | -1.691134766 | 3.42E-02 |
| 177 | Il13ra1 | 4.581027609 | 1.57946E-05 | 177 | Izumo4 | -1.684096072 | 3.92E-02 |
| 178 | Postn | 4.580856583 | 4.12492E-05 | 178 | Sirt7 | -1.68039241 | 3.97E-02 |
| 179 | Gm43351 | 4.580557682 | 0.026984445 | 179 | Tbc1d10c | -1.679885157 | 8.39E-03 |
| 180 | Dock1 | 4.573687102 | 0.000892677 | 180 | Acap1 | -1.672875116 | 1.62E-02 |
| 181 | Gm26888 | 4.56255201 | 0.020527006 | 181 | Coro1a | -1.659564812 | 3.82E-02 |
| 182 | Dpysl3 | 4.561016029 | 0.001914799 | 182 | Stap1 | -1.655870376 | 2.26E-02 |
| 183 | Lrp12 | 4.558220503 | 0.00019339 | 183 | Gstp1 | -1.654487125 | 4.05E-02 |
| 184 | Gatm | 4.55606087 | 7.13307E-08 | 184 | Emb | -1.650625587 | 2.05E-02 |

| | | | | | | | |
|---|---|---|---|---|---|---|---|
| 185 | Dse | 4.555311996 | 6.31931E-11 | 185 | A430093F15Rik | -1.646213164 | 3.56E-02 |
| 186 | Slc35f5 | 4.54447884 | 1.04E-08 | 186 | Skap1 | -1.628607784 | 2.71E-02 |
| 187 | Cd33 | 4.544357833 | 1.67509E-05 | 187 | Tnfrsf9 | -1.625204413 | 3.77E-02 |
| 188 | P2rx7 | 4.53606475 | 1.96692E-10 | 188 | Trac | -1.624427062 | 3.87E-03 |
| 189 | Iqsec2 | 4.516122361 | 0.016190397 | 189 | Cdk2ap2 | -1.615260833 | 4.35E-02 |
| 190 | Oasl2 | 4.512166865 | 9.84E-08 | 190 | Tspan13 | -1.607949996 | 5.25E-03 |
| 191 | Entpd3 | 4.505054062 | 2.07E-02 | 191 | Ctla4 | -1.566919449 | 3.91E-02 |
| 192 | Wdfy3 | 4.504622524 | 1.69982E-06 | 192 | Cd247 | -1.566212535 | 4.00E-02 |
| 193 | Tbc1d24 | 4.503601308 | 0.000456242 | 193 | Lck | -1.559137861 | 1.22E-02 |
| 194 | Liph | 4.492725765 | 0.020457617 | 194 | Rnf167 | -1.551011983 | 1.73E-02 |
| 195 | Tlr6 | 4.489841339 | 3.86567E-05 | 195 | AC149090.1 | -1.545687127 | 1.58E-02 |
| 196 | Ogn | 4.474940937 | 0.030441123 | 196 | Rexo1 | -1.545330186 | 3.99E-02 |
| 197 | Pdpn | 4.471461128 | 2.30E-03 | 197 | Lsp1 | -1.54113538 | 3.97E-02 |
| 198 | Lrp1 | 4.457354968 | 7.29151E-06 | 198 | Lat | -1.536198468 | 4.36E-02 |
| 199 | Siglec1 | 4.438437632 | 2.52E-04 | 199 | Tma7 | -1.533600971 | 4.84E-02 |
| 200 | Sema6a | 4.437564023 | 0.024923103 | 200 | Fam173a | -1.52574063 | 3.99E-02 |

**D. Neonatal spleen**

| | Upregulated | Neonatal tendon vs Adult spleen | | | Downregulated | Neonatal spleen vs Adult spleen | |
|---|---|---|---|---|---|---|---|
| | **Gene** | Log2(Fold change) | P value | | **Gene** | Log2(Fold change) | P value |
| 1 | Ryr2 | 8.217576426 | 4.72E-08 | 1 | Ighg2c | -15.62667893 | 3.14E-63 |
| 2 | Grm8 | 7.519752865 | 2.68E-06 | 2 | Ighg2b | -13.03589257 | 2.98E-22 |
| 3 | Arhgap32 | 7.099944389 | 2.29E-05 | 3 | Igkv3-7 | -11.02772951 | 6.54E-06 |
| 4 | B3galt5 | 6.729406453 | 1.38E-04 | 4 | Igkv4-72 | -10.97787864 | 7.67E-75 |
| 5 | Klk1 | 6.381809149 | 6.09E-17 | 5 | Igkv5-39 | -10.97078961 | 3.12E-32 |
| 6 | Runx2os2 | 6.267089389 | 8.87E-04 | 6 | Ighv5-17 | -10.84496933 | 2.75E-19 |
| 7 | Vdr | 6.2053448 | 2.71E-04 | 7 | Ighv3-6 | -10.29340446 | 7.54E-41 |
| 8 | B3galnt1 | 5.842906409 | 1.09E-03 | 8 | Igkv3-4 | -10.19738383 | 9.90E-18 |
| 9 | Dsg2 | 5.555319099 | 2.02E-03 | 9 | Igkv4-55 | -10.11505246 | 1.18E-07 |
| 10 | Fam129b | 5.499187939 | 0.000176096 | 10 | Ighv1-76 | -9.976267583 | 9.36E-07 |
| 11 | Chml | 5.436256585 | 0.003060707 | 11 | Ighv1-5 | -9.743894326 | 1.15E-02 |
| 12 | Itga7 | 5.431327163 | 0.000160195 | 12 | Ighg3 | -9.700607724 | 6.23E-09 |
| 13 | Pir | 5.396835362 | 0.002820356 | 13 | Igkv10-96 | -9.542120801 | 1.49E-24 |
| 14 | Cdh1 | 5.283514326 | 0.000510723 | 14 | Ighv1-80 | -9.348154585 | 2.20E-06 |
| 15 | Rgs11 | 5.059183286 | 1.00123E-05 | 15 | Ighv1-59 | -9.274153791 | 1.04E-03 |
| 16 | Obscn | 4.997154783 | 2.04E-03 | 16 | Igkv13-85 | -9.014515051 | 1.78E-13 |
| 17 | Plxdc1 | 4.970047382 | 5.17E-10 | 17 | Ighv3-1 | -9.007128756 | 1.27E-04 |
| 18 | Ankrd13b | 4.942088363 | 0.001154199 | 18 | Igkv3-5 | -8.935714217 | 4.23E-16 |
| 19 | Lefty1 | 4.748338589 | 0.000234691 | 19 | Ighv1-55 | -8.872977022 | 3.49E-18 |
| 20 | Gm5086 | 4.68327304 | 0.007780591 | 20 | Igkv4-57 | -8.805668122 | 1.08E-05 |
| 21 | Siglech | 4.567052457 | 1.03438E-26 | 21 | Ighv1-78 | -8.51400315 | 1.65E-03 |
| 22 | Gas2l3 | 4.562588264 | 5.24E-03 | 22 | Ighv1-9 | -8.158483569 | 9.47E-07 |
| 23 | Ptprf | 4.404875733 | 2.22226E-06 | 23 | Gm20429 | -8.131344176 | 1.36E-09 |
| 24 | Olfr164 | 4.399282093 | 0.000110495 | 24 | Ighv1-74 | -7.935756277 | 7.34E-06 |
| 25 | Zfat | 4.367663961 | 0.002439525 | 25 | Ighv8-8 | -7.890921636 | 3.35E-05 |
| 26 | Adcy6 | 4.31006415 | 2.11454E-05 | 26 | Igkv1-117 | -7.877345599 | 2.19E-16 |
| 27 | Gm37101 | 4.309734765 | 0.000833061 | 27 | Igkv3-2 | -7.816281951 | 1.12E-13 |
| 28 | Il17rb | 4.25372195 | 0.003225482 | 28 | Klra8 | -7.793637345 | 1.38E-08 |

| | | | | | | | |
|---|---|---|---|---|---|---|---|
| 29 | Gm12253 | 4.23534367 | 0.007103773 | 29 | Klra4 | -7.724322819 | 3.88E-03 |
| 30 | Cbx2 | 4.122171968 | 8.75211E-05 | 30 | Ighv1-53 | -7.633425489 | 1.80E-07 |
| 31 | Clec10a | 4.058223935 | 2.78E-06 | 31 | Gm43388 | -7.626886659 | 3.57E-08 |
| 32 | Gm17354 | 4.030932913 | 0.005263868 | 32 | Igkv8-24 | -7.596211528 | 7.32E-27 |
| 33 | Jade3 | 3.988715827 | 0.009286422 | 33 | Igkv10-94 | -7.558783806 | 1.44E-08 |
| 34 | Lrp8 | 3.978284957 | 1.94169E-05 | 34 | Igkv4-57-1 | -7.545450367 | 4.83E-10 |
| 35 | Amigo3 | 3.94166654 | 5.24E-04 | 35 | Ighv1-69 | -7.534436455 | 6.59E-08 |
| 36 | Nphp3 | 3.918000815 | 0.013241253 | 36 | Igkv3-12 | -7.509235793 | 2.39E-09 |
| 37 | Gm9752 | 3.89517758 | 0.000292177 | 37 | Ms4a3 | -7.447873321 | 2.94E-02 |
| 38 | Mmp14 | 3.884188179 | 0.002169779 | 38 | Igkv4-86 | -7.419666322 | 2.27E-07 |
| 39 | Dapk1 | 3.863126608 | 5.54E-04 | 39 | Rgs13 | -7.353752105 | 1.23E-06 |
| 40 | Vangl2 | 3.857911248 | 0.005839448 | 40 | Iglv1 | -7.325989657 | 1.33E-19 |
| 41 | Chdh | 3.830990181 | 3.54E-03 | 41 | Igkv8-19 | -7.261216942 | 2.46E-10 |
| 42 | Uhrf1bp1 | 3.797382204 | 0.011394883 | 42 | Ighv1-52 | -7.238316089 | 1.95E-08 |
| 43 | Ercc6l | 3.71506283 | 0.000181864 | 43 | Klre1 | -7.20288721 | 1.35E-06 |
| 44 | Gm48611 | 3.709635475 | 0.010193213 | 44 | Cxcr2 | -7.174704962 | 8.51E-11 |
| 45 | Mfsd4a | 3.656402623 | 1.70E-09 | 45 | Igkv1-110 | -7.151016517 | 1.87E-09 |
| 46 | Trbj1-4 | 3.598302745 | 0.000517908 | 46 | Igkv8-28 | -7.145447195 | 9.77E-08 |
| 47 | Sipa1l3 | 3.55556223 | 3.74E-06 | 47 | Oosp1 | -7.124145122 | 2.03E-06 |
| 48 | Vcl | 3.539096313 | 0.003880267 | 48 | Ighv6-3 | -7.007529814 | 4.4541E-11 |
| 49 | Usp28 | 3.531346408 | 1.19E-20 | 49 | Igkv3-10 | -6.995167636 | 1.81E-14 |
| 50 | Wdr62 | 3.522894327 | 0.000239122 | 50 | Ighv7-3 | -6.935112522 | 9.77E-04 |
| 51 | Plekhg5 | 3.51759927 | 0.003795859 | 51 | Myof | -6.8990257 | 1.92E-09 |
| 52 | Cd209d | 3.506854974 | 2.06347E-05 | 52 | Igkv6-23 | -6.881457526 | 8.53E-10 |
| 53 | Palm3 | 3.489590519 | 0.006191192 | 53 | Igkv19-93 | -6.861119205 | 4.54E-09 |
| 54 | Drc1 | 3.487950777 | 0.009585392 | 54 | Ighv1-18 | -6.841750532 | 2.51E-04 |
| 55 | Ddr1 | 3.424601652 | 7.90554E-05 | 55 | Igkv13-84 | -6.702838051 | 4.17E-12 |
| 56 | Fbxl22 | 3.42258119 | 0.013373644 | 56 | Aicda | -6.697694291 | 1.79E-06 |
| 57 | Gm43361 | 3.394077105 | 0.001547624 | 57 | Ighv1-58 | -6.691654457 | 3.41E-03 |
| 58 | Plxnd1 | 3.335372865 | 0.001784873 | 58 | Igkv1-133 | -6.680527648 | 5.94E-06 |
| 59 | Zc3h3 | 3.318733534 | 8.02E-05 | 59 | Iglv2 | -6.655799598 | 2.92E-17 |
| 60 | Zfp653 | 3.31062646 | 0.000139454 | 60 | Clec5a | -6.632798891 | 6.05E-06 |
| 61 | Prune1 | 3.286974189 | 0.000382571 | 61 | Clec4e | -6.555817274 | 3.04E-06 |
| 62 | Gm44321 | 3.274008625 | 0.033793288 | 62 | Igkv3-1 | -6.494978698 | 2.30E-05 |
| 63 | Ccr9 | 3.273908249 | 4.28E-04 | 63 | Igkc | -6.488229196 | 7.64E-99 |
| 64 | Gm27252 | 3.272292501 | 0.005263868 | 64 | Igkv4-91 | -6.487598303 | 3.37E-14 |
| 65 | Paqr8 | 3.244073319 | 0.000416227 | 65 | Gzma | -6.468504338 | 5.10E-06 |
| 66 | C230085N15Rik | 3.231244471 | 4.44E-08 | 66 | Pnlip | -6.467810769 | 1.87E-06 |
| 67 | Rap1gap2 | 3.226141278 | 6.37E-04 | 67 | Jchain | -6.441839486 | 7.21E-09 |
| 68 | Iqgap3 | 3.215247101 | 0.009272728 | 68 | Igkv4-80 | -6.425593298 | 2.85E-03 |
| 69 | Gabrr2 | 3.1848603 | 2.21825E-05 | 69 | Ear1 | -6.4178019 | 0.00075538 |
| 70 | Gm24187 | 3.182413788 | 0.003126569 | 70 | Igkv1-135 | -6.399891278 | 1.11E-20 |
| 71 | Dntt | 3.153894944 | 0.004913117 | 71 | Igkv8-27 | -6.277697216 | 6.75E-07 |
| 72 | Ddias | 3.149859096 | 0.008636047 | 72 | Ear6 | -6.27472174 | 9.54E-06 |
| 73 | Slco4a1 | 3.146462556 | 6.41E-05 | 73 | Igkv12-41 | -6.143202115 | 3.16E-15 |
| 74 | Zfp142 | 3.143003773 | 7.6131E-07 | 74 | Igkv17-121 | -6.136178698 | 1.87E-21 |
| 75 | Tex2 | 3.141604954 | 4.73322E-11 | 75 | Igkv6-15 | -6.12170069 | 1.04E-07 |
| 76 | Zfp169 | 3.11810818 | 3.49648E-07 | 76 | Igkv4-58 | -6.003997659 | 7.79E-06 |

| | | | | | | | |
|---|---|---|---|---|---|---|---|
| 77 | Car5b | 3.113048626 | 0.000294375 | 77 | Btnl10 | -5.975019814 | 9.03E-06 |
| 78 | Sh2b2 | 3.101318713 | 0.013976107 | 78 | Fcer2a | -5.931073839 | 1.23E-05 |
| 79 | Trim21 | 3.077535327 | 7.48E-16 | 79 | Klra7 | -5.91734313 | 3.29E-07 |
| 80 | AC160637.1 | 3.058454261 | 0.008044798 | 80 | Shank1 | -5.904720105 | 1.5776E-06 |
| 81 | Lmntd2 | 3.051657593 | 0.005629297 | 81 | Igkv12-89 | -5.888604436 | 2.22E-06 |
| 82 | Lars2 | 3.049814448 | 0.005892965 | 82 | Igkv16-104 | -5.887478778 | 1.25E-10 |
| 83 | Uty | 3.04833804 | 1.27E-04 | 83 | Igkv6-20 | -5.871488085 | 4.29E-11 |
| 84 | Sacm1l | 3.0135696 | 7.09277E-09 | 84 | Ighv1-72 | -5.858496643 | 1.43E-10 |
| 85 | Frmd5 | 3.007604221 | 0.010216795 | 85 | Iglc1 | -5.854492142 | 7.78E-10 |
| 86 | Plppr3 | 3.005354363 | 0.007103773 | 86 | Igkv2-137 | -5.800098681 | 1.92E-08 |
| 87 | Man2b2 | 3.003490172 | 1.57569E-08 | 87 | Klra13-ps | -5.796974927 | 1.28E-05 |
| 88 | Srgap3 | 2.986338684 | 5.78E-04 | 88 | Igkv14-111 | -5.784364109 | 5.2293E-05 |
| 89 | Xrcc2 | 2.981877917 | 7.80E-03 | 89 | Cd300e | -5.778617827 | 4.07E-13 |
| 90 | Abi2 | 2.970251423 | 5.22E-04 | 90 | Igkv6-32 | -5.715496117 | 7.34E-11 |
| 91 | Slc22a17 | 2.968229952 | 0.02276608 | 91 | Cr2 | -5.712194846 | 1.71E-32 |
| 92 | Slc29a4 | 2.956886541 | 2.20E-02 | 92 | Eaf2 | -5.633212325 | 3.99E-12 |
| 93 | Prkag2os1 | 2.95634033 | 2.00E-02 | 93 | Igkv6-17 | -5.620682961 | 2.65E-21 |
| 94 | Adam11 | 2.955573729 | 0.001145271 | 94 | Ighv1-26 | -5.585099528 | 4.49E-06 |
| 95 | Gm42449 | 2.948867793 | 0.014697044 | 95 | Igkv12-44 | -5.566941376 | 2.39E-16 |
| 96 | Zfp26 | 2.939630971 | 3.14099E-09 | 96 | Gm4841 | -5.555817006 | 1.25E-06 |
| 97 | Ankrd29 | 2.938721685 | 0.041924159 | 97 | Trem3 | -5.512139994 | 6.23E-09 |
| 98 | Il1rl1 | 2.925526805 | 1.14782E-09 | 98 | Igkv8-21 | -5.499678678 | 6.26E-09 |
| 99 | Gm15564 | 2.923269836 | 0.008740348 | 99 | Cldn13 | -5.496696979 | 1.35E-11 |
| 100 | Bsn | 2.872669764 | 6.53E-03 | 100 | Sirpb1c | -5.474802864 | 7.0824E-06 |
| 101 | Lifr | 2.86411324 | 1.54E-04 | 101 | Prg2 | -5.473944566 | 4.26E-03 |
| 102 | Rps4l | 2.821217777 | 0.000734383 | 102 | Cxcl9 | -5.472144417 | 9.9441E-05 |
| 103 | Grhl1 | 2.819902125 | 0.029192053 | 103 | Ighv1-82 | -5.452008125 | 7.12E-08 |
| 104 | Adamts6 | 2.817331704 | 1.86102E-06 | 104 | Ighv9-3 | -5.41443969 | 1.28E-03 |
| 105 | Gm23935 | 2.811773425 | 0.002333143 | 105 | Igkv6-25 | -5.322388179 | 2.72E-18 |
| 106 | Itga5 | 2.801735532 | 2.74E-02 | 106 | Igkv5-48 | -5.253245274 | 1.39E-06 |
| 107 | H6pd | 2.799312083 | 0.000213525 | 107 | Ell3 | -5.207943894 | 6.45E-06 |
| 108 | Trim62 | 2.797835268 | 2.41E-02 | 108 | Igkv4-68 | -5.19073408 | 7.56E-07 |
| 109 | Rasip1 | 2.794972666 | 2.52E-02 | 109 | Ppbp | -5.183123857 | 1.16E-20 |
| 110 | Mctp2 | 2.794034281 | 4.82E-06 | 110 | Cela2a | -5.174830919 | 4.15E-07 |
| 111 | Gm20342 | 2.788842358 | 0.000101251 | 111 | 2210010C04Rik | -5.130199806 | 1.66E-04 |
| 112 | Gpsm2 | 2.78878364 | 1.01392E-07 | 112 | Gcsam | -5.114401043 | 5.25E-07 |
| 113 | Frmd6 | 2.782509107 | 4.21E-08 | 113 | Wfdc17 | -5.082236263 | 3.53E-20 |
| 114 | Galnt4 | 2.779505201 | 0.000198594 | 114 | Iglc2 | -5.062747263 | 8.95E-92 |
| 115 | Trerf1 | 2.76263423 | 3.55466E-06 | 115 | Igkv9-120 | -5.030764252 | 6.99E-04 |
| 116 | Gm39323 | 2.752825517 | 2.62594E-05 | 116 | Igkv14-100 | -4.991553922 | 5.02E-05 |
| 117 | Mdm1 | 2.748417153 | 0.002610256 | 117 | Hp | -4.957189415 | 2.90E-14 |
| 118 | Firre | 2.73875865 | 1.23E-02 | 118 | Sostdc1 | -4.939097114 | 1.33E-06 |
| 119 | Nr4a1 | 2.734773736 | 1.38863E-09 | 119 | Igkv12-46 | -4.887030818 | 5.59E-03 |
| 120 | Gm48855 | 2.718466701 | 0.040595647 | 120 | Gm42870 | -4.885861088 | 8.24E-09 |
| 121 | Klf11 | 2.717020371 | 0.002104641 | 121 | Ctrb1 | -4.881199926 | 6.83E-04 |
| 122 | Gm29112 | 2.706962094 | 1.41E-03 | 122 | Pla2g2d | -4.844759093 | 1.47E-16 |

| | | | | | | | |
|---|---|---|---|---|---|---|---|
| 123 | Gm43544 | 2.695910847 | 0.042356593 | 123 | Epx | -4.839209552 | 3.65E-03 |
| 124 | Gm43313 | 2.69308163 | 0.023161866 | 124 | Cd40 | -4.833820657 | 1.42E-23 |
| 125 | Sema4d | 2.689172234 | 1.87155E-27 | 125 | Hepacam2 | -4.833174367 | 2.39E-07 |
| 126 | Mir6236 | 2.687699817 | 0.015604724 | 126 | Prg3 | -4.832283592 | 9.98E-12 |
| 127 | Colq | 2.679140824 | 6.53721E-06 | 127 | Il1b | -4.790972397 | 1.71E-06 |
| 128 | Tacc2 | 2.679010318 | 0.010025299 | 128 | Il15 | -4.742770539 | 1.26E-04 |
| 129 | Ankrd52 | 2.678911934 | 1.35E-04 | 129 | Gm31243 | -4.731650852 | 1.18E-15 |
| 130 | Sytl2 | 2.673364137 | 3.57E-10 | 130 | Gm15356 | -4.718398622 | 1.13E-03 |
| 131 | Gsk3a | 2.67237985 | 0.01311567 | 131 | Slc7a10 | -4.695940986 | 5.09E-07 |
| 132 | Txlnb | 2.666674085 | 0.049150107 | 132 | Igkv2-109 | -4.69559961 | 7.16E-08 |
| 133 | Mtmr10 | 2.666298327 | 3.31E-04 | 133 | Gm16170 | -4.694741791 | 1.65E-04 |
| 134 | Pomt1 | 2.659598705 | 0.004149939 | 134 | Bank1 | -4.693818214 | 3.82E-32 |
| 135 | Syne3 | 2.655636977 | 0.000153618 | 135 | Wfdc21 | -4.687974032 | 1.91E-09 |
| 136 | Setdb1 | 2.653120766 | 9.32247E-09 | 136 | Alox5 | -4.674914851 | 1.41E-03 |
| 137 | Spag5 | 2.650973435 | 0.000160195 | 137 | Ifi205 | -4.664950066 | 4.66E-04 |
| 138 | Usp33 | 2.637779164 | 1.11E-07 | 138 | Prtn3 | -4.642504667 | 2.82E-02 |
| 139 | Gm37529 | 2.633732057 | 0.007009941 | 139 | Slpi | -4.588499343 | 9.05E-37 |
| 140 | Dlg3 | 2.63222234 | 2.90E-03 | 140 | Fcmr | -4.57614417 | 9.17E-48 |
| 141 | Aqp3 | 2.62825712 | 0.035183734 | 141 | Rhd | -4.556769085 | 1.94E-17 |
| 142 | Rbl2 | 2.622229425 | 3.14E-12 | 142 | Igha | -4.551723546 | 1.39E-02 |
| 143 | Paqr5 | 2.621155222 | 0.022887021 | 143 | Ighd | -4.546235989 | 4.27E-61 |
| 144 | Focad | 2.618057594 | 0.001188725 | 144 | Crisp3 | -4.525168061 | 2.04E-02 |
| 145 | 4933407K13Rik | 2.615470314 | 0.004267056 | 145 | Sirpb1a | -4.515376724 | 1.32E-07 |
| 146 | Cacna1d | 2.611178025 | 0.018689958 | 146 | Anxa3 | -4.514823338 | 1.06E-07 |
| 147 | Acap3 | 2.611098213 | 0.004200489 | 147 | Scg5 | -4.497043941 | 1.22E-04 |
| 148 | Nat8f4 | 2.609497289 | 0.011666086 | 148 | Fcgr2b | -4.495135697 | 1.57E-21 |
| 149 | Fhl2 | 2.607814321 | 0.020665388 | 149 | H2-DMb2 | -4.478871089 | 1.19E-41 |
| 150 | Bub1b | 2.605419325 | 6.5588E-09 | 150 | Rasgrp3 | -4.433360958 | 6.93E-18 |
| 151 | Igsf9b | 2.604300156 | 0.000692466 | 151 | Rab30 | -4.432166674 | 3.00E-11 |
| 152 | Gm26835 | 2.601802591 | 0.008341476 | 152 | Pkhd1l1 | -4.415664016 | 8.55E-04 |
| 153 | Psd2 | 2.588760243 | 0.015042828 | 153 | Fcrl5 | -4.398686155 | 3.65E-02 |
| 154 | Dip2a | 2.58612181 | 0.023490039 | 154 | Oas1a | -4.398005005 | 4.68E-13 |
| 155 | Pdpr | 2.585389876 | 2.92E-03 | 155 | Rab7b | -4.381853717 | 4.92E-03 |
| 156 | Gm37645 | 2.559560241 | 0.033524156 | 156 | Klrk1 | -4.379060011 | 3.22E-06 |
| 157 | Hectd3 | 2.553298267 | 1.37651E-07 | 157 | Klrc2 | -4.374100063 | 7.57E-04 |
| 158 | Usp54 | 2.545430045 | 0.027983307 | 158 | 2010309G21Rik | -4.36956694 | 9.56E-06 |
| 159 | Leng8 | 2.539951495 | 2.85498E-11 | 159 | Gm4951 | -4.360714745 | 4.54E-06 |
| 160 | Emc1 | 2.535159889 | 0.00083482 | 160 | Cd79a | -4.334419427 | 5.46E-41 |
| 161 | Adcy3 | 2.531530597 | 3.96E-05 | 161 | Ear2 | -4.316351751 | 5.02E-22 |
| 162 | Gm28370 | 2.526799196 | 0.008634852 | 162 | Sdc1 | -4.315956226 | 1.25E-03 |
| 163 | Atp8b2 | 2.526475618 | 5.50596E-12 | 163 | Ffar1 | -4.310402005 | 1.11E-06 |
| 164 | Extl2 | 2.525554957 | 1.28E-03 | 164 | Ifitm6 | -4.302590843 | 6.25E-04 |
| 165 | Ercc5 | 2.524821348 | 1.72E-04 | 165 | Cd14 | -4.293277428 | 4.4684E-10 |
| 166 | Ficd | 2.518701314 | 1.59E-02 | 166 | Ly6k | -4.292720018 | 8.7156E-08 |
| 167 | Gm42141 | 2.516429859 | 1.72E-02 | 167 | Tinagl1 | -4.286586241 | 9.42E-04 |
| 168 | Slc35d1 | 2.51326569 | 3.874E-08 | 168 | Ighv5-16 | -4.28442176 | 1.16E-04 |

| | | | | | | | |
|---|---|---|---|---|---|---|---|
| 169 | Abcc4 | 2.506685888 | 1.35171E-06 | 169 | S100a9 | -4.278419762 | 7.66E-06 |
| 170 | Cog3 | 2.501641109 | 2.49E-09 | 170 | Nr3c2 | -4.260902511 | 5.26E-04 |
| 171 | Llgl1 | 2.498732201 | 4.83E-05 | 171 | Cd63 | -4.256218367 | 8.83E-13 |
| 172 | Ets2 | 2.484568031 | 9.17278E-05 | 172 | S100a8 | -4.228998452 | 9.54E-06 |
| 173 | Anxa11 | 2.478368661 | 0.001694293 | 173 | Mcemp1 | -4.21697021 | 4.47E-10 |
| 174 | Sppl2b | 2.475558971 | 6.23944E-06 | 174 | Il1rn | -4.212659753 | 1.05E-03 |
| 175 | Plxdc2 | 2.456445659 | 0.042459165 | 175 | Camp | -4.209470848 | 2.92E-05 |
| 176 | Galnt6 | 2.455113419 | 8.47099E-13 | 176 | Tspan8 | -4.19060068 | 5.68E-04 |
| 177 | Apaf1 | 2.453657067 | 1.68278E-05 | 177 | Igkv1-88 | -4.189848611 | 8.79E-04 |
| 178 | Slc25a22 | 2.453593893 | 3.77032E-05 | 178 | Cyp39a1 | -4.154787876 | 4.91E-03 |
| 179 | Ppfia4 | 2.45266127 | 0.000195665 | 179 | Ighv5-9-1 | -4.154383463 | 5.71E-04 |
| 180 | Dusp4 | 2.451138905 | 0.000176527 | 180 | Ubd | -4.143925477 | 1.88E-03 |
| 181 | Gm26538 | 2.450133443 | 0.028485851 | 181 | Lyz2 | -4.140010087 | 8.48E-42 |
| 182 | Miga2 | 2.449387464 | 0.015032119 | 182 | Gm6377 | -4.134138758 | 1.08E-04 |
| 183 | Zbtb46 | 2.449023888 | 0.013077784 | 183 | I830127L07Rik | -4.125680806 | 5.25E-07 |
| 184 | Ppfibp1 | 2.448540939 | 0.004101275 | 184 | Pycr1 | -4.123186959 | 4.68E-03 |
| 185 | Vps8 | 2.44722091 | 4.01093E-05 | 185 | 1600010M07Rik | -4.111666803 | 2.95E-04 |
| 186 | Parpbp | 2.447184527 | 1.59E-02 | 186 | Hes1 | -4.110257348 | 1.68E-05 |
| 187 | Fam196b | 2.439160706 | 0.000253692 | 187 | H2-Eb2 | -4.098032677 | 1.11E-05 |
| 188 | Upb1 | 2.429069343 | 0.029353694 | 188 | H2-M2 | -4.096877478 | 1.12E-04 |
| 189 | Gm16575 | 2.426731227 | 0.041542236 | 189 | C5ar1 | -4.050984579 | 2.80E-04 |
| 190 | Ahnak | 2.42549111 | 1.07E-23 | 190 | Prss2 | -4.040113876 | 2.33E-05 |
| 191 | Rarg | 2.416688959 | 2.48E-05 | 191 | Tacstd2 | -4.017593176 | 2.30E-03 |
| 192 | Antxr2 | 2.410493949 | 0.000107728 | 192 | Ngp | -4.016765233 | 1.01E-05 |
| 193 | Lig3 | 2.409030322 | 7.06516E-05 | 193 | BC028528 | -4.008693479 | 8.86E-14 |
| 194 | Arrdc3 | 2.40391974 | 0.000111434 | 194 | Tspan33 | -4.005904925 | 2.31E-03 |
| 195 | Dvl3 | 2.403383554 | 0.000252221 | 195 | Lcn2 | -4.005580929 | 8.82E-05 |
| 196 | Traj12 | 2.3942939 | 0.040186072 | 196 | 2900052N01Rik | -4.00377414 | 5.68E-04 |
| 197 | Sv2a | 2.394155042 | 2.26E-02 | 197 | Ighv14-2 | -3.982503095 | 3.44E-03 |
| 198 | Spata1 | 2.392485168 | 0.010408521 | 198 | Slfn14 | -3.946615918 | 2.74E-05 |
| 199 | Ankrd28 | 2.384279007 | 1.42E-03 | 199 | Cpb1 | -3.934071578 | 2.00E-03 |
| 200 | Nek2 | 2.383566045 | 1.10432E-05 | 200 | Tnfrsf13c | -3.909457984 | 1.15E-05 |

**Table S2**

Genesignatures unique to neonatal and adult tendon tregs.

Numbers represent the fold change in gene expression levels with corresponding P-values adjusted for multiple comparisons.

| A. Neonatal tendon Treg signature | | | | B. Adult tendon Treg signature | | | |
|---|---|---|---|---|---|---|---|
| | | Neonatal tendon vs Neonatal spleen | | | | Neonatal tendon vs Neonatal spleen | |
| | Gene | Log2(Fold change) | P value | | Gene | Log2(Fold change) | P value |
| 1 | Ccl7 | 16.11756401 | 3.24E-33 | 1 | Kcna4 | 7.927997105 | 1.69E-05 |
| 2 | Ccl2 | 15.04297441 | 9.36E-43 | 2 | Comp | 7.699892268 | 5.90E-05 |
| 3 | Pmp22 | 13.85656522 | 4.30E-44 | 3 | 4933407L21Rik | 7.52238626 | 1.58E-04 |
| 4 | Ccl12 | 13.72105455 | 6.01E-35 | 4 | Hlf | 7.415732483 | 5.69E-05 |
| 5 | Ccl8 | 13.54348036 | 4.51E-42 | 5 | Tmtc2 | 7.256696068 | 4.83E-04 |
| 6 | Ifnb1 | 13.40217462 | 8.33E-34 | 6 | Muc4 | 7.207588925 | 6.30E-04 |
| 7 | Clec4e | 13.22040876 | 1.19E-38 | 7 | Adam12 | 7.14744242 | 1.34E-04 |
| 8 | Gdf15 | 12.83266464 | 2.74E-31 | 8 | Hsbp1l1 | 7.108153705 | 1.41E-04 |
| 9 | C3ar1 | 12.64222572 | 2.74E-47 | 9 | Cables1 | 7.022622514 | 8.49E-04 |
| 10 | Clec4d | 12.30260428 | 6.04528E-15 | 10 | Glis2 | 6.940510159 | 6.30E-04 |
| 11 | Spp1 | 12.03273785 | 3.25153E-73 | 11 | Arhgef28 | 6.887883222 | 5.60E-03 |
| 12 | Cbr2 | 11.96442607 | 8.43578E-15 | 12 | Nmur1 | 6.808619146 | 5.48E-04 |
| 13 | H19 | 11.89183113 | 7.30298E-28 | 13 | Gm17590 | 6.704568645 | 4.39E-03 |
| 14 | Cxcl3 | 11.81247959 | 2.14632E-14 | 14 | Itga3 | 6.095730942 | 3.06E-05 |
| 15 | Ptgs2 | 11.77531257 | 1.28889E-16 | 15 | Mid2 | 6.069541259 | 6.04E-03 |
| 16 | Il6 | 11.57485 | 1.43E-24 | 16 | Gm37488 | 5.975661965 | 3.17E-02 |
| 17 | Il1rn | 11.52706704 | 5.68E-11 | 17 | Rnf152 | 5.837556231 | 1.57E-02 |
| 18 | Cyr61 | 11.44411573 | 7.16048E-25 | 18 | Flt1 | 5.813706038 | 6.27E-03 |
| 19 | Gas6 | 11.36501262 | 1.48708E-50 | 19 | Lzts1 | 5.780438845 | 8.17E-03 |
| 20 | Csf3 | 11.21059771 | 9.67917E-09 | 20 | Ceacam15 | 5.758208494 | 7.71E-05 |
| 21 | Cfh | 11.16563035 | 2.73681E-47 | 21 | Gm13684 | 5.408300833 | 2.58E-05 |
| 22 | Ms4a14 | 10.96368788 | 1.94E-25 | 22 | Nckap5 | 5.318239752 | 2.52E-03 |
| 23 | Folr2 | 10.84057863 | 6.82167E-19 | 23 | Gm8237 | 5.285327653 | 5.47E-03 |
| 24 | Plau | 10.68372292 | 1.73969E-23 | 24 | Gm38405 | 5.202337837 | 1.35E-02 |
| 25 | Lyve1 | 10.59651415 | 1.27172E-09 | 25 | Gm42684 | 5.199991212 | 3.71E-02 |
| 26 | Cav1 | 10.49769221 | 9.68567E-29 | 26 | Il23r | 5.081753991 | 1.76E-04 |
| 27 | Col6a1 | 10.42415032 | 3.83073E-17 | 27 | Gm10800 | 4.941833843 | 1.03E-03 |

| | | | | | | | |
|---|---|---|---|---|---|---|---|
| 28 | Pdpn | 10.33497969 | 1.62346E-16 | 28 | Cobl | 4.887525895 | 1.91E-02 |
| 29 | Il1a | 10.33250942 | 3.68095E-46 | 29 | Pomc | 4.882750698 | 1.15E-02 |
| 30 | 8430408G22Rik | 10.15755831 | 3.52801E-18 | 30 | Dscaml1 | 4.807492777 | 3.39E-05 |
| 31 | Flrt3 | 10.13811396 | 1.55E-16 | 31 | Ccdc184 | 4.676263359 | 1.64E-05 |
| 32 | Tnmd | 10.06990681 | 2.80177E-06 | 32 | Tub | 4.279593691 | 1.15E-02 |
| 33 | Tcim | 9.939662471 | 5.89612E-17 | 33 | Mlf1 | 4.239660656 | 8.53E-06 |
| 34 | Col5a2 | 9.891104683 | 0.000226087 | 34 | Nxnl2 | 4.215070263 | 1.91E-02 |
| 35 | Trem2 | 9.837350247 | 4.21E-26 | 35 | Snora31 | 4.204184312 | 2.52E-02 |
| 36 | Creb5 | 9.83661342 | 1.38954E-16 | 36 | Sv2c | 4.193708348 | 4.24E-08 |
| 37 | Tm4sf19 | 9.824153814 | 1.64763E-16 | 37 | Perp | 4.162226897 | 7.66E-03 |
| 38 | Gja1 | 9.688496272 | 8.78765E-31 | 38 | Gm16008 | 4.124474075 | 1.91E-03 |
| 39 | Cd14 | 9.68325677 | 1.62E-16 | 39 | Gas2 | 3.995336249 | 6.07E-06 |
| 40 | Adamts1 | 9.664856974 | 1.38024E-13 | 40 | Gm15283 | 3.938095657 | 1.94E-02 |
| 41 | P2ry2 | 9.628041056 | 1.02E-21 | 41 | A830012C17Rik | 3.902781803 | 8.46E-04 |
| 42 | Colec12 | 9.609077402 | 2.23786E-20 | 42 | Ptpn5 | 3.856799131 | 1.05E-06 |
| 43 | Fbn1 | 9.544775666 | 3.56563E-12 | 43 | Hpn | 3.809664814 | 3.71E-02 |
| 44 | Col12a1 | 9.46560151 | 1.28048E-11 | 44 | Gm45220 | 3.772977503 | 1.31E-02 |
| 45 | Rhoj | 9.459294196 | 1.59E-14 | 45 | Ttc39c | 3.751201783 | 1.14E-05 |
| 46 | Ifi205 | 9.458293328 | 6.61073E-08 | 46 | Dsel | 3.727706944 | 3.28E-02 |
| 47 | Lum | 9.403529786 | 2.17E-05 | 47 | Gm3558 | 3.707876036 | 3.01E-03 |
| 48 | Mmp19 | 9.351511736 | 9.93918E-14 | 48 | Gm13522 | 3.666300215 | 0.044138576 |
| 49 | Rab7b | 9.35140251 | 2.65E-35 | 49 | Snord118 | 3.545784143 | 6.00E-04 |
| 50 | Serpinb8 | 9.328222009 | 3.05754E-17 | 50 | B930095G15Rik | 3.544326057 | 3.77E-02 |
| 51 | Fndc1 | 9.311520388 | 0.000160997 | 51 | Ramp3 | 3.507614106 | 1.06E-03 |
| 52 | Pf4 | 9.304974869 | 1.16026E-51 | 52 | Mmp25 | 3.490741463 | 1.69E-02 |
| 53 | Igf2 | 9.297426687 | 1.11572E-19 | 53 | Gm45266 | 3.413872923 | 1.50E-02 |
| 54 | Fstl1 | 9.285708086 | 5.63678E-22 | 54 | Plcd1 | 3.342580985 | 3.01E-03 |
| 55 | Arg2 | 9.253196169 | 5.66665E-15 | 55 | C030034L19Rik | 3.286437188 | 2.60E-02 |
| 56 | Cd209g | 9.229710944 | 2.41799E-12 | 56 | Wdr66 | 3.260042572 | 3.85E-02 |
| 57 | Lhfpl2 | 9.166594294 | 9.11729E-19 | 57 | Gm6225 | 3.247198961 | 6.26E-03 |
| 58 | C5ar1 | 9.166550207 | 9.58877E-13 | 58 | Cnnm2 | 3.216964936 | 9.51E-03 |
| 59 | Pxdn | 9.160479451 | 1.10E-03 | 59 | BC049352 | 3.202751409 | 3.83E-02 |
| 60 | Nxpe5 | 9.125973118 | 3.22866E-13 | 60 | AI838599 | 3.188475882 | 3.74E-02 |
| 61 | Chp2 | 9.115743402 | 2.49306E-16 | 61 | Gm43065 | 3.169378997 | 3.35E-02 |
| 62 | Kctd19 | 9.11426844 | 4.68892E-12 | 62 | Gm47050 | 3.103205587 | 1.39E-02 |

| | | | | | | | |
|---|---|---|---|---|---|---|---|
| 63 | C4b | 9.083126853 | 7.26E-32 | 63 | Smarcd3 | 3.076947392 | 4.20E-02 |
| 64 | Tnfaip6 | 9.080019622 | 4.92695E-12 | 64 | Gm28100 | 2.948234229 | 3.01E-03 |
| 65 | Thbs4 | 9.068067737 | 0.000597563 | 65 | Slc52a3 | 2.946668366 | 3.21E-02 |
| 66 | Slc7a11 | 9.008413612 | 2.68E-05 | 66 | Nav2 | 2.827867324 | 8.31E-04 |
| 67 | Lpar1 | 8.994118281 | 3.53E-18 | 67 | Nebl | 2.753775071 | 1.71E-03 |
| 68 | Edn1 | 8.990135682 | 7.67488E-12 | 68 | Fam185a | 2.69519352 | 3.13E-02 |
| 69 | Nrep | 8.985042201 | 1.66165E-16 | 69 | Cep85l | 2.681560162 | 0.014206162 |
| 70 | Bgn | 8.98027527 | 8.85464E-23 | 70 | Gm16201 | 2.664135637 | 4.64E-02 |
| 71 | 4930430E12Rik | 8.950899631 | 1.57704E-22 | 71 | Gm16091 | 2.642497985 | 2.74E-02 |
| 72 | Wfdc17 | 8.946011811 | 1.11263E-54 | 72 | Hist3h2ba | 2.612457195 | 1.07E-03 |
| 73 | Reps2 | 8.905190527 | 5.36E-12 | 73 | Farp1 | 2.60480205 | 3.75E-02 |
| 74 | Car13 | 8.874317655 | 3.82307E-15 | 74 | Myo3b | 2.589639571 | 5.45E-03 |
| 75 | C1qtnf3 | 8.818029701 | 0.000806455 | 75 | Ric1 | 2.565156025 | 1.60E-02 |
| 76 | Mfap5 | 8.813384537 | 1.39202E-09 | 76 | 2210408F21Rik | 2.549329823 | 2.09E-02 |
| 77 | Hspg2 | 8.802896402 | 0.000118 | 77 | Gm9888 | 2.480401313 | 4.20E-02 |
| 78 | Slc13a3 | 8.787533805 | 2.08941E-19 | 78 | B9d1 | 2.470554108 | 4.78E-03 |
| 79 | Cd209f | 8.765534232 | 2.46E-05 | 79 | Cdkl3 | 2.454279026 | 4.20E-02 |
| 80 | Cxcl16 | 8.754986654 | 2.39726E-48 | 80 | Pde4d | 2.446283152 | 0.008326809 |
| 81 | Igfbp5 | 8.748992892 | 5.25794E-11 | 81 | 4930503L19Rik | 2.421419085 | 1.29E-03 |
| 82 | Rab34 | 8.747495063 | 1.79949E-10 | 82 | Ttn | 2.395367471 | 1.39E-02 |
| 83 | Ccl9 | 8.746967269 | 2.63E-46 | 83 | Birc2 | 2.37109764 | 5.59E-03 |
| 84 | Aplnr | 8.739028758 | 1.64765E-10 | 84 | Syne2 | 2.332773756 | 1.74E-02 |
| 85 | Gm15726 | 8.714598808 | 8.81034E-16 | 85 | Gm15506 | 2.313323322 | 4.28E-02 |
| 86 | Bicc1 | 8.701053434 | 6.7194E-09 | 86 | Scyl2 | 2.313024589 | 1.58E-02 |
| 87 | Oasl1 | 8.686652797 | 9.70842E-68 | 87 | Rapsn | 2.282813519 | 2.71E-03 |
| 88 | Ch25h | 8.674061022 | 2.30E-10 | 88 | Kif13b | 2.279553793 | 0.018583641 |
| 89 | Tnfsf15 | 8.660815618 | 6.09E-16 | 89 | Ctla4 | 2.278035613 | 5.24E-04 |
| 90 | Olfml2a | 8.625612159 | 1.26E-08 | 90 | 2410089E03Rik | 2.249614916 | 3.80E-02 |
| 91 | Fabp3 | 8.624267625 | 5.68865E-15 | 91 | Slc4a8 | 2.205174991 | 1.94E-02 |
| 92 | Gm42793 | 8.624104436 | 1.27E-10 | 92 | Ar | 2.171927413 | 1.54E-02 |
| 93 | Il15 | 8.591137204 | 5.53E-23 | 93 | Penk | 2.168341124 | 2.43E-07 |
| 94 | Clec5a | 8.586503371 | 1.8981E-05 | 94 | P2ry10 | 2.116439124 | 1.58E-02 |

| | | | | | | | |
|---|---|---|---|---|---|---|---|
| 95 | Tmem171 | 8.581584042 | 9.81205E-12 | 95 | Hemk1 | 1.928758313 | 2.29E-02 |
| 96 | Alox5 | 8.534472455 | 4.17672E-17 | 96 | 1810026B05Rik | 1.910504594 | 4.27E-03 |
| 97 | Slc7a2 | 8.527243027 | 1.18585E-09 | 97 | Slc36a3os | 1.898132618 | 4.02E-03 |
| 98 | Cfb | 8.523997628 | 8.38065E-17 | 98 | Man2a2 | 1.891397166 | 2.43E-02 |
| 99 | Gm48065 | 8.488491716 | 3.34531E-09 | 99 | Tmem123 | 1.890994978 | 3.30E-02 |
| 100 | Mir99ahg | 8.415808184 | 2.14E-12 | 100 | Tmx4 | 1.882899017 | 0.020044967 |
| 101 | Sparcl1 | 8.392755359 | 1.57E-16 | 101 | Appl2 | 1.863362513 | 2.38E-02 |
| 102 | Postn | 8.38685843 | 3.71474E-06 | 102 | 4933427D14Rik | 1.852862019 | 0.01030272 |
| 103 | Cracr2b | 8.378836799 | 1.20582E-09 | 103 | Padi2 | 1.851621819 | 1.39E-02 |
| 104 | Atp13a4 | 8.369627178 | 1.50449E-09 | 104 | Atxn1 | 1.794618946 | 1.42E-02 |
| 105 | Gm13391 | 8.366841057 | 7.4878E-09 | 105 | Pvrig | 1.784317891 | 1.32E-02 |
| 106 | Car3 | 8.344486096 | 5.72E-08 | 106 | Zfp839 | 1.784022719 | 4.41E-02 |
| 107 | Col4a1 | 8.326925806 | 7.40251E-07 | 107 | Atf6 | 1.783985406 | 3.84E-03 |
| 108 | Rassf8 | 8.314603557 | 6.78E-09 | 108 | Aebp2 | 1.753309225 | 3.76E-02 |
| 109 | Efcab6 | 8.278675765 | 4.51E-08 | 109 | Klrc1 | 1.734347188 | 4.97E-02 |
| 110 | Ms4a7 | 8.265358233 | 6.46E-49 | 110 | Mynn | 1.731964109 | 1.43E-02 |
| 111 | Cxcl11 | 8.186972184 | 2.02861E-08 | 111 | Dnajb13 | 1.712471847 | 1.05E-02 |
| 112 | Col4a2 | 8.185357734 | 4.61987E-25 | 112 | Vamp2 | 1.689497276 | 4.13E-02 |
| 113 | Sh3pxd2b | 8.148878226 | 2.68E-05 | 113 | Emb | 1.626805068 | 3.82E-03 |
| 114 | Serpine1 | 8.138713416 | 9.12769E-08 | 114 | 2610005L07Rik | 1.595163473 | 4.44E-02 |
| 115 | Gpnmb | 8.13575179 | 3.62606E-14 | 115 | Tnk2 | 1.580621439 | 2.35E-02 |
| 116 | Adgrf5 | 8.13572828 | 5.24206E-08 | 116 | Myo1e | 1.571349897 | 3.71E-02 |
| 117 | Meox2 | 8.129598388 | 1.66755E-08 | 117 | Sh2b1 | 1.565649638 | 3.73E-02 |
| 118 | C1qtnf1 | 8.123781046 | 1.52E-08 | 118 | Itpkb | 1.563129558 | 3.94E-02 |
| 119 | Tlr8 | 8.121243531 | 1.27058E-12 | 119 | Wnk1 | 1.472120395 | 3.21E-02 |
| 120 | Arpin | 8.103316402 | 2.3989E-08 | 120 | Slc25a19 | 1.326201018 | 3.75E-02 |
| 121 | Npl | 8.093011968 | 2.70978E-07 | 121 | Eif4e3 | 1.248030132 | 4.37E-02 |
| 122 | Myof | 8.076890501 | 2.92E-12 | 122 | Cdkn1b | 1.240475759 | 4.40E-02 |
| 123 | Gstm2 | 8.06201375 | 6.70085E-08 | 123 | Mthfs | 1.200898193 | 2.59E-02 |
| 124 | Psd3 | 8.049015334 | 4.38609E-13 | 124 | Ramp1 | 1.104854396 | 2.83E-02 |
| 125 | Gm14636 | 8.037213879 | 1.96733E-15 | 125 | Vgll4 | 1.03755019 | 4.20E-02 |
| 126 | Mfsd7a | 8.006081016 | 1.81173E-07 | 126 | | | |

| | | | | |
|---|---|---|---|---|
| 127 | Il1b | 7.999541099 | 5.58818E-10 | 127 |
| 128 | Tmem119 | 7.9874294 | 2.1291E-10 | 128 |
| 129 | Inhba | 7.987178347 | 1.20E-18 | 129 |
| 130 | Mmp13 | 7.984908542 | 2.28E-08 | 130 |
| 131 | Ace | 7.969652163 | 0.001098629 | 131 |
| 132 | Cxcl12 | 7.962761995 | 0.000645354 | 132 |
| 133 | Eps8 | 7.945734691 | 5.72E-38 | 133 |
| 134 | Ifi204 | 7.936155068 | 5.03348E-42 | 134 |
| 135 | Exoc3l4 | 7.929882273 | 1.28889E-07 | 135 |
| 136 | Oas1g | 7.922964028 | 1.00299E-11 | 136 |
| 137 | Htra3 | 7.879000432 | 1.39071E-07 | 137 |
| 138 | Ctf2 | 7.865820543 | 2.37E-07 | 138 |
| 139 | Nlrp1c-ps | 7.852094641 | 5.41864E-07 | 139 |
| 140 | Il7 | 7.846870123 | 3.02E-07 | 140 |
| 141 | Thbd | 7.844081563 | 2.143E-14 | 141 |
| 142 | Serping1 | 7.807064582 | 3.35E-07 | 142 |
| 143 | Cmklr1 | 7.799689855 | 2.96898E-17 | 143 |
| 144 | A4galt | 7.790288983 | 2.68346E-07 | 144 |
| 145 | Mrc1 | 7.769648115 | 3.66973E-61 | 145 |
| 146 | Cd63 | 7.761907328 | 2.65655E-65 | 146 |
| 147 | Msr1 | 7.756134436 | 5.72728E-09 | 147 |
| 148 | Capn6 | 7.728208887 | 4.90349E-07 | 148 |
| 149 | Mfap4 | 7.715271741 | 1.12256E-06 | 149 |
| 150 | Nfib | 7.707896543 | 9.15466E-10 | 150 |
| 151 | S100a16 | 7.686291281 | 1.31371E-11 | 151 |
| 152 | Tspan7 | 7.684710964 | 5.74318E-08 | 152 |
| 153 | Gm14461 | 7.682914011 | 1.84475E-06 | 153 |
| 154 | Mid1-ps1 | 7.680186326 | 3.36346E-06 | 154 |
| 155 | Copz2 | 7.679603276 | 1.79E-11 | 155 |
| 156 | Olr1 | 7.674116841 | 2.79724E-13 | 156 |
| 157 | Wwc1 | 7.669935755 | 2.77387E-06 | 157 |
| 158 | Pgf | 7.642809122 | 1.24548E-11 | 158 |

| | | | | |
|---|---|---|---|---|
| 159 | Mamdc2 | 7.629007346 | 3.61232E-06 | 159 |
| 160 | Ifi202b | 7.628162889 | 2.08324E-06 | 160 |
| 161 | Siglec1 | 7.627864632 | 1.76E-24 | 161 |
| 162 | Uaca | 7.626769189 | 7.14726E-11 | 162 |
| 163 | Gm44386 | 7.622870248 | 2.59695E-06 | 163 |
| 164 | Parva | 7.61823244 | 4.70E-11 | 164 |
| 165 | Tnfaip2 | 7.601930716 | 2.78E-42 | 165 |
| 166 | Dclk1 | 7.596741785 | 7.43E-09 | 166 |
| 167 | Saa3 | 7.591833256 | 2.49E-11 | 167 |
| 168 | Cped1 | 7.584230824 | 3.3366E-06 | 168 |
| 169 | Il33 | 7.561773296 | 2.83812E-14 | 169 |
| 170 | Pde7b | 7.533179051 | 3.57E-07 | 170 |
| 171 | Gm15247 | 7.522951709 | 6.75E-09 | 171 |
| 172 | Dpysl3 | 7.497763946 | 9.58943E-08 | 172 |
| 173 | Fcgr2b | 7.491545271 | 3.75731E-51 | 173 |
| 174 | Fhad1 | 7.449735086 | 1.37739E-05 | 174 |
| 175 | Gm20056 | 7.412383149 | 4.3689E-12 | 175 |
| 176 | Dram1 | 7.410740262 | 7.58563E-09 | 176 |
| 177 | Cp | 7.401988755 | 2.84843E-10 | 177 |
| 178 | Gm30329 | 7.399399901 | 7.53075E-06 | 178 |
| 179 | Gm43351 | 7.386702878 | 9.21725E-06 | 179 |
| 180 | Fxyd2 | 7.383695279 | 1.51139E-08 | 180 |
| 181 | Gm37726 | 7.381878209 | 9.81272E-08 | 181 |
| 182 | Rsad2 | 7.362809632 | 1.48708E-50 | 182 |
| 183 | Gm15340 | 7.35944337 | 1.1802E-05 | 183 |
| 184 | Abca9 | 7.341855272 | 8.64249E-27 | 184 |
| 185 | Nrp2 | 7.323030231 | 1.75339E-24 | 185 |
| 186 | Fkbp9 | 7.315121509 | 7.80E-07 | 186 |
| 187 | Cd276 | 7.292613831 | 3.76431E-11 | 187 |
| 188 | Mfap2 | 7.272465355 | 1.58775E-05 | 188 |
| 189 | Dhrs9 | 7.268869746 | 2.69794E-07 | 189 |
| 190 | F3 | 7.259625753 | 4.40E-05 | 190 |

| | | | | |
|---|---|---|---|---|
| 191 | Fcgr1 | 7.244847827 | 6.97E-26 | 191 |
| 192 | Fam20c | 7.241295947 | 2.70911E-08 | 192 |
| 193 | Emilin2 | 7.239902925 | 1.05495E-26 | 193 |
| 194 | C1ra | 7.214480026 | 4.49505E-07 | 194 |
| 195 | Gm6377 | 7.188008262 | 2.67114E-23 | 195 |
| 196 | Peg3 | 7.175151647 | 2.64224E-10 | 196 |
| 197 | Nupr1 | 7.173654351 | 2.55E-23 | 197 |
| 198 | Fbln2 | 7.170434103 | 8.65003E-05 | 198 |
| 199 | Zmynd15 | 7.161062609 | 5.85E-12 | 199 |
| 200 | Klhl13 | 7.16022305 | 0.000187615 | 200 |
| 201 | Stard8 | 7.158555492 | 5.2708E-40 | |
| 202 | Ms4a4a | 7.156539553 | 7.24726E-42 | |
| 203 | Epb41l1 | 7.155368243 | 1.5088E-08 | |
| 204 | Ms4a6d | 7.153225291 | 2.74477E-31 | |
| 205 | Vegfa | 7.152568684 | 3.21035E-11 | |
| 206 | Ankrd33b | 7.143023386 | 6.04732E-24 | |
| 207 | Aspn | 7.142739645 | 4.22814E-07 | |
| 208 | Cbr3 | 7.135602606 | 6.90937E-06 | |
| 209 | Cd300ld2 | 7.130731408 | 4.25645E-05 | |
| 210 | Mgl2 | 7.127927636 | 1.34575E-05 | |
| 211 | Dnm1 | 7.102997203 | 2.37611E-09 | |
| 212 | Ifi207 | 7.100936814 | 1.01238E-06 | |
| 213 | 9130230L23Rik | 7.096366711 | 2.38367E-06 | |
| 214 | Nlrp3 | 7.090046162 | 0.000460046 | |
| 215 | Map1b | 7.078722344 | 9.23505E-08 | |
| 216 | Cavin3 | 7.062140533 | 3.89303E-08 | |
| 217 | F830208F22Rik | 7.031776269 | 1.03079E-06 | |
| 218 | Hmox1 | 7.02363588 | 1.00562E-59 | |
| 219 | Iqsec2 | 7.013322903 | 1.70023E-05 | |
| 220 | Cd163 | 6.999304137 | 1.32608E-39 | |
| 221 | Alpk2 | 6.97532715 | 0.000467633 | |
| 222 | Pid1 | 6.950724267 | 2.23437E-29 | |

| | | | |
|---|---|---|---|
| 223 | Cygb | 6.939888765 | 4.07279E-09 |
| 224 | Lonrf3 | 6.934722421 | 1.32985E-05 |
| 225 | Mnda | 6.932250518 | 1.35288E-24 |
| 226 | Ogn | 6.928709241 | 0.043762458 |
| 227 | Fam124a | 6.920832588 | 0.000674481 |
| 228 | Procr | 6.919803169 | 3.14156E-17 |
| 229 | Tslp | 6.908653369 | 1.32987E-07 |
| 230 | F10 | 6.906555133 | 7.13835E-07 |
| 231 | Anxa3 | 6.905924927 | 9.08853E-32 |
| 232 | Ptprb | 6.878221667 | 3.42385E-10 |
| 233 | Clmp | 6.87662878 | 2.77387E-06 |
| 234 | Npy | 6.8681464 | 0.000781289 |
| 235 | Lamc2 | 6.865487707 | 9.74622E-07 |
| 236 | Rnd2 | 6.854375659 | 4.31582E-06 |
| 237 | Fam149a | 6.850608361 | 0.000369862 |
| 238 | Timp3 | 6.846023673 | 0.002144075 |
| 239 | Tlr2 | 6.8285302 | 1.18559E-43 |
| 240 | Klhl38 | 6.813983489 | 0.001106308 |
| 241 | Fmn1 | 6.811146507 | 9.43694E-15 |
| 242 | Ccl6 | 6.802103837 | 6.59632E-31 |
| 243 | Batf2 | 6.776547422 | 2.45605E-07 |
| 244 | Gab1 | 6.773979894 | 3.08579E-09 |
| 245 | Cd109 | 6.770225774 | 4.62191E-13 |
| 246 | Fbln5 | 6.738291121 | 6.12589E-08 |
| 247 | Apoe | 6.738104977 | 1.63847E-59 |
| 248 | Sgms2 | 6.730753643 | 4.51675E-14 |
| 249 | Hspa1l | 6.728721553 | 0.000630319 |
| 250 | Mest | 6.726165544 | 1.73104E-08 |
| 251 | Fblim1 | 6.715345441 | 4.25281E-11 |
| 252 | F7 | 6.710283895 | 0.001856589 |
| 253 | Ecscr | 6.693164527 | 1.91203E-05 |
| 254 | Gm5345 | 6.684715603 | 0.000276375 |

| | | | |
|---|---|---|---|
| 255 | Cttnbp2nl | 6.676586885 | 4.84898E-18 |
| 256 | Six1 | 6.660324293 | 0.001242535 |
| 257 | Rasal2 | 6.636642731 | 2.12984E-06 |
| 258 | Fcgrt | 6.633746023 | 3.84997E-37 |
| 259 | Rilp | 6.626716285 | 1.02079E-06 |
| 260 | Il5 | 6.614529912 | 0.004661966 |
| 261 | Ctgf | 6.601601658 | 8.41412E-12 |
| 262 | Gm14221 | 6.586908537 | 1.33593E-05 |
| 263 | Tnf | 6.575484752 | 3.37611E-37 |
| 264 | Fkbp10 | 6.559476282 | 3.49988E-06 |
| 265 | Etv1 | 6.547047744 | 3.90833E-09 |
| 266 | Serpinb9b | 6.536848439 | 0.006088279 |
| 267 | Sphk1 | 6.534987383 | 0.0022873 |
| 268 | Slc7a8 | 6.529281154 | 1.12125E-22 |
| 269 | Aoah | 6.516114303 | 1.81931E-22 |
| 270 | Snx24 | 6.51320255 | 7.22771E-06 |
| 271 | Rusc2 | 6.512598564 | 5.29364E-06 |
| 272 | Entpd3 | 6.503657562 | 7.5587E-05 |
| 273 | App | 6.492652451 | 4.57416E-38 |
| 274 | Cpne8 | 6.49095791 | 1.17189E-09 |
| 275 | E230013L22Rik | 6.490197188 | 3.35538E-05 |
| 276 | Egfl7 | 6.490172577 | 2.24193E-13 |
| 277 | Scamp5 | 6.485978001 | 2.20603E-22 |
| 278 | Hacd4 | 6.481910203 | 1.1822E-28 |
| 279 | Arhgap28 | 6.480457998 | 2.6071E-06 |
| 280 | Phldb1 | 6.47988355 | 8.59409E-13 |
| 281 | Plxna4os1 | 6.476980528 | 0.000314293 |
| 282 | Ugt1a7c | 6.471987628 | 3.85966E-07 |
| 283 | Mamld1 | 6.46143223 | 0.0003783 |
| 284 | Akr1b8 | 6.46099582 | 1.30427E-05 |
| 285 | Gm29340 | 6.45160168 | 9.05258E-06 |
| 286 | Fosl1 | 6.441433231 | 1.80431E-07 |

| | | | |
|---|---|---|---|
| 287 | Rhobtb1 | 6.437131463 | 1.35252E-07 |
| 288 | Rin2 | 6.435884279 | 8.50468E-31 |
| 289 | Cd300ld | 6.426113921 | 1.97898E-15 |
| 290 | Grhl2 | 6.421797188 | 0.004039723 |
| 291 | Fnip2 | 6.418757848 | 1.01846E-16 |
| 292 | Sprr2e | 6.411326246 | 0.0051567 |
| 293 | Phf11d | 6.397109601 | 5.80126E-25 |
| 294 | Fam198b | 6.386294727 | 0.000194498 |
| 295 | C1qb | 6.380454978 | 1.48675E-46 |
| 296 | Dpep2 | 6.378213447 | 4.42653E-18 |
| 297 | Ldb2 | 6.362437575 | 6.35853E-05 |
| 298 | Clec4a2 | 6.359817984 | 2.63027E-16 |
| 299 | Gm12031 | 6.342791129 | 3.48779E-05 |
| 300 | Gm7497 | 6.340190842 | 0.007038758 |
| 301 | Met | 6.336272548 | 0.038129144 |
| 302 | Cx3cr1 | 6.332121728 | 1.51207E-19 |
| 303 | Fkbp1b | 6.331286457 | 4.81245E-08 |
| 304 | Ptprm | 6.306421941 | 1.79047E-05 |
| 305 | Lpl | 6.299366831 | 1.43255E-45 |
| 306 | Rgl1 | 6.296018611 | 9.04128E-57 |
| 307 | Cavin2 | 6.277611927 | 6.0079E-09 |
| 308 | Vgf | 6.276441326 | 0.004895888 |
| 309 | Fgd4 | 6.272012287 | 5.5463E-06 |
| 310 | Slc6a8 | 6.265213189 | 1.21318E-10 |
| 311 | Adm | 6.250652742 | 5.76903E-12 |
| 312 | Kdelr3 | 6.234544436 | 6.43516E-05 |
| 313 | Adgrg6 | 6.223753655 | 3.41015E-06 |
| 314 | Clec14a | 6.215588202 | 0.039468399 |
| 315 | Adora2b | 6.18217263 | 2.67635E-06 |
| 316 | Best1 | 6.169585297 | 1.74031E-05 |
| 317 | Hk2 | 6.161582075 | 1.38596E-32 |
| 318 | Trim47 | 6.145836488 | 1.10671E-12 |

| | | | |
|---|---|---|---|
| 319 | Cryba4 | 6.133408408 | 1.28186E-05 |
| 320 | Lrrc36 | 6.128461233 | 0.014855428 |
| 321 | B4galt6 | 6.116411477 | 1.16848E-07 |
| 322 | 2610203C22Rik | 6.110808117 | 0.015630081 |
| 323 | Nid1 | 6.097336045 | 4.94079E-08 |
| 324 | Ptges | 6.096811951 | 0.014181306 |
| 325 | AW822252 | 6.092511848 | 0.002074405 |
| 326 | Cdh13 | 6.080075916 | 4.91078E-08 |
| 327 | Fmo1 | 6.079996207 | 0.001093143 |
| 328 | Tmem37 | 6.062549363 | 1.35321E-19 |
| 329 | 4932438H23Rik | 6.046648646 | 4.11373E-05 |
| 330 | Tgfbi | 6.045627684 | 1.98152E-39 |
| 331 | Ifit3b | 6.044012842 | 1.72571E-27 |
| 332 | F11r | 6.042028604 | 3.51963E-10 |
| 333 | Pkdcc | 6.03355968 | 6.30129E-06 |
| 334 | Nos3 | 6.011232567 | 5.2712E-06 |
| 335 | Zfp9 | 6.000543882 | 0.023593232 |
| 336 | Mefv | 5.995028629 | 2.09658E-13 |
| 337 | Gpx3 | 5.9910549 | 2.68549E-16 |
| 338 | Olfr433 | 5.989920723 | 0.001750347 |
| 339 | Igfbp6 | 5.985971933 | 0.000457192 |
| 340 | 2310043M15Rik | 5.975971572 | 0.001195752 |
| 341 | Fgd5 | 5.975589475 | 4.51587E-05 |
| 342 | Lyz2 | 5.966710074 | 4.00748E-58 |
| 343 | Shroom4 | 5.962755153 | 1.16345E-05 |
| 344 | Vsig4 | 5.961789015 | 0.004991649 |
| 345 | Rab3il1 | 5.949987327 | 7.04061E-38 |
| 346 | Plat | 5.94007676 | 2.27613E-05 |
| 347 | Camk1 | 5.93542893 | 2.07332E-20 |
| 348 | Timp1 | 5.930447089 | 1.44691E-05 |
| 349 | C1qc | 5.929474495 | 1.50233E-46 |
| 350 | Il12b | 5.917224077 | 0.013115461 |

| | | | |
|---|---|---|---|
| 351 | Gpr176 | 5.913068381 | 0.034626251 |
| 352 | Rnf150 | 5.893219514 | 6.96128E-06 |
| 353 | Myo1b | 5.892231659 | 3.63296E-05 |
| 354 | Hal | 5.892229903 | 0.000189214 |
| 355 | Rab11fip5 | 5.861709599 | 8.06538E-24 |
| 356 | Dmxl2 | 5.859705107 | 2.30021E-11 |
| 357 | Sulf2 | 5.85248218 | 2.67114E-23 |
| 358 | Maoa | 5.827137636 | 9.38342E-09 |
| 359 | Kcnk13 | 5.82045607 | 1.91184E-07 |
| 360 | Fndc7 | 5.8156847 | 3.38879E-06 |
| 361 | Gdpd1 | 5.811833843 | 5.95856E-05 |
| 362 | Tmem132a | 5.793428763 | 4.30543E-05 |
| 363 | Rnf144b | 5.789402854 | 4.08753E-07 |
| 364 | Hpgds | 5.777977083 | 3.10085E-37 |
| 365 | Oas2 | 5.774140238 | 2.68796E-14 |
| 366 | Gm37800 | 5.772359945 | 0.04790918 |
| 367 | Ccl22 | 5.768064474 | 0.007389056 |
| 368 | Mx2 | 5.768045271 | 2.44699E-19 |
| 369 | Slc35f5 | 5.767143124 | 1.09478E-19 |
| 370 | C1qa | 5.764879608 | 3.72857E-32 |
| 371 | Fn1 | 5.74656788 | 7.13404E-32 |
| 372 | Pla2g7 | 5.728191252 | 3.46962E-25 |
| 373 | Fcgr3 | 5.72267042 | 1.88079E-40 |
| 374 | Mill2 | 5.722628384 | 0.003515218 |
| 375 | Dusp8 | 5.722320887 | 0.000286849 |
| 376 | Mid1 | 5.719311492 | 8.66558E-22 |
| 377 | Msrb3 | 5.712303071 | 1.85668E-08 |
| 378 | Esam | 5.70385608 | 1.93692E-07 |
| 379 | Marcksl1 | 5.700351434 | 3.99521E-36 |
| 380 | Mafb | 5.688224014 | 1.25236E-35 |
| 381 | Fam13a | 5.67861444 | 0.02111573 |
| 382 | Tmem202 | 5.676452299 | 7.27853E-06 |

| | | | |
|---|---|---|---|
| 383 | Gm10616 | 5.670289928 | 0.011687239 |
| 384 | Rbm47 | 5.648779624 | 2.39901E-20 |
| 385 | Bhlhe41 | 5.646093021 | 1.63475E-11 |
| 386 | Csf3r | 5.641568272 | 7.32017E-14 |
| 387 | Klf4 | 5.617143903 | 1.79949E-10 |
| 388 | 9930022D16Rik | 5.605612061 | 1.57593E-07 |
| 389 | Cd36 | 5.602746742 | 4.55727E-32 |
| 390 | Adgre1 | 5.595383388 | 1.11047E-51 |
| 391 | Ltc4s | 5.594773148 | 8.08444E-06 |
| 392 | Tnfsf12 | 5.587336333 | 0.002606164 |
| 393 | Snx8 | 5.58558843 | 1.0599E-20 |
| 394 | Cd93 | 5.581328628 | 1.8299E-23 |
| 395 | Rab20 | 5.57544074 | 2.18325E-14 |
| 396 | 4930512H18Rik | 5.566712312 | 0.00031174 |
| 397 | Dock1 | 5.552042804 | 3.89303E-08 |
| 398 | Clec4a3 | 5.540156234 | 1.98262E-25 |
| 399 | Cyp11a1 | 5.537485879 | 1.59814E-06 |
| 400 | Gm4951 | 5.51940658 | 1.53437E-07 |
| 401 | Igf1 | 5.510069146 | 1.25236E-35 |
| 402 | Ccl24 | 5.500301915 | 3.0517E-12 |
| 403 | Pdgfc | 5.497469784 | 3.39042E-12 |
| 404 | Gm17276 | 5.49586413 | 0.039148362 |
| 405 | Cd63-ps | 5.49586413 | 0.039148362 |
| 406 | Rgs7bp | 5.492864554 | 0.002583506 |
| 407 | Trpv4 | 5.48713098 | 5.17428E-07 |
| 408 | Tie1 | 5.486138088 | 2.10408E-05 |
| 409 | Ifitm2 | 5.481592637 | 7.57667E-39 |
| 410 | Gm37787 | 5.468791156 | 0.000180171 |
| 411 | Syngr1 | 5.464030579 | 2.86767E-10 |
| 412 | Tinagl1 | 5.453104605 | 3.29312E-06 |
| 413 | Slc35e4 | 5.453045187 | 7.09832E-06 |
| 414 | Mtus1 | 5.449744868 | 2.5211E-08 |

| | | | |
|---|---|---|---|
| 415 | Hebp1 | 5.449057565 | 0.001548137 |
| 416 | Hk3 | 5.436107257 | 2.56774E-15 |
| 417 | Fosb | 5.435241746 | 7.74176E-17 |
| 418 | Lbp | 5.41862607 | 2.8526E-09 |
| 419 | Tln2 | 5.418152104 | 9.34526E-09 |
| 420 | Mx1 | 5.396212795 | 1.71008E-05 |
| 421 | Tlr13 | 5.388823208 | 9.75027E-06 |
| 422 | Ptpro | 5.361028601 | 7.16889E-13 |
| 423 | Gm13822 | 5.358910342 | 0.001345781 |
| 424 | Lgmn | 5.350517506 | 2.16648E-45 |
| 425 | Tek | 5.349572678 | 2.99119E-09 |
| 426 | Tppp3 | 5.346464029 | 0.003322614 |
| 427 | Plxna2 | 5.344802882 | 2.53137E-05 |
| 428 | Hcar2 | 5.344321922 | 7.4182E-05 |
| 429 | Cd300c2 | 5.343570714 | 5.11949E-23 |
| 430 | Afap1l1 | 5.340682817 | 4.81245E-08 |
| 431 | Tnfrsf11a | 5.334913213 | 8.18898E-11 |
| 432 | Cd68 | 5.334109223 | 2.9438E-31 |
| 433 | Cgref1 | 5.332216443 | 0.001177623 |
| 434 | Fnbp1l | 5.326061741 | 1.59842E-07 |
| 435 | Rbp1 | 5.315782324 | 0.000324591 |
| 436 | Pcolce | 5.30964255 | 1.01584E-08 |
| 437 | Tubb6 | 5.306117626 | 1.5408E-21 |
| 438 | Ifit3 | 5.298053704 | 1.78588E-24 |
| 439 | Csf1r | 5.280089247 | 3.33587E-42 |
| 440 | Sema6a | 5.266860696 | 0.000863912 |
| 441 | Ifitm3 | 5.262631404 | 2.98147E-25 |
| 442 | Cryab | 5.257548356 | 0.00399426 |
| 443 | Mcam | 5.23729125 | 2.07418E-09 |
| 444 | Ctsk | 5.235922793 | 2.48265E-06 |
| 445 | Tfec | 5.214502356 | 1.24479E-09 |
| 446 | Cdh5 | 5.213145967 | 1.73402E-07 |

| | | | |
|---|---|---|---|
| 447 | Plpp3 | 5.20940387 | 8.08831E-08 |
| 448 | Trim30c | 5.209390371 | 0.001397839 |
| 449 | Ehd2 | 5.198941431 | 2.56173E-10 |
| 450 | Dusp3 | 5.182937385 | 2.16872E-13 |
| 451 | Clec4a1 | 5.182861991 | 6.56887E-17 |
| 452 | Depdc7 | 5.163238744 | 6.96291E-05 |
| 453 | Rab31 | 5.151265956 | 9.18102E-27 |
| 454 | Cd70 | 5.149431919 | 0.004351969 |
| 455 | C77080 | 5.147310485 | 8.24132E-06 |
| 456 | Olfml3 | 5.140910176 | 2.28858E-08 |
| 457 | Clec4n | 5.120564296 | 1.32465E-20 |
| 458 | Col8a2 | 5.115202621 | 0.043710796 |
| 459 | Fam71f2 | 5.112133217 | 6.16234E-05 |
| 460 | Bmp2 | 5.096309357 | 8.19986E-12 |
| 461 | Gprc5c | 5.096063778 | 2.42424E-10 |
| 462 | Tnfrsf23 | 5.086784102 | 4.04651E-07 |
| 463 | Gatm | 5.064259978 | 1.66058E-14 |
| 464 | Ctnnd1 | 5.05193246 | 1.15254E-06 |
| 465 | Adcy9 | 5.051164037 | 0.001030839 |
| 466 | Grb10 | 5.039691232 | 2.50817E-05 |
| 467 | Isg15 | 5.021961862 | 5.40777E-23 |
| 468 | Papss2 | 5.014748787 | 0.000125764 |
| 469 | Parvb | 5.004783192 | 0.00010289 |
| 470 | Slc22a4 | 5.002833906 | 6.72233E-05 |
| 471 | Ptafr | 4.99231161 | 4.43745E-21 |
| 472 | Pik3r6 | 4.986875074 | 6.54785E-06 |
| 473 | Cd40 | 4.984717358 | 1.10293E-13 |
| 474 | Acta1 | 4.971630343 | 9.43839E-06 |
| 475 | Hist1h2be | 4.968703665 | 5.80996E-13 |
| 476 | Igsf6 | 4.957845695 | 4.85318E-25 |
| 477 | Aqp1 | 4.952208009 | 0.000185666 |
| 478 | Ramp2 | 4.950175633 | 0.001120338 |

| | | | |
|---|---|---|---|
| 479 | Lrrc25 | 4.945797318 | 5.76762E-11 |
| 480 | Cd300lg | 4.92619798 | 0.019091526 |
| 481 | C5ar2 | 4.921153677 | 0.031943296 |
| 482 | Slc16a7 | 4.918289417 | 0.001141132 |
| 483 | Rasgef1b | 4.913675913 | 2.23684E-38 |
| 484 | Plek | 4.913149939 | 4.73461E-25 |
| 485 | Hist3h2a | 4.911821491 | 2.75103E-18 |
| 486 | Tmem106a | 4.909670358 | 6.13878E-13 |
| 487 | Gm43775 | 4.9087723 | 0.000257801 |
| 488 | Hnmt | 4.906761711 | 0.00018435 |
| 489 | Gm26888 | 4.900533268 | 0.001602511 |
| 490 | Nlrc4 | 4.89535717 | 0.001399932 |
| 491 | Gbgt1 | 4.884171099 | 0.012450855 |
| 492 | Oit3 | 4.883773429 | 0.017618577 |
| 493 | Dusp18 | 4.882771827 | 1.75348E-05 |
| 494 | Serpinh1 | 4.879996954 | 0.000169884 |
| 495 | Slpi | 4.864158947 | 2.4642E-19 |
| 496 | C3 | 4.861665088 | 5.47667E-16 |
| 497 | Adap2os | 4.852306541 | 1.02731E-06 |
| 498 | Cd300lb | 4.849595192 | 2.96703E-10 |
| 499 | Fzd7 | 4.848518236 | 0.000770225 |
| 500 | Kitl | 4.843798089 | 0.000352537 |
| 501 | Rhob | 4.833045998 | 8.29513E-15 |
| 502 | Efemp2 | 4.820673777 | 3.87246E-05 |
| 503 | Pyroxd2 | 4.819652988 | 2.96981E-06 |
| 504 | Lgals3 | 4.812433075 | 9.57583E-27 |
| 505 | Jdp2 | 4.803811793 | 9.32101E-06 |
| 506 | Slc7a7 | 4.80355857 | 1.2357E-12 |
| 507 | Tspan33 | 4.802288535 | 0.000397758 |
| 508 | Igf2bp2 | 4.790029233 | 0.004007169 |
| 509 | Cavin1 | 4.788905384 | 9.01419E-10 |
| 510 | Oas1a | 4.776405104 | 2.82564E-09 |

| | | | |
|---|---|---|---|
| 511 | Ociad2 | 4.756991 | 0.013227335 |
| 512 | Cfap45 | 4.7462089 | 0.000172696 |
| 513 | Sorbs3 | 4.742967942 | 1.10453E-06 |
| 514 | Fabp5 | 4.74209741 | 4.76557E-28 |
| 515 | Tnip3 | 4.740990786 | 3.07132E-09 |
| 516 | Adora3 | 4.72848647 | 0.01892556 |
| 517 | Tmem267 | 4.722307949 | 0.033277585 |
| 518 | Itgam | 4.718967003 | 2.09658E-13 |
| 519 | Disc1 | 4.715245927 | 8.09837E-05 |
| 520 | Vwf | 4.7033581 | 2.5211E-08 |
| 521 | Csf2ra | 4.697500478 | 5.96402E-15 |
| 522 | Cebpd | 4.689175576 | 1.14456E-06 |
| 523 | Snx7 | 4.686262725 | 0.000234273 |
| 524 | Mir22hg | 4.683851448 | 1.13656E-14 |
| 525 | Lrp1 | 4.683413743 | 1.46712E-14 |
| 526 | Cmpk2 | 4.676583817 | 0.000180434 |
| 527 | Slc16a9 | 4.675853077 | 0.004000797 |
| 528 | Tlr3 | 4.671060775 | 0.000842042 |
| 529 | Lrp12 | 4.667406818 | 4.49991E-08 |
| 530 | Ang | 4.66188189 | 4.45797E-05 |
| 531 | Ifih1 | 4.661878256 | 7.67075E-20 |
| 532 | Btbd3 | 4.660735866 | 0.000995612 |
| 533 | Zc2hc1a | 4.66059676 | 0.000161128 |
| 534 | Plekhg3 | 4.659033841 | 5.78589E-06 |
| 535 | Rasa4 | 4.638608848 | 1.42275E-15 |
| 536 | Spire1 | 4.636582735 | 1.38002E-05 |
| 537 | Pln | 4.628735377 | 0.017442926 |
| 538 | Pla2g15 | 4.628124383 | 3.39042E-12 |
| 539 | Tspan12 | 4.627044221 | 0.024596777 |
| 540 | Cd209b | 4.624990524 | 0.030564787 |
| 541 | Cav2 | 4.615520142 | 2.15423E-09 |
| 542 | Fer | 4.613207984 | 3.25537E-09 |

| | | | |
|---|---|---|---|
| 543 | Gfpt2 | 4.6095766 | 0.003322614 |
| 544 | Hlx | 4.60694683 | 2.15636E-06 |
| 545 | Gm37648 | 4.603903189 | 0.006752874 |
| 546 | Gm5844 | 4.603454853 | 9.11543E-05 |
| 547 | Maff | 4.602760729 | 6.13157E-06 |
| 548 | Ninj1 | 4.585012515 | 4.31486E-25 |
| 549 | Abi3bp | 4.578320661 | 0.001247325 |
| 550 | Liph | 4.577381125 | 0.001772165 |
| 551 | Abca1 | 4.5764321 | 1.11572E-19 |
| 552 | A930007I19Rik | 4.576314987 | 8.09396E-05 |
| 553 | Gm5431 | 4.575926422 | 9.31425E-05 |
| 554 | Gm17024 | 4.575894234 | 0.002844695 |
| 555 | Fcgr4 | 4.575702388 | 4.52962E-06 |
| 556 | Oasl2 | 4.565810413 | 5.76903E-12 |
| 557 | P2ry6 | 4.563828382 | 4.43527E-12 |
| 558 | Gpr137b | 4.554927683 | 5.88524E-21 |
| 559 | Cebpa | 4.550987466 | 8.39253E-05 |
| 560 | Bst1 | 4.548609731 | 3.10742E-07 |
| 561 | Trp53inp2 | 4.545366479 | 3.36802E-12 |
| 562 | Lyz1 | 4.540392688 | 8.78008E-11 |
| 563 | Slfn4 | 4.524512297 | 1.57726E-08 |
| 564 | Armcx6 | 4.520015599 | 2.71929E-05 |
| 565 | Hyal1 | 4.516227014 | 4.99944E-11 |
| 566 | Sestd1 | 4.507886561 | 2.84826E-08 |
| 567 | Tmem140 | 4.50073443 | 1.10147E-14 |
| 568 | Gm8818 | 4.494811505 | 1.59842E-07 |
| 569 | Sult1a1 | 4.488727534 | 6.6458E-08 |
| 570 | Selenbp1 | 4.486294435 | 2.91255E-09 |
| 571 | Arhgap23 | 4.485766669 | 0.003055807 |
| 572 | Auts2 | 4.484034069 | 0.016150294 |
| 573 | Gamt | 4.482877894 | 0.00076721 |
| 574 | Cdc42ep1 | 4.471205924 | 0.009034791 |

| | | | |
|---|---|---|---|
| 575 | Renbp | 4.469297564 | 2.14094E-13 |
| 576 | Ltbr | 4.469106512 | 2.76131E-06 |
| 577 | Tbxas1 | 4.456404375 | 3.07524E-13 |
| 578 | Ccdc80 | 4.453979736 | 0.000145913 |
| 579 | Sdc1 | 4.453780107 | 0.009850329 |
| 580 | Adgrl4 | 4.450473796 | 1.20542E-06 |
| 581 | Plet1 | 4.447017918 | 0.011867349 |
| 582 | Hdc | 4.434972736 | 2.07483E-06 |
| 583 | Bach1 | 4.419975314 | 5.95445E-21 |
| 584 | Cd302 | 4.412255972 | 2.37628E-13 |
| 585 | Fermt2 | 4.408764754 | 0.001710577 |
| 586 | Gas2l1 | 4.391073623 | 2.31358E-10 |
| 587 | Ak1 | 4.387133048 | 0.015995116 |
| 588 | Optn | 4.383425618 | 1.00849E-08 |
| 589 | Clec7a | 4.379764779 | 3.78526E-07 |
| 590 | Tns3 | 4.378802525 | 7.63578E-10 |
| 591 | Adamts2 | 4.376578227 | 0.043664581 |
| 592 | Fcer1g | 4.372121695 | 1.1822E-28 |
| 593 | Anxa1 | 4.346983506 | 1.50814E-26 |
| 594 | Spi1 | 4.343843919 | 2.50033E-12 |
| 595 | Kctd4 | 4.336882255 | 0.003107351 |
| 596 | Aebp1 | 4.333149259 | 0.003240077 |
| 597 | Acacb | 4.314978079 | 0.003247251 |
| 598 | Rai14 | 4.314755278 | 3.46974E-05 |
| 599 | BC028528 | 4.313895684 | 3.32776E-13 |
| 600 | Wdfy3 | 4.287400884 | 9.79332E-13 |
| 601 | Acta2 | 4.286401692 | 0.005172381 |
| 602 | Basp1 | 4.282182842 | 6.05808E-05 |
| 603 | Gm14052 | 4.281495636 | 0.00024682 |
| 604 | Gm42725 | 4.279212835 | 8.50356E-10 |
| 605 | Pdgfb | 4.270023316 | 0.001753809 |
| 606 | Tanc2 | 4.268977973 | 3.16346E-09 |

| | | | |
|---|---|---|---|
| 607 | Hfe | 4.257832262 | 1.11366E-09 |
| 608 | Slamf8 | 4.242114864 | 1.00149E-06 |
| 609 | Aif1 | 4.23108557 | 9.66242E-17 |
| 610 | Fgfr1 | 4.229323738 | 1.77537E-09 |
| 611 | Zeb2 | 4.226451001 | 1.78346E-14 |
| 612 | Map3k6 | 4.218870425 | 0.004238277 |
| 613 | Htra1 | 4.217561521 | 0.017606781 |
| 614 | Ncf2 | 4.216700514 | 3.30292E-18 |
| 615 | Selenop | 4.214368115 | 7.96977E-32 |
| 616 | Ifi203-ps | 4.212213965 | 0.000521781 |
| 617 | Tmem88 | 4.203523258 | 0.002677986 |
| 618 | Tgm2 | 4.199043779 | 9.98313E-13 |
| 619 | Pald1 | 4.197267635 | 0.039393833 |
| 620 | Gm14023 | 4.196547629 | 0.010578739 |
| 621 | Tceal8 | 4.193905102 | 4.79648E-08 |
| 622 | Muc1 | 4.1914845 | 0.002119311 |
| 623 | Ttyh2 | 4.190003576 | 9.28427E-07 |
| 624 | Trf | 4.186706189 | 2.05404E-18 |
| 625 | Selenom | 4.181699002 | 0.000516337 |
| 626 | Gprc5a | 4.171480747 | 0.000152507 |
| 627 | Ms4a6c | 4.156326412 | 2.18952E-19 |
| 628 | M1ap | 4.153322293 | 0.020514196 |
| 629 | Lacc1 | 4.153018483 | 3.60528E-08 |
| 630 | Zfp385a | 4.148572447 | 0.000661121 |
| 631 | Enpep | 4.148270022 | 0.00044213 |
| 632 | Gpr84 | 4.138107831 | 0.007198299 |
| 633 | Mlkl | 4.136463936 | 1.17085E-05 |
| 634 | Coprs | 4.119163755 | 0.013672769 |
| 635 | Zfp703 | 4.11805587 | 1.32086E-07 |
| 636 | Adap2 | 4.11327297 | 6.20102E-10 |
| 637 | Tmcc3 | 4.106144095 | 1.56017E-05 |
| 638 | Aph1c | 4.092914696 | 0.000337535 |

| | | | |
|---|---|---|---|
| 639 | Ltbp4 | 4.090725102 | 0.000303144 |
| 640 | Sash1 | 4.08654405 | 5.75072E-06 |
| 641 | Tnfrsf12a | 4.086304474 | 0.001136181 |
| 642 | Clic4 | 4.081027948 | 3.7855E-18 |
| 643 | Dpt | 4.077376846 | 0.009896676 |
| 644 | Alpk1 | 4.05287853 | 2.67471E-06 |
| 645 | Zfand2a | 4.037288179 | 7.8581E-16 |
| 646 | Il13ra1 | 4.033144049 | 4.17595E-09 |
| 647 | Gpr157 | 4.032518099 | 0.000158238 |
| 648 | Mgll | 4.028995751 | 0.002607564 |
| 649 | Kctd12 | 4.023331308 | 1.0532E-10 |
| 650 | Arhgap29 | 4.021427814 | 1.22975E-05 |
| 651 | Wwc2 | 4.018781784 | 0.001362397 |
| 652 | Ushbp1 | 4.012661747 | 0.00059345 |
| 653 | Tbc1d24 | 4.012094253 | 3.27291E-05 |
| 654 | Lamb2 | 4.010453337 | 0.000695527 |
| 655 | Adam15 | 4.00443582 | 2.24193E-13 |
| 656 | F630028O10Rik | 3.998517023 | 0.000158108 |
| 657 | H2-M2 | 3.99784001 | 0.008967562 |
| 658 | Rcn3 | 3.995199867 | 4.8813E-06 |
| 659 | Ctsc | 3.990620472 | 1.86909E-24 |
| 660 | Stx3 | 3.985890209 | 1.03079E-06 |
| 661 | Ppbp | 3.981657892 | 2.51476E-05 |
| 662 | A930037H05Rik | 3.976406664 | 0.00033946 |
| 663 | Ropn1l | 3.972555894 | 0.039983305 |
| 664 | Fam213b | 3.971255743 | 2.26968E-09 |
| 665 | Flvcr2 | 3.970715126 | 0.023473692 |
| 666 | Itgb5 | 3.964002437 | 0.016043329 |
| 667 | Pkn3 | 3.962203464 | 0.003975768 |
| 668 | Slc30a1 | 3.961147743 | 4.22369E-06 |
| 669 | Tlr4 | 3.950315224 | 3.86208E-08 |
| 670 | Kdr | 3.946730399 | 1.76166E-06 |

| | | | |
|---|---|---|---|
| 671 | Lrg1 | 3.945856575 | 0.000748037 |
| 672 | Agap1 | 3.934750376 | 0.000603248 |
| 673 | Svip | 3.930047394 | 0.002332826 |
| 674 | Ppic | 3.929372003 | 7.01568E-07 |
| 675 | Epor | 3.923853056 | 0.024230401 |
| 676 | Stom | 3.91676681 | 2.89375E-15 |
| 677 | 2810025M15Rik | 3.913992906 | 0.000597563 |
| 678 | Nostrin | 3.907769988 | 0.001469715 |
| 679 | Enpp2 | 3.904788369 | 0.00024682 |
| 680 | Abcd2 | 3.894830751 | 0.004524754 |
| 681 | Gch1 | 3.894113628 | 5.90817E-12 |
| 682 | Slc11a1 | 3.893000699 | 1.35119E-12 |
| 683 | Osbpl1a | 3.891640022 | 4.27294E-05 |
| 684 | Gm26522 | 3.890310749 | 0.000125672 |
| 685 | Deptor | 3.889260987 | 1.49301E-09 |
| 686 | Steap4 | 3.882837769 | 0.008396013 |
| 687 | Galc | 3.881122893 | 5.71612E-06 |
| 688 | Dusp16 | 3.880051711 | 8.9432E-09 |
| 689 | Ubtd1 | 3.877213555 | 7.08356E-07 |
| 690 | Rassf4 | 3.874804954 | 1.39483E-19 |
| 691 | Qpct | 3.869337736 | 5.28887E-06 |
| 692 | Wwp1 | 3.861502622 | 3.98303E-25 |
| 693 | Hgsnat | 3.846067193 | 2.27565E-19 |
| 694 | Rab32 | 3.844243446 | 7.90012E-14 |
| 695 | Gm13571 | 3.841689056 | 0.001490203 |
| 696 | Tgfb1i1 | 3.841662977 | 0.025472714 |
| 697 | Stard9 | 3.83775815 | 3.88814E-07 |
| 698 | Fndc3b | 3.836378843 | 4.64906E-08 |
| 699 | Eln | 3.832576649 | 0.034868035 |
| 700 | Glis3 | 3.814790843 | 0.044914504 |
| 701 | Hmcn2 | 3.804754784 | 0.04326918 |
| 702 | Tdrd7 | 3.801786423 | 5.42165E-10 |

| | | | |
|---|---|---|---|
| 703 | Lst1 | 3.794841986 | 3.87161E-08 |
| 704 | Gm12164 | 3.79012749 | 0.001907719 |
| 705 | Fam114a1 | 3.789723738 | 5.49038E-05 |
| 706 | Plxnb2 | 3.788439287 | 2.41835E-10 |
| 707 | Lpcat2 | 3.787562766 | 2.07489E-06 |
| 708 | Ccl1 | 3.780500539 | 0.010107425 |
| 709 | Frmd4a | 3.773553112 | 1.30676E-08 |
| 710 | Ftl1 | 3.758990899 | 4.27988E-23 |
| 711 | Gpt2 | 3.755235149 | 6.46177E-05 |
| 712 | Fth-ps3 | 3.748904584 | 0.032127031 |
| 713 | Gm22774 | 3.746336672 | 0.000245959 |
| 714 | Col14a1 | 3.743696497 | 0.0048346 |
| 715 | Alox5ap | 3.738225945 | 1.01487E-11 |
| 716 | Slc2a6 | 3.721438294 | 9.76246E-08 |
| 717 | Spred1 | 3.720867655 | 2.3831E-05 |
| 718 | Pros1 | 3.720101241 | 1.51738E-12 |
| 719 | Sap30 | 3.718057119 | 6.32039E-09 |
| 720 | Lif | 3.717259105 | 0.000225312 |
| 721 | Fbxo30 | 3.716621149 | 3.82046E-13 |
| 722 | Jup | 3.708951444 | 1.74783E-09 |
| 723 | Marcks | 3.70665323 | 0.002912638 |
| 724 | Cacnb3 | 3.700554384 | 0.004052866 |
| 725 | Apbb2 | 3.695472646 | 0.025642898 |
| 726 | Tspan4 | 3.69337087 | 2.14416E-13 |
| 727 | Ticam2 | 3.681701851 | 0.005102512 |
| 728 | Dusp22 | 3.673967544 | 4.71095E-07 |
| 729 | Gab2 | 3.673799209 | 1.61739E-05 |
| 730 | B230216N24Rik | 3.671649933 | 0.013316466 |
| 731 | Atg4c | 3.669975493 | 4.45428E-05 |
| 732 | Ddx60 | 3.665740771 | 3.13641E-19 |
| 733 | Tlr5 | 3.661176879 | 0.042484281 |
| 734 | Serpine2 | 3.653334533 | 0.029448708 |

| | | | |
|---|---|---|---|
| 735 | Abcc3 | 3.646816892 | 8.38054E-10 |
| 736 | Gm37039 | 3.636965481 | 0.010950646 |
| 737 | Gm26857 | 3.629425067 | 0.001469715 |
| 738 | Ftl1-ps1 | 3.627917612 | 3.20945E-13 |
| 739 | Slc31a2 | 3.618639265 | 2.55213E-07 |
| 740 | Pirb | 3.617930206 | 1.60345E-15 |
| 741 | Scamp1 | 3.614987488 | 2.31206E-09 |
| 742 | Id1 | 3.608888293 | 0.011058947 |
| 743 | Tcp11l1 | 3.607956388 | 0.000989148 |
| 744 | Cpeb4 | 3.600982974 | 1.67012E-14 |
| 745 | Cpne2 | 3.593513099 | 3.13082E-05 |
| 746 | Plekhf1 | 3.592192789 | 0.024106971 |
| 747 | Cald1 | 3.591761255 | 0.000870919 |
| 748 | Gm45546 | 3.57872078 | 0.004276333 |
| 749 | Ccnyl1 | 3.576950077 | 0.024018468 |
| 750 | Src | 3.573098281 | 4.13386E-07 |
| 751 | Ctsf | 3.569482605 | 9.90435E-05 |
| 752 | Rcan1 | 3.567203445 | 2.71664E-08 |
| 753 | Stxbp6 | 3.566977559 | 0.005432803 |
| 754 | BC022687 | 3.558520925 | 0.014474536 |
| 755 | Gm42690 | 3.557699567 | 0.04783264 |
| 756 | Fth1 | 3.557238435 | 2.66143E-20 |
| 757 | Eng | 3.554838222 | 2.41836E-06 |
| 758 | Ppfibp2 | 3.554287768 | 3.55548E-06 |
| 759 | Spats2l | 3.554046692 | 0.04973702 |
| 760 | Slc9a3r2 | 3.549307696 | 0.002478271 |
| 761 | Ehd1 | 3.545693949 | 1.57208E-12 |
| 762 | Gm12346 | 3.544926373 | 1.04285E-07 |
| 763 | Ctsb | 3.543851295 | 6.97067E-26 |
| 764 | Pdlim4 | 3.538615267 | 5.72677E-11 |
| 765 | Gm6977 | 3.532751514 | 0.010426069 |
| 766 | Rhbdf1 | 3.531097325 | 0.005132697 |

| | | | |
|---|---|---|---|
| 767 | Scpep1 | 3.525390904 | 9.2231E-17 |
| 768 | Atp6v1a | 3.522838132 | 4.10233E-21 |
| 769 | 1500002C15Rik | 3.518176871 | 0.01095484 |
| 770 | Irak3 | 3.512339508 | 0.000239037 |
| 771 | P2ry12 | 3.510741655 | 3.48779E-05 |
| 772 | Tnfrsf21 | 3.51048139 | 7.06144E-06 |
| 773 | Zfhx3 | 3.50883397 | 0.001143785 |
| 774 | Osm | 3.505505218 | 1.95779E-11 |
| 775 | Hist1h1d | 3.505123847 | 0.006803205 |
| 776 | Gm15590 | 3.496696618 | 1.79541E-07 |
| 777 | Adgrl2 | 3.488381328 | 0.000125323 |
| 778 | Amz1 | 3.487606585 | 1.53426E-05 |
| 779 | Mcoln2 | 3.473920402 | 6.93203E-05 |
| 780 | Tbc1d32 | 3.466763304 | 0.001931025 |
| 781 | Anxa5 | 3.465688858 | 3.04103E-18 |
| 782 | Dnajb4 | 3.464328131 | 3.72709E-06 |
| 783 | Gm43323 | 3.464109456 | 0.039666465 |
| 784 | Plxna1 | 3.463229691 | 1.08767E-06 |
| 785 | Rasgrp3 | 3.462225608 | 6.19595E-05 |
| 786 | Pstpip2 | 3.458984718 | 0.00162209 |
| 787 | Tmem205 | 3.45668902 | 0.020474244 |
| 788 | Ralgds | 3.447543259 | 3.33461E-10 |
| 789 | Adarb1 | 3.44751594 | 0.010483335 |
| 790 | Nme4 | 3.44223786 | 0.015551552 |
| 791 | Gm6297 | 3.434550592 | 0.042075871 |
| 792 | Trim36 | 3.426264712 | 0.005282307 |
| 793 | Mfsd11 | 3.421307997 | 1.31386E-11 |
| 794 | Pla2g4a | 3.419566922 | 0.000451298 |
| 795 | Cd86 | 3.41909325 | 2.22E-13 |
| 796 | Nod2 | 3.415944277 | 0.010646526 |
| 797 | Tspan15 | 3.41092384 | 0.039073653 |
| 798 | Amotl1 | 3.410596546 | 0.002071248 |

| | | | |
|---|---|---|---|
| 799 | Lama5 | 3.407645134 | 0.011532406 |
| 800 | Plagl1 | 3.406870224 | 9.05892E-16 |
| 801 | Cd300a | 3.399165082 | 3.38687E-11 |
| 802 | Cdc42ep4 | 3.398481583 | 5.25438E-06 |
| 803 | Cstb | 3.393181958 | 2.27821E-12 |
| 804 | Grn | 3.392895658 | 5.74914E-19 |
| 805 | Tmem176b | 3.39202055 | 2.22433E-17 |
| 806 | C2 | 3.38720358 | 0.009640508 |
| 807 | Nectin4 | 3.385973083 | 0.004302506 |
| 808 | Pdxk | 3.382077472 | 1.62441E-15 |
| 809 | Gm7160 | 3.380929822 | 0.013068691 |
| 810 | Glul | 3.374930536 | 1.32465E-20 |
| 811 | Cmtm3 | 3.367835432 | 1.4404E-11 |
| 812 | Cst3 | 3.366023326 | 7.08733E-20 |
| 813 | Lhfp | 3.362788265 | 0.000978693 |
| 814 | Cd33 | 3.356982673 | 3.41464E-10 |
| 815 | Itsn1 | 3.352760552 | 1.30917E-09 |
| 816 | Zbtb21 | 3.34739765 | 4.81245E-08 |
| 817 | Arhgap22 | 3.344718579 | 0.012315596 |
| 818 | Atp6v0a1 | 3.340356715 | 2.42992E-11 |
| 819 | Rnf180 | 3.340006108 | 0.048145543 |
| 820 | 4931440P22Rik | 3.334335134 | 0.040651386 |
| 821 | Mertk | 3.321341653 | 8.60036E-07 |
| 822 | Anpep | 3.320190841 | 0.027325727 |
| 823 | Gstm1 | 3.313595819 | 6.41315E-07 |
| 824 | Slamf7 | 3.305286658 | 2.19268E-11 |
| 825 | Zfp516 | 3.304098588 | 0.000610086 |
| 826 | Plod1 | 3.302752499 | 5.69635E-08 |
| 827 | Cpq | 3.300149287 | 0.001648926 |
| 828 | Socs3 | 3.299703048 | 1.2486E-12 |
| 829 | Slc37a2 | 3.29469658 | 6.72349E-17 |
| 830 | Tubb4a | 3.292848441 | 0.006954718 |

| | | | |
|---|---|---|---|
| 831 | Rnf130 | 3.284495762 | 1.70214E-09 |
| 832 | Armcx3 | 3.284391373 | 7.06594E-06 |
| 833 | Ppp1r9a | 3.276268107 | 0.003414123 |
| 834 | Pdgfa | 3.267029756 | 0.004870409 |
| 835 | Sqstm1 | 3.266111871 | 1.06339E-18 |
| 836 | Ccl5 | 3.264059858 | 6.50615E-14 |
| 837 | Aifm2 | 3.262706778 | 0.00182243 |
| 838 | Ctbp2 | 3.25986169 | 0.003892041 |
| 839 | Col15a1 | 3.259444605 | 0.008682576 |
| 840 | Lima1 | 3.251419734 | 5.17092E-06 |
| 841 | Eml1 | 3.249208905 | 0.029961256 |
| 842 | Mir155hg | 3.249023828 | 5.12077E-06 |
| 843 | Cpe | 3.247901038 | 0.009035868 |
| 844 | Smagp | 3.240557861 | 0.013503996 |
| 845 | Card19 | 3.240095644 | 1.72691E-09 |
| 846 | Blvrb | 3.235780921 | 2.09658E-13 |
| 847 | Hist1h2ad | 3.231905519 | 0.019388133 |
| 848 | Ftl2 | 3.22313852 | 0.001023955 |
| 849 | Map3k20 | 3.218492247 | 0.01088059 |
| 850 | Mgst1 | 3.21489371 | 0.000138505 |
| 851 | Luzp1 | 3.206437676 | 5.39799E-06 |
| 852 | Gbp5 | 3.205646881 | 4.3677E-06 |
| 853 | Tbc1d9 | 3.202490427 | 0.013541712 |
| 854 | Ftl1-ps2 | 3.20045022 | 0.034153802 |
| 855 | B4galt5 | 3.200020543 | 3.57246E-07 |
| 856 | Litaf | 3.190440232 | 9.8742E-15 |
| 857 | Apobr | 3.190030034 | 0.000109289 |
| 858 | Pfkfb4 | 3.18616091 | 7.08912E-06 |
| 859 | Usp2 | 3.172103694 | 0.000185874 |
| 860 | Ssfa2 | 3.167574534 | 7.09969E-10 |
| 861 | Coq10b | 3.166805713 | 1.1519E-14 |
| 862 | Atp6v0c | 3.163264812 | 2.14642E-10 |

| | | | |
|---|---|---|---|
| 863 | Nectin2 | 3.16115248 | 0.018698878 |
| 864 | Nfix | 3.160489526 | 0.000169598 |
| 865 | Nat9 | 3.159935367 | 0.000621263 |
| 866 | Mitf | 3.156830978 | 0.001821399 |
| 867 | Arhgef40 | 3.138179505 | 0.022103001 |
| 868 | Zik1 | 3.136750299 | 0.024256683 |
| 869 | Afdn | 3.127099841 | 0.004657386 |
| 870 | Klf9 | 3.126723385 | 0.025642898 |
| 871 | Hexa | 3.121328795 | 8.25134E-18 |
| 872 | Nampt | 3.11992398 | 1.47441E-11 |
| 873 | Casp4 | 3.118851643 | 2.22961E-07 |
| 874 | Txnrd1 | 3.110484669 | 4.34388E-09 |
| 875 | Idh1 | 3.108194333 | 5.52725E-08 |
| 876 | Myo5a | 3.10684739 | 9.81205E-12 |
| 877 | Gm15472 | 3.100766331 | 0.033201943 |
| 878 | Ckap4 | 3.092054381 | 0.041532351 |
| 879 | Rtp4 | 3.086003946 | 5.52957E-12 |
| 880 | Gm26780 | 3.080064563 | 0.024537198 |
| 881 | Snx10 | 3.078791495 | 5.47943E-08 |
| 882 | Pak1 | 3.073956667 | 3.74956E-05 |
| 883 | 3110043O21Rik | 3.073309294 | 7.95364E-11 |
| 884 | Ptk2 | 3.07234519 | 0.00124512 |
| 885 | S100a1 | 3.064480578 | 7.95399E-05 |
| 886 | Sytl3 | 3.064445327 | 9.19637E-05 |
| 887 | Gpr137b-ps | 3.063722989 | 8.00239E-11 |
| 888 | Sgk1 | 3.063135764 | 1.31155E-16 |
| 889 | Gm7204 | 3.062552173 | 0.014570538 |
| 890 | Steap3 | 3.06096111 | 0.012078133 |
| 891 | Cfp | 3.060848878 | 1.45719E-06 |
| 892 | Ddah2 | 3.059645133 | 0.001981185 |
| 893 | Pkp4 | 3.056893357 | 2.65691E-05 |
| 894 | Acoxl | 3.052223242 | 0.006510237 |

| | | | |
|---|---|---|---|
| 895 | Plaur | 3.051840671 | 8.50617E-08 |
| 896 | Bag3 | 3.046955895 | 7.73661E-05 |
| 897 | Prdx1 | 3.036784676 | 1.04778E-14 |
| 898 | Spic | 3.034368344 | 2.77387E-06 |
| 899 | Acvrl1 | 3.034350386 | 0.0009021 |
| 900 | Mocos | 3.031375568 | 0.019430464 |
| 901 | Il1r1 | 3.01725173 | 0.004013892 |
| 902 | Rasgrp4 | 3.017036562 | 0.00043074 |
| 903 | Tmem176a | 3.005544806 | 7.66469E-09 |
| 904 | Itgb4 | 3.000189665 | 0.03749354 |
| 905 | Pi4k2a | 2.996463141 | 9.21725E-06 |
| 906 | 4921511C10Rik | 2.98633173 | 0.012188467 |
| 907 | Gpr34 | 2.983414318 | 1.26312E-09 |
| 908 | Nav1 | 2.982883176 | 0.000243626 |
| 909 | Ifi44 | 2.98125814 | 0.047121342 |
| 910 | Ehd4 | 2.977727311 | 1.69622E-15 |
| 911 | Tspan17 | 2.97268219 | 0.000840432 |
| 912 | Sqor | 2.968132845 | 0.000181562 |
| 913 | Zbtb10 | 2.96626283 | 0.000661121 |
| 914 | Gm22751 | 2.965330396 | 0.044547683 |
| 915 | Gca | 2.963013168 | 0.040774914 |
| 916 | Tcn2 | 2.960839717 | 4.77492E-07 |
| 917 | Naaa | 2.957787902 | 2.32934E-06 |
| 918 | 44256 | 2.957248089 | 0.007039349 |
| 919 | Npdc1 | 2.954193404 | 0.018486917 |
| 920 | Spdl1 | 2.953804108 | 0.004222568 |
| 921 | Bmf | 2.952517555 | 0.012109131 |
| 922 | Zfp667 | 2.951607718 | 0.013869013 |
| 923 | E330020D12Rik | 2.949545525 | 0.003293365 |
| 924 | Zfp36 | 2.943924948 | 2.42782E-14 |
| 925 | Wipi1 | 2.936587076 | 0.009179366 |
| 926 | Dhdh | 2.933658355 | 0.02780514 |

| | | | |
|---|---|---|---|
| 927 | Tnfrsf1a | 2.928338406 | 4.62191E-13 |
| 928 | Gm26603 | 2.91639708 | 0.024287651 |
| 929 | Csrp2 | 2.912103408 | 0.017421176 |
| 930 | Lamp1 | 2.909209587 | 9.18301E-12 |
| 931 | Plekho1 | 2.899859499 | 0.000136267 |
| 932 | Uap1l1 | 2.891890116 | 1.85056E-10 |
| 933 | Mxi1 | 2.888605475 | 0.000381829 |
| 934 | Syk | 2.882617394 | 2.0872E-08 |
| 935 | Tmem141 | 2.87890314 | 0.025463048 |
| 936 | Il18bp | 2.872636551 | 1.01842E-05 |
| 937 | Tns1 | 2.871820405 | 2.29861E-14 |
| 938 | Znfx1 | 2.870458492 | 3.05029E-09 |
| 939 | Asah1 | 2.854429168 | 1.84983E-13 |
| 940 | Entpd1 | 2.854265847 | 4.295E-14 |
| 941 | 2900093K20Rik | 2.850995886 | 0.038341799 |
| 942 | Abcb1a | 2.844532355 | 0.001412707 |
| 943 | Fem1c | 2.842908209 | 2.04557E-12 |
| 944 | Ctss | 2.839948656 | 5.48403E-15 |
| 945 | Gm37472 | 2.838519845 | 0.021987156 |
| 946 | Atp6v0d2 | 2.834979731 | 0.002745479 |
| 947 | Gm8355 | 2.832045584 | 0.024409449 |
| 948 | Dclre1c | 2.827032449 | 7.51443E-08 |
| 949 | 44263 | 2.824530291 | 0.007763534 |
| 950 | Ogfrl1 | 2.822258592 | 0.006897616 |
| 951 | Usp18 | 2.81547914 | 1.00781E-05 |
| 952 | Apobec1 | 2.813556217 | 9.67917E-09 |
| 953 | Dnase1l1 | 2.799947902 | 1.67637E-06 |
| 954 | Ly86 | 2.799524313 | 1.23434E-08 |
| 955 | Trim46 | 2.795683915 | 0.004600484 |
| 956 | Ly96 | 2.79496055 | 2.6837E-07 |
| 957 | Rgcc | 2.794096222 | 8.65033E-09 |
| 958 | Atp8b4 | 2.794079049 | 0.02378503 |

| | | | |
|---|---|---|---|
| 959 | Tmem159 | 2.793177296 | 0.026427213 |
| 960 | Zfand5 | 2.791534994 | 6.13878E-13 |
| 961 | Mpzl1 | 2.791437527 | 0.026216229 |
| 962 | Sdc3 | 2.78720037 | 0.000235264 |
| 963 | Fam212a | 2.783680239 | 0.004025309 |
| 964 | Sowahc | 2.7820008 | 0.010948119 |
| 965 | Lamp2 | 2.77559651 | 4.77125E-13 |
| 966 | Ifit1bl2 | 2.773524632 | 0.018516592 |
| 967 | Cd180 | 2.771376866 | 6.93491E-05 |
| 968 | Clcn5 | 2.770738978 | 8.68654E-06 |
| 969 | Nuak2 | 2.768013166 | 9.8817E-05 |
| 970 | Gm26762 | 2.767193474 | 0.004970483 |
| 971 | Rragd | 2.760810405 | 5.80159E-07 |
| 972 | Cpeb2 | 2.752602479 | 0.00074753 |
| 973 | Cask | 2.750060278 | 0.022039589 |
| 974 | Qk | 2.749424558 | 5.31285E-11 |
| 975 | Ctsd | 2.747231013 | 3.17829E-12 |
| 976 | Hpgd | 2.74335766 | 6.45195E-08 |
| 977 | Maged1 | 2.741219536 | 4.59341E-05 |
| 978 | Kifap3 | 2.739593947 | 0.001330255 |
| 979 | Milr1 | 2.730272461 | 0.000231337 |
| 980 | Serpinb6a | 2.729343275 | 8.04024E-11 |
| 981 | Trim25 | 2.722616088 | 2.32062E-11 |
| 982 | Trim16 | 2.721925344 | 0.03052354 |
| 983 | Tuft1 | 2.720038695 | 0.048439474 |
| 984 | Cysltr1 | 2.716593947 | 0.000120099 |
| 985 | Xdh | 2.711715837 | 7.95036E-09 |
| 986 | Timp2 | 2.711111978 | 4.09714E-11 |
| 987 | Rap2b | 2.709901869 | 4.11772E-06 |
| 988 | Spryd7 | 2.709283059 | 3.66389E-06 |
| 989 | Bcar3 | 2.707531345 | 0.000487307 |
| 990 | Dnajb1 | 2.700746347 | 1.4344E-08 |

| | | | |
|---|---|---|---|
| 991 | Fam214b | 2.693351362 | 2.80933E-07 |
| 992 | Il15ra | 2.686669652 | 3.99246E-05 |
| 993 | Arap3 | 2.685516295 | 0.025071098 |
| 994 | Homer3 | 2.682483656 | 0.003494619 |
| 995 | Wdr60 | 2.67973844 | 0.006823092 |
| 996 | Lonrf1 | 2.678514338 | 0.018141956 |
| 997 | Nfe2l2 | 2.666752095 | 7.65699E-11 |
| 998 | Pon3 | 2.666575464 | 0.000280464 |
| 999 | Pfkfb3 | 2.658521304 | 1.16757E-07 |
| 1000 | Gpr160 | 2.658441621 | 0.001446521 |
| 1001 | Mgst3 | 2.652473556 | 0.000845181 |
| 1002 | Efhd2 | 2.651698073 | 4.72707E-10 |
| 1003 | Cln8 | 2.645048527 | 5.86437E-06 |
| 1004 | Lzts2 | 2.644846358 | 0.012354326 |
| 1005 | Nagk | 2.637873079 | 4.59341E-05 |
| 1006 | Nqo1 | 2.635724657 | 0.03955922 |
| 1007 | Slc35f6 | 2.632726833 | 8.7804E-09 |
| 1008 | Cnrip1 | 2.617676141 | 0.024984846 |
| 1009 | Lat2 | 2.615309563 | 1.32969E-05 |
| 1010 | Fam43a | 2.613024709 | 0.000295136 |
| 1011 | Amdhd2 | 2.611298108 | 1.70268E-08 |
| 1012 | Slc16a10 | 2.608035733 | 0.009586179 |
| 1013 | Chpt1 | 2.607395095 | 0.023854265 |
| 1014 | Nacc2 | 2.604720559 | 0.001876265 |
| 1015 | Tceal9 | 2.594263736 | 1.4344E-08 |
| 1016 | Dpp7 | 2.58895905 | 7.97617E-07 |
| 1017 | Ggh | 2.58831863 | 6.70014E-05 |
| 1018 | Mfsd1 | 2.587988196 | 3.21189E-11 |
| 1019 | Nenf | 2.587217738 | 0.00258123 |
| 1020 | Cdc42bpa | 2.587171522 | 0.025333898 |
| 1021 | Gng12 | 2.58701449 | 9.16103E-06 |
| 1022 | Cdk14 | 2.585271862 | 0.024341598 |

| | | | |
|---|---|---|---|
| 1023 | Atp6v0b | 2.581942198 | 8.7828E-09 |
| 1024 | Ldlr | 2.577896054 | 0.001368413 |
| 1025 | Fam83g | 2.576815542 | 0.00877044 |
| 1026 | Arhgap10 | 2.572744846 | 0.000735977 |
| 1027 | Med12l | 2.570147434 | 0.024099802 |
| 1028 | Tpst1 | 2.557700589 | 0.000873799 |
| 1029 | Lrrk1 | 2.55649069 | 0.00058408 |
| 1030 | Sirpa | 2.554311895 | 1.63948E-09 |
| 1031 | Dtx4 | 2.548830797 | 0.037003799 |
| 1032 | Rybp | 2.547544897 | 0.030916658 |
| 1033 | Kif3a | 2.547289916 | 0.000146704 |
| 1034 | Lpin2 | 2.545571713 | 1.14456E-06 |
| 1035 | Ciart | 2.543393702 | 0.020852091 |
| 1036 | Fnip1 | 2.530159859 | 1.63526E-06 |
| 1037 | Per3 | 2.530145454 | 0.001214308 |
| 1038 | Coq8a | 2.529629652 | 1.29476E-07 |
| 1039 | Parp12 | 2.524380941 | 0.003515218 |
| 1040 | Mapk3 | 2.524094647 | 7.27527E-05 |
| 1041 | Ptgs1 | 2.52275189 | 0.004021455 |
| 1042 | Dhx58 | 2.520487931 | 4.66001E-08 |
| 1043 | Dusp14 | 2.519492429 | 0.001432479 |
| 1044 | D730003I15Rik | 2.517405531 | 0.00243106 |
| 1045 | Sele | 2.510402114 | 0.039315325 |
| 1046 | Atp6v0d1 | 2.509992563 | 9.90685E-10 |
| 1047 | Fhl1 | 2.508981013 | 0.034245468 |
| 1048 | Fcna | 2.50834437 | 6.13157E-06 |
| 1049 | Ripk2 | 2.505261479 | 4.76588E-05 |
| 1050 | N4bp1 | 2.50402017 | 1.32081E-05 |
| 1051 | Gm24245 | 2.503169953 | 0.021397013 |
| 1052 | Pld2 | 2.500984158 | 0.00192128 |
| 1053 | P4ha1 | 2.495395986 | 3.68644E-09 |
| 1054 | Rgs18 | 2.49458773 | 0.001855622 |

| | | | |
|---|---|---|---|
| 1055 | Serpinf1 | 2.492352746 | 0.012431753 |
| 1056 | Slc26a11 | 2.487745113 | 0.004986936 |
| 1057 | E2f1 | 2.484538161 | 0.011731619 |
| 1058 | Rabgef1 | 2.476759799 | 1.05055E-05 |
| 1059 | Ctsa | 2.476485122 | 2.64141E-12 |
| 1060 | Csf2rb | 2.47409801 | 5.3206E-05 |
| 1061 | Ap2a2 | 2.468755694 | 1.20598E-11 |
| 1062 | Cryzl2 | 2.464067775 | 0.044928266 |
| 1063 | Ctsz | 2.463371379 | 2.97346E-11 |
| 1064 | Pnp | 2.462385948 | 2.60611E-06 |
| 1065 | Nfia | 2.456305791 | 0.00188604 |
| 1066 | Cltb | 2.447682366 | 0.000542283 |
| 1067 | Tmem2 | 2.444687512 | 0.000231418 |
| 1068 | Eno2 | 2.442400818 | 0.006541384 |
| 1069 | Ocrl | 2.441706574 | 2.92238E-06 |
| 1070 | Mcfd2 | 2.438657452 | 4.78318E-05 |
| 1071 | 3110082I17Rik | 2.43582985 | 0.001668368 |
| 1072 | Mafg | 2.431944328 | 3.64948E-07 |
| 1073 | Daam1 | 2.431142326 | 0.002497648 |
| 1074 | Stxbp1 | 2.423927324 | 0.018207899 |
| 1075 | Cdkn1c | 2.40948849 | 0.008974377 |
| 1076 | Crip2 | 2.407474843 | 5.38059E-07 |
| 1077 | Csf2rb2 | 2.406952944 | 0.001844734 |
| 1078 | Cyp27a1 | 2.404068125 | 0.000165886 |
| 1079 | Myo7a | 2.403710659 | 0.000514829 |
| 1080 | Sdcbp | 2.402171793 | 1.70165E-10 |
| 1081 | Ptov1 | 2.393609308 | 0.002287887 |
| 1082 | Gda | 2.380550682 | 0.013895046 |
| 1083 | Mef2c | 2.379948622 | 2.85241E-06 |
| 1084 | Gcnt2 | 2.377619718 | 0.018327669 |
| 1085 | Lmo2 | 2.376509533 | 0.001169575 |
| 1086 | Ube2q2 | 2.375129298 | 0.000581671 |

| | | | |
|---|---|---|---|
| 1087 | Gm14305 | 2.370574849 | 0.033464318 |
| 1088 | Ubc | 2.366384138 | 1.63992E-10 |
| 1089 | 2310022A10Rik | 2.363426161 | 0.000224589 |
| 1090 | Nceh1 | 2.356072979 | 2.93859E-07 |
| 1091 | Mdfic | 2.353636231 | 3.55548E-06 |
| 1092 | Fndc3a | 2.352541083 | 6.90937E-06 |
| 1093 | Nt5dc2 | 2.342035982 | 0.041947052 |
| 1094 | Plekhn1 | 2.339485424 | 0.004969326 |
| 1095 | Dusp6 | 2.337841798 | 4.88827E-08 |
| 1096 | Fam102b | 2.335988719 | 0.00570128 |
| 1097 | P2rx4 | 2.333910885 | 1.05001E-09 |
| 1098 | Slc31a1 | 2.333509994 | 1.11017E-05 |
| 1099 | Hck | 2.320690515 | 9.42681E-05 |
| 1100 | Ip6k2 | 2.320330819 | 0.003526098 |
| 1101 | Anxa7 | 2.319491162 | 1.61952E-07 |
| 1102 | Icam1 | 2.308152068 | 1.12162E-07 |
| 1103 | Gm20594 | 2.305796584 | 0.009711568 |
| 1104 | Slc39a1 | 2.30387509 | 0.001637889 |
| 1105 | Lyn | 2.299898587 | 6.12589E-08 |
| 1106 | Cited2 | 2.29092623 | 5.67958E-05 |
| 1107 | Tyrobp | 2.288883902 | 1.19817E-08 |
| 1108 | Cd83 | 2.287393505 | 1.51139E-08 |
| 1109 | Mtss1 | 2.286822773 | 1.39071E-07 |
| 1110 | Ntpcr | 2.283571101 | 0.011375186 |
| 1111 | Ap1s2 | 2.281997264 | 7.08356E-07 |
| 1112 | Arrdc4 | 2.275122558 | 8.54844E-06 |
| 1113 | Wdsub1 | 2.271690331 | 0.003089739 |
| 1114 | Trps1 | 2.266534994 | 6.90978E-05 |
| 1115 | Homer1 | 2.265427171 | 0.016560707 |
| 1116 | Sod2 | 2.261837218 | 1.07071E-05 |
| 1117 | Sipa1l2 | 2.257891028 | 0.015444021 |
| 1118 | 1110032A03Rik | 2.255532907 | 0.006106993 |

| | | | |
|---|---|---|---|
| 1119 | Blnk | 2.253475219 | 2.54287E-05 |
| 1120 | Gm42418 | 2.252062993 | 2.67103E-07 |
| 1121 | Hist1h4c | 2.244925338 | 0.016150294 |
| 1122 | Wls | 2.24185111 | 4.12173E-07 |
| 1123 | Col4a3bp | 2.241615703 | 8.26831E-06 |
| 1124 | Fam26f | 2.232718681 | 0.039573676 |
| 1125 | Cass4 | 2.226828953 | 0.004562363 |
| 1126 | Cndp2 | 2.224332779 | 1.41541E-05 |
| 1127 | Snx16 | 2.219061317 | 0.00119969 |
| 1128 | Creg1 | 2.215135225 | 2.09912E-09 |
| 1129 | Tifa | 2.21472991 | 0.000442476 |
| 1130 | Hmgb1-ps5 | 2.21210043 | 0.036803709 |
| 1131 | Pea15a | 2.207113508 | 1.03676E-07 |
| 1132 | Hist1h4m | 2.201882413 | 0.044650886 |
| 1133 | Slc48a1 | 2.197252119 | 1.92844E-06 |
| 1134 | Hspa8 | 2.193520797 | 3.90872E-09 |
| 1135 | Etv5 | 2.192792913 | 0.002277154 |
| 1136 | Unc93b1 | 2.191975094 | 2.86646E-07 |
| 1137 | Cybb | 2.189188187 | 2.08444E-06 |
| 1138 | Crat | 2.182290525 | 0.005815357 |
| 1139 | Plbd1 | 2.17708559 | 0.00109276 |
| 1140 | Rnh1 | 2.171577129 | 7.75305E-09 |
| 1141 | Nedd4 | 2.16594876 | 0.000311107 |
| 1142 | Evi5 | 2.16547114 | 0.000310891 |
| 1143 | Pcyox1 | 2.163254111 | 0.000605761 |
| 1144 | Gns | 2.159807314 | 5.33297E-07 |
| 1145 | Gusb | 2.156254593 | 1.84313E-09 |
| 1146 | Tubb2a | 2.14725744 | 9.62769E-05 |
| 1147 | Wdr91 | 2.138552933 | 0.000296434 |
| 1148 | Tmem86a | 2.136518452 | 0.004207751 |
| 1149 | Themis2 | 2.132138653 | 0.000449259 |
| 1150 | Pomk | 2.130290768 | 0.000257908 |

| | | | |
|---|---|---|---|
| 1151 | Gcnt1 | 2.126031932 | 0.001222791 |
| 1152 | Gfpt1 | 2.12462775 | 0.001391522 |
| 1153 | Cyfip1 | 2.124451232 | 2.45966E-07 |
| 1154 | Samd8 | 2.122850595 | 0.001473151 |
| 1155 | Zfp991 | 2.120860193 | 0.014943095 |
| 1156 | Arhgef10l | 2.119981027 | 0.038900494 |
| 1157 | Mtm1 | 2.118130617 | 0.005760605 |
| 1158 | Capn5 | 2.116951499 | 0.0257178 |
| 1159 | Tpp1 | 2.114767741 | 1.77625E-09 |
| 1160 | Adssl1 | 2.112173171 | 0.005792109 |
| 1161 | Pigz | 2.111598951 | 0.01996141 |
| 1162 | Mb21d1 | 2.110784318 | 0.027786242 |
| 1163 | Plpp2 | 2.110575641 | 0.028935477 |
| 1164 | Skap2 | 2.108224223 | 4.35777E-05 |
| 1165 | Specc1 | 2.102135278 | 0.001263538 |
| 1166 | Ccnd1 | 2.101100233 | 0.00570128 |
| 1167 | Ppp4r2 | 2.099395906 | 0.000187615 |
| 1168 | Cd38 | 2.098734925 | 4.04645E-07 |
| 1169 | Clcn6 | 2.096270785 | 0.033461313 |
| 1170 | Atp6ap2 | 2.091151524 | 2.62631E-10 |
| 1171 | Osgin1 | 2.084839187 | 0.002771229 |
| 1172 | Nedd4l | 2.080531773 | 0.014354262 |
| 1173 | Casp1 | 2.078809032 | 0.000188837 |
| 1174 | Ddhd1 | 2.076611004 | 0.007468343 |
| 1175 | Adrb2 | 2.074761955 | 5.18297E-05 |
| 1176 | Nr1d1 | 2.07377479 | 0.010948119 |
| 1177 | Ifi30 | 2.070576254 | 1.41971E-06 |
| 1178 | Irf5 | 2.068831544 | 2.64172E-08 |
| 1179 | Slc44a1 | 2.068275697 | 0.015808016 |
| 1180 | Slc38a6 | 2.061133174 | 0.019015625 |
| 1181 | Tpcn2 | 2.060231886 | 0.002603562 |
| 1182 | Igfbp4 | 2.052090005 | 2.22961E-07 |

| | | | |
|---|---|---|---|
| 1183 | Alcam | 2.050466176 | 0.013429178 |
| 1184 | Cd200r4 | 2.050224571 | 0.023193368 |
| 1185 | Insig1 | 2.048925093 | 0.001697642 |
| 1186 | Slc2a1 | 2.048702434 | 0.001247553 |
| 1187 | Gde1 | 2.043716785 | 9.0929E-05 |
| 1188 | Slc38a7 | 2.039544834 | 0.003034274 |
| 1189 | Sbno2 | 2.033090436 | 1.61643E-06 |
| 1190 | Heatr5a | 2.027474829 | 0.010248023 |
| 1191 | Snx9 | 2.027251803 | 1.13759E-05 |
| 1192 | Ralb | 2.026473222 | 3.56375E-05 |
| 1193 | Vwa5a | 2.023180727 | 5.23894E-07 |
| 1194 | Vkorc1 | 2.022214042 | 0.004743925 |
| 1195 | Med7 | 2.021225371 | 0.000798889 |
| 1196 | Zcchc24 | 2.018097386 | 0.020992615 |
| 1197 | Cyp4v3 | 2.017536941 | 0.019997826 |
| 1198 | Il11ra1 | 2.01268299 | 0.03108557 |
| 1199 | Cln5 | 2.002700859 | 0.000109932 |
| 1200 | mt-Tl1 | 2.001898305 | 0.019242241 |
| 1201 | Tbc1d30 | 2.000824646 | 0.017106762 |
| 1202 | Ptprj | 1.997490937 | 0.003600951 |
| 1203 | Rnf157 | 1.993299692 | 0.018975345 |
| 1204 | Gab3 | 1.989548073 | 0.018518916 |
| 1205 | Rnf141 | 1.987496329 | 0.001233098 |
| 1206 | C1qtnf12 | 1.983605161 | 0.013241419 |
| 1207 | Trim30d | 1.981676838 | 0.002868523 |
| 1208 | Il18 | 1.978999482 | 0.027314419 |
| 1209 | Naglu | 1.972007249 | 0.000404403 |
| 1210 | Cln3 | 1.971489302 | 1.25809E-05 |
| 1211 | Slc6a6 | 1.967916337 | 1.12732E-08 |
| 1212 | Fes | 1.9673645 | 0.004786837 |
| 1213 | Gga2 | 1.964343891 | 0.001717804 |
| 1214 | Slc27a1 | 1.962965127 | 0.001707541 |

| | | | |
|---|---|---|---|
| 1215 | Snx29 | 1.96104052 | 0.002951127 |
| 1216 | Slfn5 | 1.957907276 | 0.000109422 |
| 1217 | Fgd2 | 1.957480136 | 0.000506167 |
| 1218 | Rrbp1 | 1.956517706 | 1.99642E-07 |
| 1219 | Idua | 1.950970145 | 5.65493E-05 |
| 1220 | Tm6sf1 | 1.944658449 | 0.003192068 |
| 1221 | Sh3bp2 | 1.943936131 | 0.002416641 |
| 1222 | Lmbrd1 | 1.943601483 | 2.61772E-05 |
| 1223 | Atp6v1b2 | 1.942166678 | 8.36388E-05 |
| 1224 | Ebf1 | 1.939975809 | 0.026879055 |
| 1225 | Gabarapl1 | 1.9396689 | 0.000417969 |
| 1226 | Gna12 | 1.936999459 | 0.012097106 |
| 1227 | Pde4b | 1.93487007 | 3.58397E-06 |
| 1228 | Cflar | 1.934499451 | 2.85507E-06 |
| 1229 | Tspan3 | 1.931785428 | 2.45015E-05 |
| 1230 | Rnasel | 1.928480108 | 6.12195E-05 |
| 1231 | Slc9a9 | 1.92841219 | 1.46374E-05 |
| 1232 | Adap1 | 1.928398966 | 0.029181543 |
| 1233 | Tpd52 | 1.927465747 | 2.04508E-05 |
| 1234 | Vps26a | 1.915719909 | 8.7202E-06 |
| 1235 | Tor1aip2 | 1.915675577 | 3.25592E-06 |
| 1236 | Fez2 | 1.913968261 | 0.011483402 |
| 1237 | Nfkbie | 1.908333465 | 1.82242E-05 |
| 1238 | Tfpi | 1.907781609 | 0.003698555 |
| 1239 | Amz2 | 1.906690964 | 0.002193304 |
| 1240 | AW011738 | 1.906180383 | 0.010850907 |
| 1241 | 2810013P06Rik | 1.904878839 | 0.000463867 |
| 1242 | Rbbp9 | 1.901697553 | 0.025669588 |
| 1243 | Tpi1 | 1.895721975 | 5.27762E-06 |
| 1244 | Etv3 | 1.892668419 | 2.86996E-05 |
| 1245 | 5031439G07Rik | 1.890688236 | 0.002212807 |
| 1246 | Dstn | 1.887683569 | 3.77296E-05 |

| | | | |
|---|---|---|---|
| 1247 | Diaph2 | 1.887444218 | 0.0019878 |
| 1248 | Gbp2 | 1.886278926 | 0.000109501 |
| 1249 | Etfrf1 | 1.883388673 | 0.006701007 |
| 1250 | Igsf8 | 1.879880394 | 0.00040528 |
| 1251 | Spag9 | 1.878786062 | 0.003975818 |
| 1252 | St6galnac4 | 1.877701592 | 7.22768E-05 |
| 1253 | Tnfrsf1b | 1.87151437 | 2.40405E-07 |
| 1254 | Sp140 | 1.870694019 | 0.000221521 |
| 1255 | Atp6v1g1 | 1.868199504 | 2.11358E-05 |
| 1256 | B430306N03Rik | 1.86696978 | 0.042509843 |
| 1257 | Esd | 1.866315858 | 0.000232382 |
| 1258 | Pgam1 | 1.865846386 | 1.16446E-05 |
| 1259 | Rab9 | 1.861256575 | 0.030752634 |
| 1260 | Abhd12 | 1.860124456 | 9.91752E-06 |
| 1261 | Acer3 | 1.857479157 | 0.001321175 |
| 1262 | Cyb5r3 | 1.857034175 | 1.29053E-06 |
| 1263 | Mindy1 | 1.856343956 | 0.00044082 |
| 1264 | Mctp1 | 1.854854525 | 0.001948609 |
| 1265 | Akr1a1 | 1.850425647 | 7.08356E-07 |
| 1266 | Serinc1 | 1.850116866 | 4.26224E-07 |
| 1267 | Axl | 1.846893124 | 0.004494237 |
| 1268 | Pepd | 1.845055355 | 0.001485231 |
| 1269 | Slc2a8 | 1.844897623 | 0.005203433 |
| 1270 | Rufy3 | 1.842648943 | 0.042001936 |
| 1271 | Eif4ebp1 | 1.839832426 | 0.002807098 |
| 1272 | Soat1 | 1.836450343 | 2.04508E-05 |
| 1273 | Gnaq | 1.834954015 | 0.007467798 |
| 1274 | Rnf149 | 1.832095201 | 0.019325146 |
| 1275 | Polg2 | 1.827695464 | 0.028015079 |
| 1276 | Dnajb2 | 1.821445093 | 0.025199293 |
| 1277 | Prkab2 | 1.820072974 | 0.026029221 |
| 1278 | Sde2 | 1.81335091 | 2.68204E-05 |

| | | | |
|---|---|---|---|
| 1279 | Lgals3bp | 1.806020123 | 0.001555739 |
| 1280 | Il4ra | 1.803744065 | 7.64061E-07 |
| 1281 | Nfam1 | 1.803704059 | 0.004818021 |
| 1282 | Pxdc1 | 1.801199358 | 0.046430467 |
| 1283 | Csnk1e | 1.800218028 | 0.009711568 |
| 1284 | Baiap2 | 1.800110563 | 0.001310604 |
| 1285 | Gclm | 1.797771544 | 0.016150294 |
| 1286 | Tep1 | 1.795108856 | 0.000903221 |
| 1287 | Arl2 | 1.789201926 | 0.033859579 |
| 1288 | Alas1 | 1.786858321 | 0.000111822 |
| 1289 | Ccr3 | 1.785966889 | 0.000662015 |
| 1290 | Map2k3 | 1.781636817 | 3.05474E-08 |
| 1291 | Herc6 | 1.780225501 | 0.002563366 |
| 1292 | Pvr | 1.774423163 | 0.000117047 |
| 1293 | Dennd4c | 1.771473313 | 0.019015625 |
| 1294 | Mvp | 1.771094079 | 7.22768E-05 |
| 1295 | Inppl1 | 1.768915328 | 0.043898364 |
| 1296 | P2rx7 | 1.768249743 | 5.69506E-07 |
| 1297 | Hcfc1r1 | 1.754963986 | 0.005753607 |
| 1298 | Rtn4 | 1.752537602 | 0.000368834 |
| 1299 | 5330406M23Rik | 1.749577264 | 0.028003099 |
| 1300 | Ddit3 | 1.748967112 | 0.014246677 |
| 1301 | Arfgef2 | 1.748166478 | 0.007406783 |
| 1302 | Rab5c | 1.743180759 | 0.012354326 |
| 1303 | Plbd2 | 1.742067073 | 0.000144812 |
| 1304 | Il10rb | 1.740819469 | 9.02657E-07 |
| 1305 | Idh2 | 1.739351802 | 0.00184456 |
| 1306 | Nudt9 | 1.738157803 | 0.0004856 |
| 1307 | C920009B18Rik | 1.737104996 | 0.010113446 |
| 1308 | mt-Nd5 | 1.73601161 | 5.99678E-07 |
| 1309 | Noct | 1.730083035 | 0.023209415 |
| 1310 | Npc2 | 1.721495521 | 4.35777E-05 |

| | | | |
|---|---|---|---|
| 1311 | Gabarap | 1.721327445 | 5.72208E-06 |
| 1312 | Kdm6b | 1.71901896 | 0.000113451 |
| 1313 | Cd9 | 1.71759169 | 9.31425E-05 |
| 1314 | Gm28437 | 1.714325785 | 8.75135E-08 |
| 1315 | Prkx | 1.70657794 | 0.001485231 |
| 1316 | Ggta1 | 1.704516235 | 0.018857566 |
| 1317 | Itm2b | 1.704368504 | 8.95907E-06 |
| 1318 | Stx12 | 1.700414418 | 0.005375008 |
| 1319 | Sfmbt1 | 1.697032749 | 0.012555048 |
| 1320 | Snx30 | 1.69240032 | 0.006181788 |
| 1321 | Tor3a | 1.692044126 | 0.003697521 |
| 1322 | Cnn3 | 1.688473499 | 0.011080831 |
| 1323 | Scarb1 | 1.688068281 | 0.000870919 |
| 1324 | Aldh2 | 1.686202191 | 0.000138447 |
| 1325 | Npc1 | 1.683128608 | 1.84084E-05 |
| 1326 | Sec24d | 1.682445116 | 0.006552886 |
| 1327 | Dnase2a | 1.682116926 | 0.001310604 |
| 1328 | Paox | 1.677685535 | 0.026916336 |
| 1329 | Gaa | 1.674446471 | 0.004617402 |
| 1330 | Fads1 | 1.670871325 | 0.01932544 |
| 1331 | Gbp3 | 1.669340629 | 0.001098629 |
| 1332 | St3gal1 | 1.667287149 | 0.001681004 |
| 1333 | Hspa5 | 1.657719006 | 3.57246E-07 |
| 1334 | Fuca1 | 1.65443662 | 0.00010442 |
| 1335 | Serpinb1a | 1.64969895 | 0.0045417 |
| 1336 | Gm12250 | 1.637698318 | 0.007105915 |
| 1337 | Laptm4a | 1.637272204 | 1.04803E-05 |
| 1338 | Acp2 | 1.636196446 | 0.014075777 |
| 1339 | Mfge8 | 1.632485363 | 0.00829693 |
| 1340 | Rreb1 | 1.631186935 | 0.005268137 |
| 1341 | Zfp326 | 1.629097031 | 0.003833699 |
| 1342 | Pdcd1 | 1.628841916 | 0.000192212 |

| | | | |
|---|---|---|---|
| 1343 | D1Ertd622e | 1.628415097 | 0.006008515 |
| 1344 | Ckb | 1.626587586 | 0.024230401 |
| 1345 | Gnpda1 | 1.622725083 | 6.13504E-05 |
| 1346 | Lancl2 | 1.621398955 | 0.049007258 |
| 1347 | Cd200r1 | 1.618388118 | 0.000526227 |
| 1348 | Nfic | 1.612245403 | 0.002110798 |
| 1349 | Camkk2 | 1.607956752 | 0.017206802 |
| 1350 | Arfgap3 | 1.605412256 | 0.027325727 |
| 1351 | Dhx34 | 1.604815985 | 0.039147078 |
| 1352 | Vps37c | 1.598717239 | 0.004912939 |
| 1353 | Sema4a | 1.596872912 | 0.002918957 |
| 1354 | Rab1 | 1.595184849 | 1.50851E-05 |
| 1355 | Ifngr2 | 1.593113572 | 0.003446447 |
| 1356 | Snx2 | 1.591805895 | 0.00021236 |
| 1357 | Arl13b | 1.586619869 | 0.029978636 |
| 1358 | Mul1 | 1.584044537 | 0.028608861 |
| 1359 | Hexb | 1.580856853 | 0.004322333 |
| 1360 | Tax1bp3 | 1.580153126 | 0.028786567 |
| 1361 | Grina | 1.579036069 | 1.0399E-05 |
| 1362 | Rab11a | 1.578674752 | 0.000642445 |
| 1363 | Hspe1 | 1.573847248 | 0.001413845 |
| 1364 | R3hdm2 | 1.570742174 | 0.024053536 |
| 1365 | Oas3 | 1.5704204 | 0.028003099 |
| 1366 | Parp14 | 1.56908136 | 0.004634153 |
| 1367 | Mafk | 1.56892312 | 0.007392511 |
| 1368 | Gclc | 1.568059959 | 0.024893752 |
| 1369 | Pofut2 | 1.566707568 | 0.013085551 |
| 1370 | Atp6v1h | 1.557372478 | 0.000745114 |
| 1371 | Tmem109 | 1.556374452 | 0.009221187 |
| 1372 | Pde1b | 1.554427577 | 0.030604998 |
| 1373 | Socs6 | 1.552104848 | 0.011084982 |
| 1374 | Mapkbp1 | 1.551263221 | 0.039667299 |

| | | | |
|---|---|---|---|
| 1375 | Ctsh | 1.551132291 | 0.002408227 |
| 1376 | H2-DMb1 | 1.550563074 | 0.03738061 |
| 1377 | Synj1 | 1.543799516 | 0.008543753 |
| 1378 | Tmem5 | 1.541333919 | 0.00570128 |
| 1379 | Ndel1 | 1.540085476 | 0.000966464 |
| 1380 | Twf1 | 1.537685956 | 0.002857004 |
| 1381 | Mfsd6 | 1.53553981 | 0.019601993 |
| 1382 | Samd9l | 1.534831177 | 0.001801423 |
| 1383 | Rheb | 1.533573807 | 0.000139484 |
| 1384 | Zcchc2 | 1.529575943 | 0.014768561 |
| 1385 | Nrros | 1.525617818 | 0.000576706 |
| 1386 | Por | 1.524050162 | 0.007161964 |
| 1387 | Flcn | 1.521023778 | 0.000101024 |
| 1388 | Dcxr | 1.520709452 | 0.029901375 |
| 1389 | Fam76a | 1.520032497 | 0.032193277 |
| 1390 | Socs4 | 1.519662261 | 0.045975294 |
| 1391 | Pqlc2 | 1.518406373 | 0.031518846 |
| 1392 | Sh3bp5 | 1.518269059 | 0.000203775 |
| 1393 | Ppfia1 | 1.515954378 | 0.032281336 |
| 1394 | Fam126a | 1.515739609 | 0.002498431 |
| 1395 | Smcr8 | 1.511991463 | 0.022413936 |
| 1396 | Txndc16 | 1.505881905 | 0.008738936 |
| 1397 | Atp6v1c1 | 1.504755463 | 0.001919358 |
| 1398 | Gpr65 | 1.501398566 | 0.001543484 |
| 1399 | Dnaja1 | 1.49427113 | 1.13126E-06 |
| 1400 | Tmem189 | 1.493594006 | 0.013074036 |
| 1401 | Katnbl1 | 1.490747977 | 0.024124115 |
| 1402 | Tmem87b | 1.489651859 | 0.049619414 |
| 1403 | Dennd5a | 1.487923801 | 0.006051578 |
| 1404 | Sat1 | 1.485160317 | 3.64457E-05 |
| 1405 | Nbdy | 1.479056839 | 0.039393177 |
| 1406 | Rnf13 | 1.477517086 | 0.002053233 |

| | | | |
|---|---|---|---|
| 1407 | Sh3bgrl | 1.475671561 | 2.07437E-05 |
| 1408 | Tmem242 | 1.474475227 | 0.044928266 |
| 1409 | Lamtor3 | 1.472669897 | 0.001344551 |
| 1410 | Maf | 1.471960385 | 0.000761565 |
| 1411 | Arid5b | 1.471648667 | 0.000831186 |
| 1412 | Vps18 | 1.46704491 | 0.023912488 |
| 1413 | Camta2 | 1.466811625 | 0.01635012 |
| 1414 | Oser1 | 1.463339532 | 0.019568049 |
| 1415 | Ptpn23 | 1.460980569 | 0.016579981 |
| 1416 | Arhgef7 | 1.460862691 | 0.025642898 |
| 1417 | Aff1 | 1.460239115 | 0.000665371 |
| 1418 | Atp6ap1 | 1.447029217 | 4.6632E-05 |
| 1419 | Map4k4 | 1.441803666 | 0.030820499 |
| 1420 | Baz2b | 1.437615516 | 0.004380156 |
| 1421 | Ccdc86 | 1.437141034 | 0.036495474 |
| 1422 | Osbpl8 | 1.43502055 | 0.004141524 |
| 1423 | Gng2 | 1.429544444 | 0.000635366 |
| 1424 | Degs1 | 1.429473106 | 0.000180467 |
| 1425 | Plekhm1 | 1.423250171 | 0.027225261 |
| 1426 | Mpp1 | 1.421062069 | 0.000798971 |
| 1427 | Irf7 | 1.418645054 | 0.040115662 |
| 1428 | Fuca2 | 1.418585335 | 0.006106993 |
| 1429 | mt-Nd2 | 1.415839713 | 0.000455632 |
| 1430 | Slc43a2 | 1.415134906 | 0.002720146 |
| 1431 | Adnp2 | 1.414773238 | 0.02732173 |
| 1432 | Sh3glb1 | 1.414746536 | 0.000245959 |
| 1433 | Dennd1a | 1.412956578 | 0.01088059 |
| 1434 | Rsu1 | 1.403394448 | 0.006649308 |
| 1435 | Irf1 | 1.401994588 | 0.000441424 |
| 1436 | Lamtor1 | 1.400470836 | 0.00077185 |
| 1437 | Psap | 1.399184667 | 0.000115372 |
| 1438 | Lrp6 | 1.396920964 | 0.04097282 |

| | | | |
|---|---|---|---|
| 1439 | Nfkbid | 1.395635309 | 0.000169598 |
| 1440 | Arpc3 | 1.392580942 | 0.00333004 |
| 1441 | Snx6 | 1.390899285 | 0.002212159 |
| 1442 | Prcp | 1.389587892 | 0.03173135 |
| 1443 | Fcho2 | 1.389097123 | 0.002083788 |
| 1444 | Tmem256 | 1.386834581 | 0.006171619 |
| 1445 | Tagln2 | 1.379031346 | 0.000686048 |
| 1446 | Plin3 | 1.376449361 | 0.002508766 |
| 1447 | Birc3 | 1.373270151 | 3.09117E-05 |
| 1448 | Stard4 | 1.371203554 | 0.025144818 |
| 1449 | Stam2 | 1.370866245 | 0.011317835 |
| 1450 | Ostf1 | 1.370348318 | 0.000442476 |
| 1451 | Fam110a | 1.368891838 | 0.019954632 |
| 1452 | Lrrfip2 | 1.363925448 | 0.023110213 |
| 1453 | Eea1 | 1.361133057 | 0.005203133 |
| 1454 | mt-Co3 | 1.357707096 | 0.000167125 |
| 1455 | Ecm1 | 1.356534428 | 0.005189611 |
| 1456 | Phyh | 1.35295782 | 0.032022848 |
| 1457 | Cmip | 1.350889771 | 0.047924192 |
| 1458 | Alkbh5 | 1.343573269 | 0.040645808 |
| 1459 | mt-Atp6 | 1.343398876 | 0.000601671 |
| 1460 | Reps1 | 1.342487799 | 0.013253168 |
| 1461 | Slc29a3 | 1.334795939 | 0.007911656 |
| 1462 | Selenof | 1.333375679 | 0.0048346 |
| 1463 | Rilpl2 | 1.325678198 | 0.003406091 |
| 1464 | Nfkbib | 1.317620258 | 0.018141956 |
| 1465 | Galnt1 | 1.31654441 | 0.021039752 |
| 1466 | Atox1 | 1.313685322 | 0.006241189 |
| 1467 | Tmbim4 | 1.309006694 | 0.006128401 |
| 1468 | Pld4 | 1.306568656 | 0.007450444 |
| 1469 | Fam50a | 1.304819585 | 0.027789267 |
| 1470 | Tbc1d5 | 1.300015291 | 0.043999088 |

| | | | |
|---|---|---|---|
| 1471 | Chd7 | 1.299554492 | 0.004401993 |
| 1472 | Snx3 | 1.296978818 | 0.004232719 |
| 1473 | S100a11 | 1.293276081 | 0.003207643 |
| 1474 | Gm28875 | 1.291243173 | 0.033312931 |
| 1475 | Rnf2 | 1.283446963 | 0.027906696 |
| 1476 | Tpm1 | 1.281852855 | 0.007656761 |
| 1477 | Cnih4 | 1.278121405 | 0.029155104 |
| 1478 | Gpcpd1 | 1.272923445 | 0.001044893 |
| 1479 | Slc38a1 | 1.262622981 | 0.005948866 |
| 1480 | Stard3nl | 1.257781377 | 0.022219377 |
| 1481 | 1700017B05Rik | 1.256199591 | 0.001854944 |
| 1482 | Slc20a1 | 1.25353707 | 0.017887693 |
| 1483 | Slk | 1.252117927 | 0.012315596 |
| 1484 | Irgm1 | 1.251230146 | 0.012966946 |
| 1485 | Purb | 1.248509822 | 0.022189395 |
| 1486 | Snap23 | 1.246179894 | 0.021528542 |
| 1487 | Pttg1ip | 1.238775842 | 0.00749554 |
| 1488 | Dram2 | 1.235756785 | 0.048690248 |
| 1489 | Cox17 | 1.234437003 | 0.02246644 |
| 1490 | Arl8b | 1.223021062 | 0.001247704 |
| 1491 | Higd1a | 1.217497006 | 0.016893791 |
| 1492 | Evi2a | 1.216758741 | 0.026029221 |
| 1493 | Eif4e | 1.210283865 | 0.001037045 |
| 1494 | Selenok | 1.202098177 | 0.002661711 |
| 1495 | Nr3c1 | 1.200888456 | 0.002732741 |
| 1496 | Pgk1 | 1.200225575 | 0.001983108 |
| 1497 | Nptn | 1.196497082 | 0.002277154 |
| 1498 | Myo1c | 1.194506965 | 0.016085708 |
| 1499 | Reep3 | 1.189230924 | 0.027362498 |
| 1500 | Arap1 | 1.188589311 | 0.009249851 |
| 1501 | Atraid | 1.183362398 | 0.022746683 |
| 1502 | mt-Nd6 | 1.18235922 | 0.003041083 |

| | | | |
|---|---|---|---|
| 1503 | mt-Co2 | 1.179244162 | 0.016034899 |
| 1504 | Gm10925 | 1.179232288 | 0.01230544 |
| 1505 | Ubb | 1.174558255 | 0.001325668 |
| 1506 | Mylip | 1.174377252 | 0.025583296 |
| 1507 | Tab2 | 1.173371617 | 0.030243602 |
| 1508 | Atf4 | 1.173158583 | 0.047482737 |
| 1509 | Hnrnph2 | 1.172157213 | 0.019160366 |
| 1510 | Gm28661 | 1.168419972 | 0.020293893 |
| 1511 | Prdx6 | 1.165003949 | 0.003192068 |
| 1512 | Phf20 | 1.161816066 | 0.026516023 |
| 1513 | Dnajb6 | 1.151339917 | 0.011327723 |
| 1514 | Pld3 | 1.148857747 | 0.002067425 |
| 1515 | Hsp90ab1 | 1.147928486 | 0.002495891 |
| 1516 | Ube2l6 | 1.147629283 | 0.022829909 |
| 1517 | Pdia6 | 1.147249638 | 0.010675262 |
| 1518 | Stx7 | 1.14231404 | 0.049084986 |
| 1519 | Eif1a | 1.137942934 | 0.041208456 |
| 1520 | Gm1821 | 1.131695965 | 0.028929747 |
| 1521 | Hsd17b12 | 1.129668635 | 0.02802397 |
| 1522 | Phactr2 | 1.122057375 | 0.038848152 |
| 1523 | Atl3 | 1.118225371 | 0.046298304 |
| 1524 | mt-Nd1 | 1.117628079 | 0.000515771 |
| 1525 | Tmed10 | 1.112281895 | 0.015907956 |
| 1526 | Plcxd2 | 1.1089119 | 0.021989553 |
| 1527 | Atpif1 | 1.104907437 | 0.047815846 |
| 1528 | Man2b1 | 1.097852283 | 0.016771028 |
| 1529 | Rab7 | 1.097826985 | 0.007286581 |
| 1530 | P4hb | 1.087183403 | 0.007026919 |
| 1531 | Pdlim5 | 1.078524665 | 0.038004374 |
| 1532 | Txn1 | 1.07446015 | 0.03275395 |
| 1533 | Sirt2 | 1.074066477 | 0.038546023 |
| 1534 | Tbc1d15 | 1.068243267 | 0.021416981 |

| | | | |
|---|---|---|---|
| 1535 | Ccnl1 | 1.066192119 | 0.004415902 |
| 1536 | Gpr183 | 1.064340352 | 0.015578951 |
| 1537 | Nfkb1 | 1.062589915 | 0.023699791 |
| 1538 | Rbfa | 1.061263506 | 0.036050715 |
| 1539 | Vamp8 | 1.060611819 | 0.027868422 |
| 1540 | Serinc3 | 1.054562252 | 0.001903608 |
| 1541 | Atf1 | 1.053784161 | 0.019043664 |
| 1542 | Nras | 1.052372935 | 0.003566593 |
| 1543 | Mapre2 | 1.047945366 | 0.015963343 |
| 1544 | Pgd | 1.043582664 | 0.01892556 |
| 1545 | Scamp2 | 1.042921278 | 0.04758186 |
| 1546 | Relb | 1.041698958 | 0.038500689 |
| 1547 | Rnf125 | 1.039220874 | 0.021507748 |
| 1548 | Arf4 | 1.036204479 | 0.007469137 |
| 1549 | Map2k1 | 1.033801306 | 0.025144818 |
| 1550 | Trim26 | 1.031403381 | 0.013971346 |
| 1551 | Napa | 1.026778052 | 0.014693298 |
| 1552 | Samsn1 | 1.02344481 | 0.013241419 |
| 1553 | Capza2 | 1.021552599 | 0.026660103 |
| 1554 | 2610001J05Rik | 1.016642957 | 0.045583792 |
| 1555 | Osbpl9 | 1.01314117 | 0.004156986 |
| 1556 | 2010111I01Rik | 1.011650518 | 0.042396332 |
| 1557 | Lipa | 1.009093881 | 0.035698244 |
| 1558 | Hdlbp | 1.008704593 | 0.017040343 |
| 1559 | Ywhag | 1.006549329 | 0.046495871 |
| 1560 | Capg | 1.000286218 | 0.025585424 |
